# Dishevelled2 Condensates control Axon Initial Segment (AIS) assembly and function through MTCL1-Dependent microtubule recruitment

**DOI:** 10.64898/2026.10.08.757526

**Authors:** Sophie Belmonte, Sonia Poirault-Chassac, Juliette Villagomez, Charlotte Isch, Ana Dorrego-Rivas, Margot Donier, Emilie Chagny, Ellyn Renou, Louis Vacher, Nathalie Aubailly, Maïté Marie Moreau, Melissa David, Magali Mondin, Mireille Montcouquiol, Matthieu Sainlos, Arne Battefeld, Nathalie Sans, Jerome Ezan

**Author notes:** Equal contribution (Sophie Belmonte, Sonia Poirault-Chassac). Equal Senior contribution (Nathalie Sans, Jerome Ezan). This work is dedicated to the memory of our esteemed colleague, friend, partner, mentor, Mireille Montcouquiol who recently passed away.

## Abstract

In the epithelium, Planar Cell Polarity (PCP) signaling coordinates cellular organization along the plane of the tissue, perpendicular to the apicobasal axis. In neurons, polarity is instead expressed through the structural and molecular asymmetry that separates the axon from the somatodendritic compartment. Core PCP proteins are essential for neuronal development and function, and have been linked to neurodevelopmental disorders, including autism spectrum disorders and epilepsy. However, their precise roles in the establishment of neuronal polarity remain incompletely understood.

Here, we identify Dishevelled2 (Dvl2), a central PCP signaling scaffold, as a novel component of the axon initial segment (AIS), the proximal axonal domain that maintains neuronal polarity and initiates action potentials. We show that Dvl2 defines, both *in vivo* and in cultured hippocampal neurons, a proximal subdomain positioned upstream of the major AIS marker Ankyrin-G (Ank-G) and is closely associated with microtubule bundles. Dvl2 is already detected during early axon specification, suggesting that it participates in the initial organization of the proximal axon. Functional depletion of Dvl proteins disrupts AIS architecture, reduces the accumulation of Ankyrin-G and other core AIS components, and impairs the organization of AIS microtubule bundles. These structural defects translate into altered neuronal excitability, with a reduced AIS-dependent component of the action potential.

Mechanistically, we identify MTCL1 as a microtubule-associated partner that recruits Dvl2 from biomolecular condensate assemblies onto microtubules. This recruitment requires the DIX and PDZ domains of Dvl2. Notably, selective intracellular binders targeting the Dvl2 PDZ domain perturb Dvl2 condensate organization and compromise AIS formation.

Together, our data identify Dvl2 as an organizer of the proximal AIS and reveal a Dvl2-MTCL1-dependent mechanism coupling condensate scaffolding, microtubule organization, and neuronal excitability.

**HIGHLIGHTS:**

- Dvl2 is an early component of the proximal axon initial segment (AIS)
- Dvl2 associates with AIS microtubule bundles and depends on MTCL1 for microtubule recruitment
- Dvl depletion disrupts AIS cytoskeletal organization, AnkG accumulation, sodium channel clustering and action potential initiation
- Perturbing Dvl2 condensate organization impairs AIS integrity

## INTRODUCTION

Neurons are highly polarized cells whose specialized compartments receive, process, and transmit electrical signals. This polarity is established during axon specification and maintained throughout neuronal life^1,2^. A major landmark of neuronal polarity is the axon initial segment (AIS), a specialized proximal axonal domain that is enriched in Ankyrin-G (AnkG), cell adhesion molecules, and highly organized cytoskeletal networks^3–6^. It links neuronal polarity and excitability by concentrating voltage-gated sodium and potassium channels within a domain that initiates action potentials^7–9^. Its disruption has been linked to neurodevelopmental disorders, like autism, intellectual disability, and epilepsy, as well as neurodegenerative and psychiatric diseases^10–13^.

During development, AIS assembly is coordinated with axonal specification through recruitment of specialized protein complexes to the proximal axon and cytoskeletal organization^14^. Early AIS assembly relies on a crosslinked microtubule (MT) scaffold^15^, combined with nanoscale actin organization that anchors and stabilizes key AIS proteins^16,17^. The AIS also preserves neuronal polarity by acting as a selective diffusion and trafficking barrier that maintains the molecular identity of the axonal compartment^6,18,19^. The AIS is organized around a dense macromolecular scaffold built by AnkG, the master organizer of the AIS, together with βIV-spectrin and multiple membrane and cytoskeletal partners^20–22^. In addition, a specialized proximal MT architecture precedes and stabilizes AIS maturation, including the MT-associated proteins tripartite motif-containing 46 (TRIM46) and Microtubule Cross-Linking factor 1 (MTCL1), which partially overlap with the Ankyrin-G-positive domain^23–25^. TRIM46 organizes parallel MT arrays, whereas MTCL1 crosslinks MTs to maintain AIS structure, but how they connect MTs to AIS scaffolding and polarity signaling remains unclear.

Planar Cell Polarity (PCP) signaling provides an attractive framework to understand how local cytoskeletal organization can be coupled to neuronal compartmentalization. In epithelial tissues, PCP signaling coordinates the orientation of cells within the plane of the tissue, perpendicular to the apicobasal axis^26–29^. The core PCP pathway comprises Frizzled, Vangl, and Celsr at the membrane, together with cytoplasmic Prickle, Dishevelled, and Diversin^30–32^, which coordinate adhesion, cytoskeletal dynamics, and polarized cell behavior ^33–36^. Although neurons do not display an obvious planar reference axis comparable to epithelial sheets, PCP proteins are widely expressed in the nervous system and have been implicated in axonal outgrowth and axon guidance ^37–39^, synaptic function ^40,41^, myelination^42^, neuromuscular junction formation^43^ and hippocampal-dependent cognitive processes ^44,45^. Mutations in PCP genes are associated with neurodevelopmental disorders, including neural tube defects, autism spectrum disorders, intellectual disability and epilepsy^46–51^, yet the molecular bases of these pathologies remain poorly understood. Recently, we identified Prickle2 as an AIS-associated neuronal polarity determinant that promotes early AIS assembly by binding Ankyrin-G and regulating MT bundling^52^. Other PCP-associated proteins also contribute to AIS organization: Daam1 remodels longitudinal actin fibers during AIS plasticity^53^, whereas Scrib organizes membrane-associated ion-channel complexes^54^. Collectively, these findings establish PCP proteins as compartment-specific axonal regulators and suggest that PCP-linked disorders may involve dysfunction across neuronal domains, including synapses and the AIS. Identifying additional AIS-associated PCP components would further define their role in organizing the proximal axonal cytoskeleton and molecular architecture.

The three mammalian isoform of Dishevelled (Dvl) proteins are central cytosolic scaffolds shared by canonical Wnt/β-catenin and non-canonical Wnt/PCP signaling^55–57^. In canonical signaling, they promote β-catenin stabilization and transcriptional activation, whereas in PCP pathways they more directly regulate cytoskeletal remodeling, adhesion, and polarized cell behavior^58,59^. In the nervous system, Wnt/Dvl signaling contributes to synaptic plasticity and memory^60,61^, and its deregulation has been linked to autism^62^ and Alzheimer’s disease^63^. Dishevelled proteins are composed of conserved DIX, PDZ, and DEP domains^64^ that support multivalent interactions, oligomerization, membrane recruitment, and signalosome assembly^65–67^. Importantly, Dishevelled proteins form dynamic condensate-like assemblies that concentrate signaling components and regulate pathway output through biomolecular condensation and liquid–liquid phase separation^68–71^. In neurons, biomolecular condensates have emerged as important organizers of synaptic compartments, RNA granules, cytoskeletal regulators, and disease-linked protein assemblies^72,73^. Previous studies linked Dishevelled to neuronal microtubule regulation, with Dvl1 promoting GSK3β-dependent stabilization of MAP1B and MAP2 and Dvl supporting axon formation through atypical PKC-dependent microtubule assembly^74–76^. However, whether specific Dvl isoforms regulate AIS assembly, proximal microtubules, or excitability remains unclear, and their compartment-specific functions in mature neurons are poorly defined^77^.

Here, we identify Dvl2 as a proximal AIS component associated with microtubules from early axon specification through maturation. Dvl depletion disrupts AIS architecture, reduces key scaffold and ion-channel AIS components, impairs microtubule bundling, and alters action potential initiation. Mechanistically, MTCL1 recruits Dvl2 condensate-like assemblies to microtubules through the DIX and PDZ domains, while PDZ-directed binders perturb Dvl2 organization and Ankyrin-G integrity. Together, these findings define a Dvl2–MTCL1 mechanism linking condensate scaffolding, microtubule organization, AIS assembly, and neuronal excitability. They further suggest that condensate-based regulation of cytoskeletal scaffolds may represent a previously unrecognized principle of AIS assembly, neuronal polarity and excitability.

## RESULTS

### Dvl2 is specifically enriched in the proximal AIS, both *in vivo* and *in vitro*, relative to Ankyrin-G. Fig. 1/S1-S4

Building on our hypothesis that PCP proteins can act as *bona fide* AIS regulators, we examined the localization of additional core PCP components at the AIS, with a particular focus on Dvl2. To examine the *in vivo* distribution and localization of Dvl2 in the adult forebrain, we performed immunofluorescence labeling on rat and mouse brain sections using a well-characterized Dvl2-specific antibody^78,79^. Our results demonstrated that Dvl2 is significantly enriched within the AIS, defined by AnkG, across the cortex and several hippocampal regions, including CA1 and the dentate gyrus (DG), both during early postnatal development (P7) and in juvenile animals (P21–25) **(Fig. 1A-B and Fig. S1A-C)**. Notably, the Dvl2 localization pattern appeared particularly pronounced in the most proximal region of the AIS. To further define Dvl2 localization *in vitro*, we generated spatiotemporal profiles of normalized fluorescence intensity in rat hippocampal neurons at DIV3–4, during early AIS formation, and at DIV7–10, when the AIS is structurally mature. Although Dvl2 is present throughout the somatodendritic compartment, it is markedly enriched at the AIS, where its axonal labeling only partially overlaps with AnkG and is preferentially concentrated within the proximal AIS domain (**Fig. 1C; Fig. S2)**. Quantitative analyses confirmed a proximal enrichment of Dvl2 along the AIS, with a fluorescence peak located just upstream of AnkG and progressively decreasing distally **(Fig. 1D)**. While Dvl2 and AnkG profiles largely overlapped at DIV3, the Dvl2 peak progressively shifted proximally from DIV4 onward and remained displaced through DIV10 **(Fig. S2A– C)**. Furthermore, our data revealed that Dvl2 is also enriched at the AIS of axon-carrying dendrite (AcD) neurons **(Fig. S2D)**, which share a similar cytoskeletal architecture with soma-derived AISs^80,81^. At DIV14, Dvl2 staining overlapped with TRIM46^23^ in the proximal AIS, whereas only partial overlap was observed with AnkG **(Fig. 1E)**. To further validate the Dvl2 antibody, we mapped its epitope to amino acids 600–614 of mouse Dvl2, a region divergent among Dvl paralogues but highly conserved across mouse, human, and rat Dvl2, supporting both isoform specificity and cross-species use **(Fig. S3A)**. By contrast, the Dvl3 antibody^82^ showed predominantly diffuse somatic labeling, with no apparent AIS enrichment **(Fig. S3B-C)**. Most importantly, we independently confirmed the enrichment of Dvl2 at the AIS using a second Dvl2 antibody^83,84^, excluding an antibody-specific staining artifact **(Fig. S3A,D)**. Finally, western blotting and immunofluorescence in COS-7 cells expressing HA-Dvl1, HA-Dvl2, or HA-Dvl3 confirmed that each antibody selectively recognized its corresponding isoform without detectable cross-reactivity (**Fig. S4)**. Altogether, these findings suggest that Dvl2 delineates a subdomain positioned between the soma and the proximal boundary of the AIS, exhibiting a labeling pattern reminiscent of that of TRIM46 and MTCL1 that are early markers of AIS formation known to stabilize microtubules the stabilization of microtubules (MTs)^23,25^.

**Figure 1.**
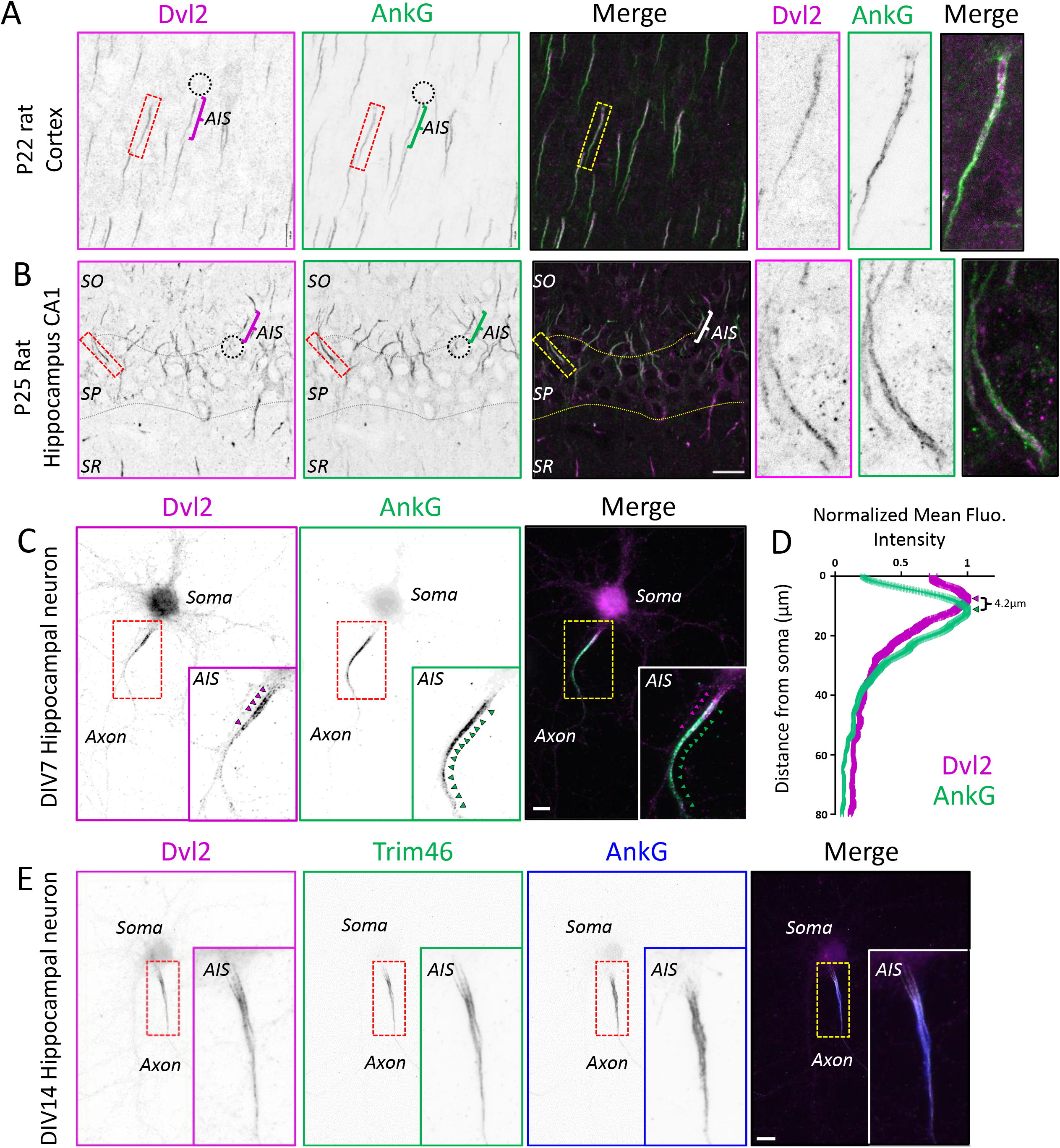
Dvl2 is enriched at the AIS *in vivo* and *in vitro.* (**A-B**) Illustrations of coronal section of brains immunolabeled for Dvl2 (magenta) and AnkG (green) in the rat cortex at P22 (A) and in the CA1 region of the rat hippocampus at P25 (B). Dvl2 is enriched in the proximal part of the AIS (bracket, see high magnifications). Dotted circles outline the cell body and brackets the AIS. (**C**) DIV7 hippocampal neurons in vitro immunolabeled for Dvl2 (magenta), AnkG (green). Dvl2 is strongly enriched at the proximal part of the AIS (see inset, magenta arrowheads) and colocalizes partially with AnkG (bracket and see inset, green arrowheads). (**D**) Linescans of normalized fluorescence intensity at the AIS of Dvl2 (magenta) and AnkG (green) at DIV7. Note the upstream, more proximal, shift of the fluorescence Dvl2 intensity peak compared to AnkG. Values are normalized to the maximum intensity of each marker and displayed with ± SEM. (**E**) DIV14 hippocampal neurons in vitro immunolabeled for Dvl2 (magenta), TRIM46 (green) and AnkG (blue). Dvl2 staining overlaps with TRIM46 immunolabeling at the proximal AIS and but only partly with AnkG. Scale bars: 20 µm in A, 10 µm in B, C and E.

### Dvl2 is an early *bona fide* MT-associated component of the AIS. Fig. 2

To delineate the subcellular distribution of Dvl2 during the earliest phases of neuronal polarization and AIS assembly, we examined rat hippocampal neurons at DIV1, corresponding to stage 3 and the onset of axonal specification^85^. At this stage, Dvl2 and AnkG appeared as discrete aggregates exhibiting a discontinuous distribution along the nascent axon **(Fig. 2A)**. Furthermore, endogenous Dvl2 accumulated within axons and linescan analysis confirmed a partial overlap with TRIM46 immunolabeling **(Fig. 2B)**. The presence of Dvl2 in neurons at the stage of axonal specification supports the hypothesis that Dvl2 functions in the early determination of neuronal polarity, possibly by promoting microtubule stabilization. Because Dvl2 exhibited an early and spatially restricted enrichment in developing axons, we next sought to resolve its precise localization within polarized, more mature neurons using high-resolution imaging approaches. To further investigate the precise localization of Dvl2 in the AIS, we first employed super-resolution ultrastructure expansion microscopy (U-ExM)^86^. This approach revealed Dvl2 enrichment along TRIM46-positive MT bundles within the axonal shaft, suggesting a close spatial association with the cytoskeletal network **(Fig. 2C)**. We further validated this association using single-molecule localization microscopy (STORM), which revealed extensive colocalization of Dvl2 with stable microtubules. Coordinate-based analysis yielded a Manders overlap coefficient of 0.884 for Dvl2 relative to tubulin, confirming a high degree of spatial overlap at nanoscale resolution (**Fig. 2D)**. At the nanoscale, Dvl2 formed discrete puncta aligned along microtubule tracks, but this spatial overlap does not necessarily reflect a direct physical interaction; rather, Dvl2 may associate with microtubules indirectly via linker proteins such as MTCL1 or TRIM46. Such an organization suggests that Dvl2 may contribute to the establishment or maintenance of the AIS, where microtubule organization is critical for neuronal polarity^24^.

**Fig. 2:**
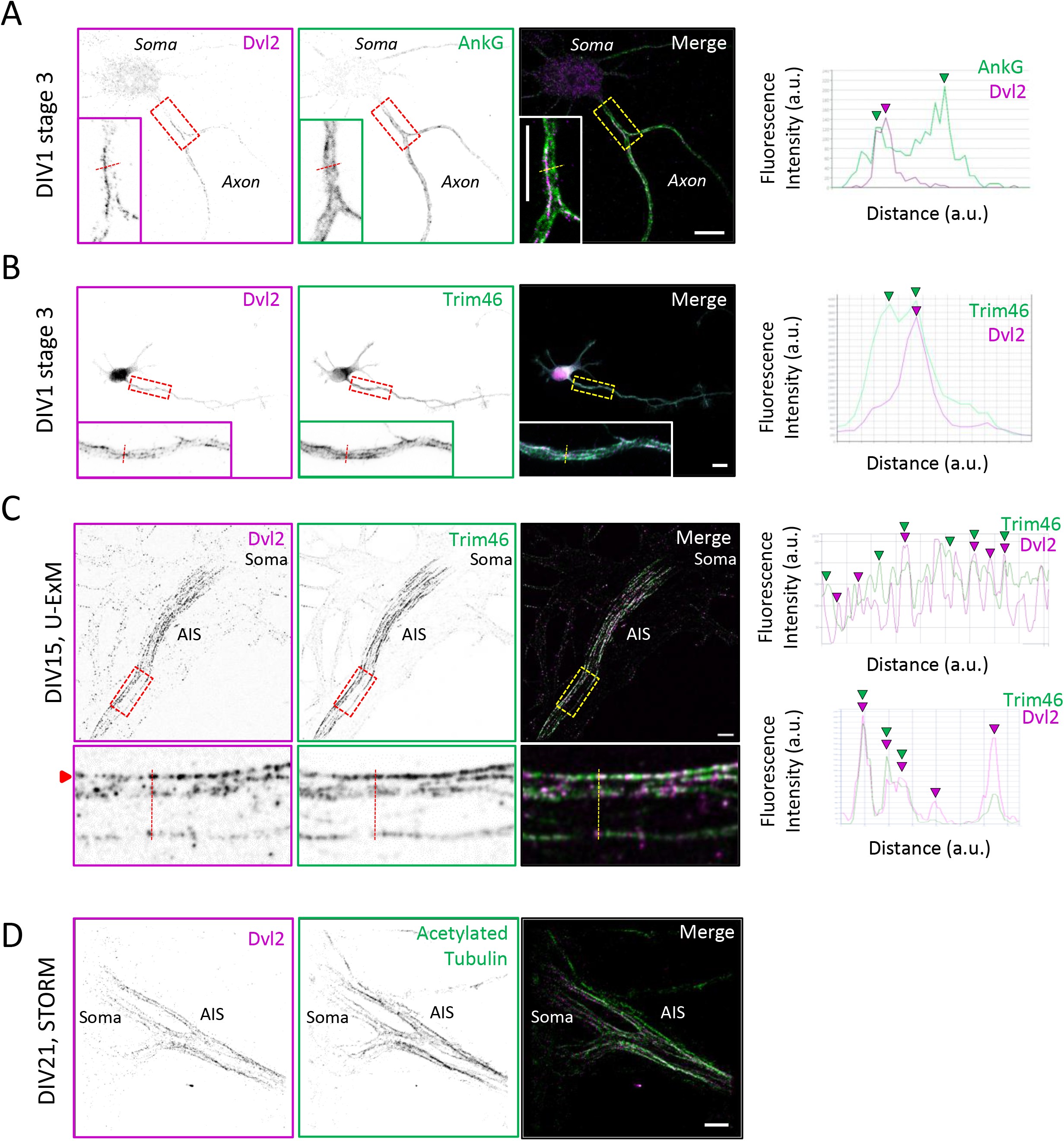
Dvl2 is an early MT-associated *bona fide* component of the AIS. (**A**) Illustration of a stage 3 neuron at DIV1 with Dvl2 (green) and AnkG (magenta) labelling imaged by confocal microscopy. Dvl2 is detected in the specified, nascent axon (see high magnification) with a partial colocalization with AnkG as illustrated on the linescan. **(B**) Illustration of a stage 3 neuron at DIV1 with Dvl2 (magenta) and TRIM46 (green) labelling imaged by conventional microscopy. Note that Dvl2 colocalize only partly in clusters (see high magnification) with TRIM46 as illustrated on the linescan. (**C**) Ultrastructure expansion microscopy (U-ExM) reveals TRIM46+ microtubules (green) decorated with Dvl2 (magenta) in DIV15 hippocampal neurons. (**D**) Representative image of a dSTORM reconstruction of DIV21 hippocampal neurons stained for Dvl2 (magenta) and acetylated tubulin (green). Scale bars: 10 µm in A and B, 1 µm in C, 2 µm in D.

### Dvl proteins are required for AIS assembly, maintenance, cytoskeletal organization, and sodium channel clustering Fig. 3, S5-S6

Because establishment of the AIS depends on the coordinated recruitment of scaffolding, membrane, and cytoskeletal components, we examined whether early depletion of Dvl affected AIS formation in developing neurons. In DIV7 control neurons, Dvl2 and AnkG delineated a single, continuous AIS in the proximal axon, whereas shDvl nucleofection efficiently reduced Dvl2 levels, confirming the efficacy of the previously validated construct^76^ **(Fig. 3A-D).** In shDvl-nucleofected neurons, we observed a significant increase in the proportion of neurons displaying abnormal AIS morphologies, including a thinner AIS, fragmented AIS labeling, or even a complete loss of detectable AnkG signal **(Fig. 3B-D)**. Quantification confirmed a strong increase in the proportion of neurons with a thin AIS after Dvl depletion , together with a marked redistribution of AIS phenotypes from predominantly normal in controls toward fragmented (2–3 fragments) (47.5 ± 3.8% vs 21.9 ± 2.4% in Ctrl neurons), very fragmented (≥4 fragments) (27.8 ± 4.1% vs 0.9 ± 0.9% in Ctrl neurons), or absent AIS structures in shDvl neurons (7.8 ± 1.7% vs none in Ctrl neurons) **(Fig. 3E-F)**. These observations indicate that Dvl2 is required for proper neuronal polarity but also AIS assembly and structural integrity. Consistent with these structural defects, depletion of Dvl2 reduced the accumulation of key AIS components. Quantification of fluorescence intensity showed that both AnkG and Dvl2 levels were significantly decreased in shDvl neurons by 42% and 51% respectively [intensity levels were 2.4 ± 0.1 × 10^3^ a.u. arbitrary units (a.u.) for AnkG and 6.2 ± 0.1 × 10^2^ a.u. for Dvl2 compared to the control values of 4.2 ± 0.1 × 10^3^ a.u. for AnkG and 12.7 ± 0.1 × 10^2^ a.u. for Dvl2], excluding cells with complete AIS loss **(Fig. 3G-H)**. Other AIS-associated markers, including TRIM46^23^, Pk2^87^ and Scrib^54^, were also significantly reduced upon Dvl depletion. In shDvl neurons, TRIM46, Pk2 and Scrib fluorescence intensities decreased by approximately 57%, 29% and 52%, respectively, reaching 1.6 ± 0.1 × 10³ a.u., 6.3 ± 0.5 × 10³ a.u. and 4.8 ± 0.4 × 10³ a.u., compared with 3.6 ± 0.2 × 10³ a.u., 8.9 ± 0.5 × 10³ a.u. and 9.9 ± 0.6 × 10³ a.u. in control neurons **(Fig. S5)**. Also, the residual AISs in shDvl neurons were significantly longer than in controls (49.0 ± 1.7 µm vs 33.7 ± 0.9 µm) **(Fig. 3I)** and were shifted distally, with an increased proportion of AISs starting >7 µm from the soma **(Fig. 3J-K)**. This suggests that Dvl depletion does not simply abolish AIS formation, but instead disrupts its proper organization and compaction. Considering AIS architecture is tightly linked to the underlying microtubule cytoskeleton, we next assessed the organization of TRIM46-positive microtubule bundles using ultrastructure expansion microscopy^86^. In control neurons, TRIM46 labeling revealed robust, tightly packed bundles within the AIS **(Fig. 3L,N)**. By contrast, in shDvl neurons, TRIM46 labeling was markedly reduced and appeared less organized, consistent with a thinner AIS structure **(Fig. 3M,O)**. Quantification confirmed a strong reduction in both the number of TRIM46-positive bundles (1.5 ± 0.5% vs 5.4 ± 0.6% in Ctrl neurons) and axon thickness (0.7 ± 0.1µm vs 1.3 ± 0.1µm in Ctrl neurons) following Dvl depletion **(Fig. 3P,Q)**. These results indicate that Dvl2 is required for proper cytoskeletal organization at the AIS and suggest that the AIS fragmentation phenotype is associated with defective microtubule bundle assembly or stabilization. Given that voltage-gated sodium channel clustering is a hallmark of AIS functional organization, we next examined Pan-Nav distribution in control and shDvl neurons. Whereas control neurons showed strong Pan-Nav enrichment along a single AIS, shDvl neurons displayed reduced Pan-Nav labeling together with the same spectrum of structural defects observed for AnkG, including fragmented or absent AIS-associated staining **(Fig. 3R-T)**. Quantification confirmed a significant 37% reduction in Pan-Nav fluorescence intensity in shDvl neurons, from 1.8 ± 0.1 × 10³ a.u. in controls to 1.2 ± 0.1 × 10³ a.u. following Dvl depletion **(Fig. 3U)**. Finally, since early Dvl depletion impaired AIS formation, we also asked whether Dvl proteins are required for maintenance of an already established AIS **(Fig. S6)**. To address this, neurons were transfected with shDvl at DIV13 and analyzed at DIV17, after AIS formation had largely occurred. In control neurons, Dvl2 and AnkG displayed continuous enrichment along the AIS, whereas Dvl-depleted neurons showed reduced Dvl2 staining together with fragmented AnkG-positive segments **(Fig. S6A)**. Quantification confirmed a significant 20% decrease in Dvl2 fluorescence intensity at the AIS following late shDvl expression, from 3.9 ± 0.3 × 10³ a.u. in control neurons to 3.1 ± 0.3 × 10³ a.u. in shDvl neurons **(Fig. S6B)**. Moreover, phenotypic analysis revealed a marked reduction in the proportion of neurons with a normal AIS, accompanied by an increase in fragmented and very fragmented AISs, with some neurons showing complete AIS loss **(Fig. S6C)**. Together, these data indicate that Dvl proteins are required not only for AIS formation during neuronal development, but also for the maintenance and structural integrity of the mature AIS. Together, these findings establish Dvl proteins as key organizers of AIS formation and maintenance. Dvl depletion alters AIS morphology and positioning, reduces multiple AIS-associated proteins, disrupts TRIM46-positive microtubule bundles, and impairs Pan-Nav clustering. Thus, Dvl loss compromises AIS compaction, structural stability, and molecular specialization, placing Dvl2 within the cytoskeletal and scaffolding mechanisms that preserve AIS structure and possibly neuronal excitability.

**Fig 3:**
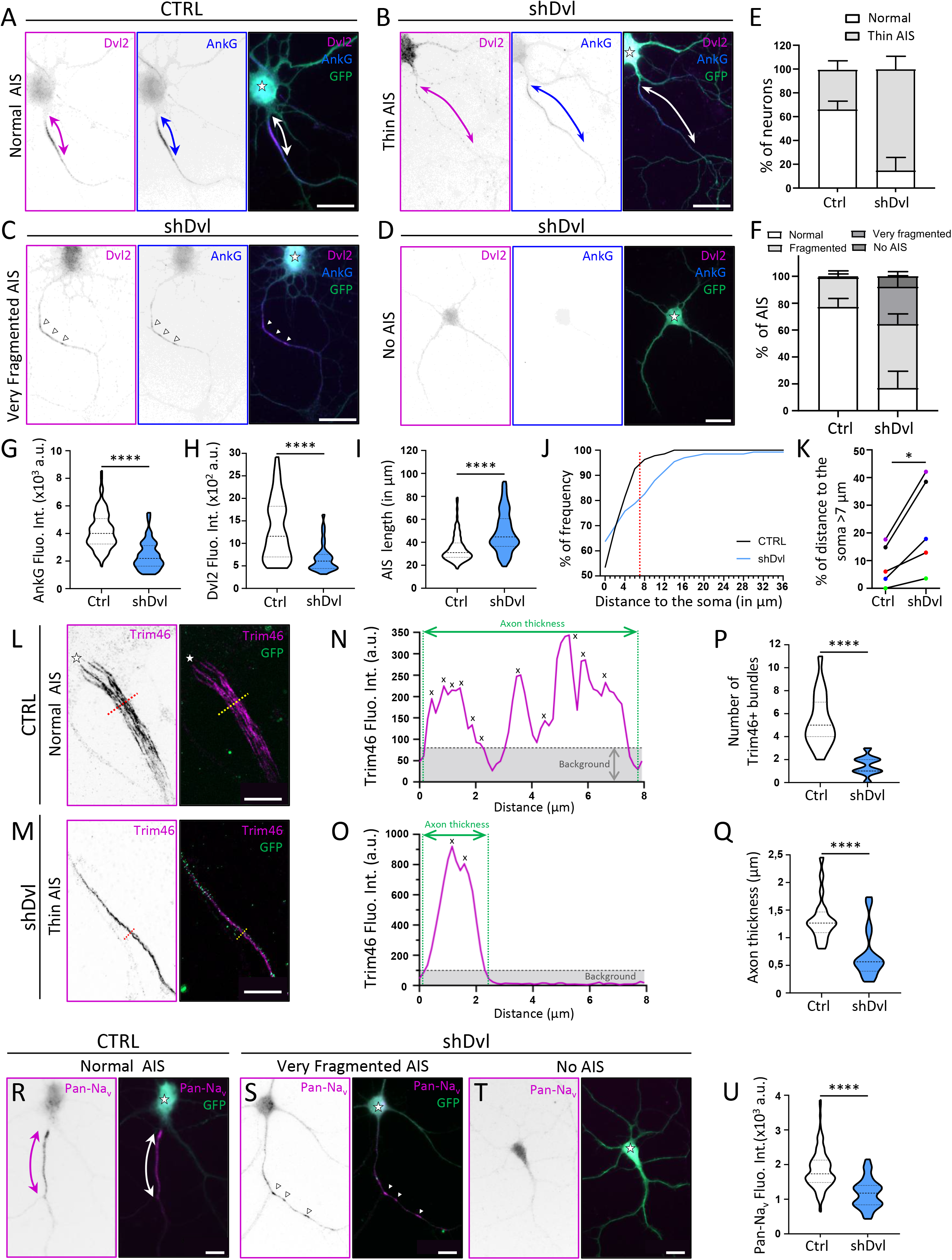
Dishevelled regulates AIS formation. (**A-D**) Illustration of DIV7 neurons transfected with a control or shDvl plasmid and labelled for Dvl2 (magenta) and AnkG (blue). Stars show the cell body and double arrows the AIS. Dvl-induced AIS fragmentation is illustrated as small, interspersed AnkG+ clusters as in (Very Fragmented AIS, arrowheads). (**E**) Percentages of neurons with a thinner AIS (as shown in (A ; “Thin AIS”)) with shDvl. Means ± SEM are shown. (**F**) Percentages of neurons per phenotype described as normal AIS (1 fragment), fragmented (2-3 fragments), very fragmented (≥4 fragments) and loss of AIS with shDvl. Means ± SEM are shown. (**G-H**) Quantifications of fluorescence intensity levels of AnkG and Dvl2 in shDvl neurons (excluding no AIS phenotypes) show downregulations in AnkG and Dvl2 levels. (**I**) Quantifications AIS length in shDvl neurons (excluding no AIS phenotypes) show AIS lengthening. **(J)** Relative frequency of distance to the soma indicates differences in the distribution of values, which can be delineated into two populations separated by the red dotted line at 7 µm. The starting point of the measurement is the intersection between the soma and the axon, or between the dendrite and the axon in the case of an AcD, whereas the end point is the beginning of the AIS. **(K)** Percentages of distances to the soma >7 µm (population to the right of the red dotted line) show an increased proportion across the different cultures (represented by the differently coloured dots). (**L-M)** Illustration of a DIV7 neuron transfected with a control or shDvl plasmid and labelled for Trim46 (magenta) after ultrastructure expansion microscopy (U-ExM). Red and yellow dotted line corresponds to Line Scan representation **(N-O)** of the area. Background is indicated in gray area under the black dotted line and MT bundles by a cross. (**P-Q**) Quantifications of Trim46+ MT bundles number or axon thickness on GFP labelling in shDvl neurons after U-ExM show strong downregulations respectively of MT bundles number or axon thickness. (**R-T**) Illustration of DIV7 neurons transfected with a control or shDvl plasmid and labelled for Pan-Na_v_ (magenta). Stars show the cell body and double arrows the AIS. Dvl-induced AIS fragmentation is also illustrated as small, interspersed Pan-Na_v_ + clusters as in (“Very Fragmented AIS”, arrowheads). (**U**) Quantifications of fluorescence intensity levels of Pan-Na_v_ in shDvl neurons show strong downregulations Pan-Na_v_ levels. Scale bars : 20 µm in A to D and P to R ; 4.6 µm in J and K (normalized to the expansion factor).

### Dvl knock-down *in vitro* changes AP initiation at the AIS Fig. 4, S7

At the AIS ion channels are enriched and anchored to the underlying cytoskeleton. We therefore assessed, whether the observed disruption of AIS organization after Dvl knock-down resulted in functional alterations of neuron properties and AIS excitability. We performed whole-cell patch-clamp recordings at DIV9 from nucleofected neurons that were identified by their GFP fluorescence **(Fig. 4A)**. Current-clamp recordings **(Fig. 4B)** showed that the resting membrane potential was similar between the two groups (control -61.1 ± 1.8 mV, n = 13; shDvl -57.1 ±1.5 mV, n = 15, unpaired t-test p = 0.1, **Fig. 4C**), input resistance was increased in shDvl neurons 1019 ± 164 MΩ (n = 16) compared to control neurons 540 ± 110 MΩ (n = 13, Mann Whitney test p = 0.02, **Fig. 4D**). This was in-line with a smaller cell body size of the sampled shDvl neurons of 250 ± 14 µm^2^ (n = 16) compared to control neurons (n = 13): 308 ± 19 µm^2^ (unpaired t-test p = 0.02). We hypothesized, that the disassembled AIS, in absence of Dvl2, interferes with the precise action potential (AP) initiation in the proximal AIS. We therefore investigated single APs as a read-out of AIS dependent excitability, to distinguish between AIS and somatic components of the action potential. Single APs were reliably evoked by short current injections of 3 ms **(Fig. 4E)**, allowing analysis of threshold, amplitude and half-width **(Fig. S7A)**. APs comparison between control and shDvl neurons revealed a 10-mV depolarized threshold in shDvl neurons compared to control neurons (shDvl: -24.7 ±1.2 mV, n = 16; vs. control: -34.3 ± 1.1 mV, n = 14, unpaired t-test p = 0.0001, **Fig. 4F**). Additionally, the action potential amplitude (Fig. 4H) was reduced by about 20 mV in shDvl neurons while the half-width was unchanged (control: 1.6 ± 0.1 ms, n = 14; shDvl: 1.9 ± 0.2 ms, n = 16, Mann-Whitney test, p = 0.75) suggesting that the somatically measured AP half-width is defined in majority by somatically located Nav and Kv channels. To investigate AP initiation at the AIS, we differentiated the AP voltage recordings to reveal the AIS and somatic components of each AP^7^ **(Fig. 4H, S7B)**. This analysis revealed a strong reduction in the first peak component of the first derivative **(Fig. 4I)** of the AP voltage, as well as of the maximum peak voltage change of the soma (**Fig. 4J**). Measuring the time delay between the first and second peak of the AP derivative revealed a longer delay for shDvl neurons of 550 ± 56 µs compared to control neurons: 330 ± 48 µs (Mann-Whitney test, p = 0.0035). Taken together, these results suggested that shDvl neurons have a reduced AP initiation at the AIS consistent with the disrupted AIS and a shift of the AIS to the distal axon reflected in the timing delay for the initiation of the axonal AP. However, the strong reduction of the axonal AP component did not abolish the capability to generate a similar maximum number of repetitive APs during 800 ms long current injections, although the threshold current was reduced in sh-Dvl neurons in-line with their lower measured input resistance **(Fig. S7C-E)**. At the end of the electrophysiology experiments, neurons were fixed and we performed post-hoc immunolabelling to correlate morphological parameters with electrophysiological parameters **(Fig. 4K-M)**. In agreement with other measurements (**Fig. 3**), we found that AIS onset was more variable and the AIS length was increased in the shDvl neurons **(Fig. 4L)**. Correlating the AIS onset versus the amplitude of the axonal action potential component revealed a correlation in the control neurons that was lost in shDvl expressing neurons **(Fig. 4M)**. Taken together, we show that overall neuronal activity is altered and that AIS action potential initiation and sodium channels are reduced after Dvl dependent AIS disruption. These data demonstrate that Dvl AIS organization is important for proper establishment of AIS function.

**Figure 4:**
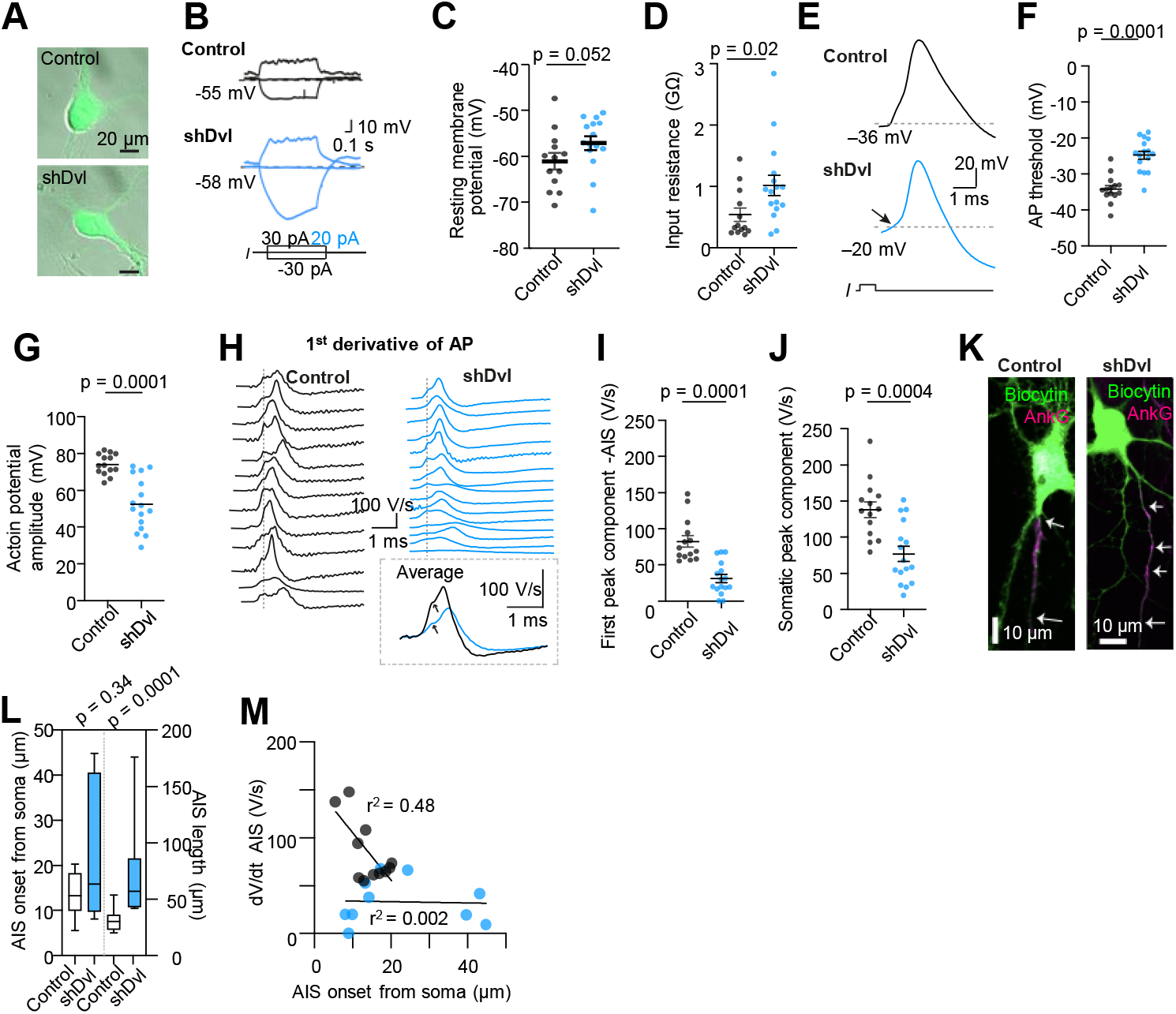
Axonal excitability is reduced in shDvl knockdown neurons. (A) Oblique contrast images overlayed with GFP fluorescence of control and shDvl neurons. (B) Example subthreshold voltage traces of a control (DIV 8) and a shDvl (DIV 10) neuron. (C) Summary plot showing resting membrane potential (D) Summary plot of input resistance reveals that the input resistance in shDvl neurons is consistently increased. (E) Examples of single action potentials (DIV 8) after a 3 ms current injection show a depolarized AP threshold indicated by a grey dotted line. (F) Summary data of AP threshold shows a more positive threshold in shDvl neurons suggesting a mainly somatic contribution (G) Summary plot of AP amplitude shows around 20 mV reduction in shDvl neurons. (H) 1st derivative of the action potential (dV/dt) from each neuron reveals a strong reduction of the AIS action potential component in shDvl neurons. This is also reflected in the averaged derivative for each condition in which all traces were aligned to their onset before averaging. (I) and (J) Summary plots of axonal and somatic component of the derived action potential (dV/dt). (K) Examples of biocytin filled and post-hoc labelled neurons for morphological analysis of recorded neurons. (L) Summary data of AIS onset and AIS length from recorded neurons (control: n = 11; shDvl: n = 10) show higher variability in the onset and a longer AIS in shDvl neurons in line with Fig.3I-K. (M) Correlation of the dV/dt of the AIS action potential component with the onset of the AIS determined from images as in (K) reveals a strong correlation in control neurons that is absent in the shDvl neurons, in-line with the strong reduction of the AIS AP. Data was fit with a linear equation: control Y = -4.961*X + 154.9, P = 0.017, R2 = 0.48; shDvl Y = -0.06304*X + 34.75, P = 0.91, R2 = 0.002.

### MTCL1 mediates selective recruitment of Dvl2 to microtubules and supports its compartmentalization in the AIS. Fig. 5/S8

To test whether Dvl2 recruitment to microtubules (MT) depends on specific AIS-resident microtubule-associated proteins, COS7 cells were co-transfected with HA-Dvl2 together with either TRIM46-myc or GFP-MTCL1. In cells expressing TRIM46-myc, HA-Dvl2 retained its typical punctate, condensate-like distribution^66^ and did not show evident relocalization onto TRIM46-positive MT bundles **(Fig. 5A)**. By contrast, co-expression of GFP-MTCL1 caused a marked redistribution of HA-Dvl2 condensates toward the MTCL1-decorated MT network, indicating that MTCL1, but not TRIM46, under these conditions, efficiently promotes Dvl2 association with microtubules **(Fig. 5B)**. This difference was confirmed by colocalization analysis. Mander’s coefficient showed that, in MTCL1-positive cells, approximately 70% of the HA-Dvl2 signal overlapped with GFP-MTCL1-decorated microtubules, indicating robust recruitment of overexpressed Dvl2 to these structures. In contrast, TRIM46-transfected cells displayed only minimal overlap between HA-Dvl2 puncta and the TRIM46-positive microtubule network **(Fig. 5C)**. Together, these data identify MTCL1 as an efficient recruiter of Dvl2 to microtubules in this assay. To determine whether MTCL1 microtubule-binding regions are required for Dvl2 recruitment, HA-Dvl2 was co-expressed in COS7 cells with V5-tagged wild-type MTCL1 or ΔKR, ΔN, and ΔKRΔN mutants^88^ (**Fig. S8)**. Wild-type MTCL1 recruited Dvl2 to the MTCL1-positive microtubule network, whereas the ΔKR mutant lost filamentous localization and was recruited to HA-Dvl2 condensates, suggesting interdependent MTCL1–Dvl2 localization. In contrast, Dvl2 recruitment to MTCL1-positive structures was strongly reduced with ΔN and ΔNΔKR mutants, indicating that the ΔN-mutated region is required for efficient Dvl2 recruitment to microtubules **(Fig. S8)**. In addition, reciprocal co-immunoprecipitation demonstrated that Dvl2 and MTCL1 are present in the same molecular complex, indicating that MTCL1-mediated recruitment of Dvl2 to microtubules is accompanied by a detectable physical association between the two proteins **(Fig. 5D)**. To evaluate the relevance of this interaction in neurons, we examined whether MTCL1 depletion affects Dvl2 recruitment to the AIS. In shMTCL1 neurons, Dvl2 signal along the AIS was visibly reduced compared with control neurons **(Fig. 5E)**. Conversely, V5-MTCL1 was enriched along the AnkG-positive proximal axonal segment in control neurons, whereas this enrichment was reduced and more discontinuous following Dvl depletion **(Fig. 5F)**. Together, these reciprocal effects suggest that Dvl proteins and MTCL1 cooperate in the organization or maintenance of a specialized MT-rich neuronal compartment, and that perturbation of either partner compromises the accumulation of AIS-associated components and Dvl2 enrichment at the AIS. Overall, these data support a model in which MTCL1 acts as a microtubule-associated adaptor that recruits Dvl2 from condensate-like cytoplasmic assemblies onto MTs and promotes its enrichment in a polarized neuronal microtubule domain.

**Fig. 5.**
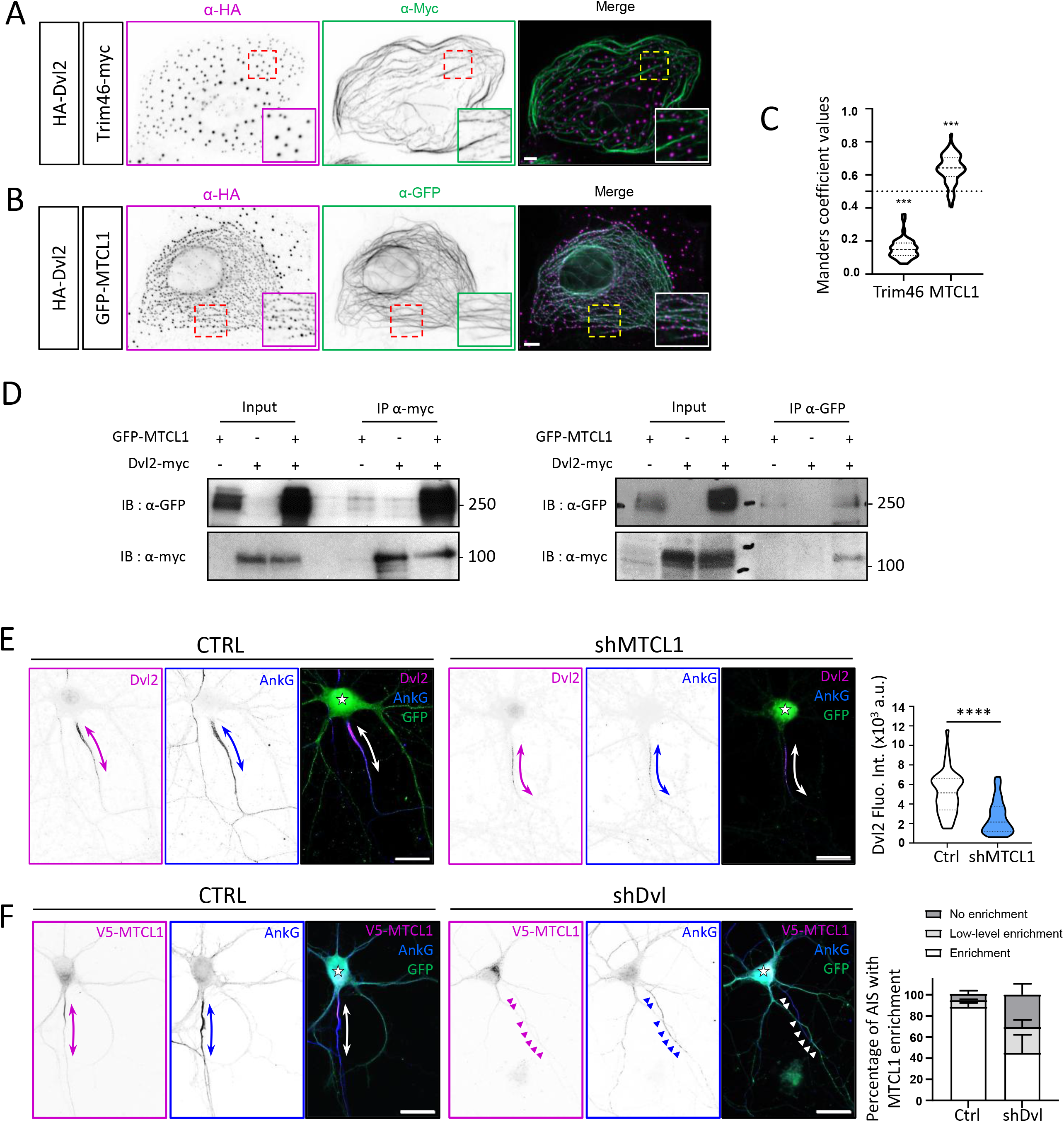
Dvl2 is recruited by MTCL1 to MT. **(A, B)** COS7 cells transfected for 24hrs with HA-Dvl2 (magenta) either with (A) Trim46-myc (green) or (B) GFP-MTCL1 (green). While HA-Dvl2 forms liquid-like condensates that are independent of Trim46+ microtubules (MTs), HA-Dvl2 is recruited in 64% of cells cotransfected with GFP-MTCL1 **(C)** Quantification of the specific HA-Dvl2 recruitment to MTCL1+ MTs assessed by Mander’s colocalization coefficients derived with JACoP plugin (0 indicates no colocalization, 1 indicates perfect colocalization) in Trim46-myc and GFP-MTCL1–transfected cells. The Mander’s coefficient indicates that ∼70% of the HA-Dvl2 detected overlaps with GFP-MTCL1-decorated microtubules. **(D)** Co-immunoprecipitation of GFP-MTCL1 with Dvl2-myc shows immunoprecipitated GFP-MTCL1 with anti-myc and immunoprecipitated Dvl2-myc with anti-GFP. Protein sizes are estimated in kDa. (**E**) Illustration of DIV7 neurons transfected with a control or shMTCL1 plasmid and labelled for Dvl2 (magenta) and AnkG (blue). Stars show the cell body and double arrows the AIS. Quantifications of fluorescence intensity levels of Dvl2 in shMTCL1 neurons shows strong downregulation Dvl2 levels. (**F**) Illustration of DIV7 neurons co-transfected with V5-MTCL1 and either a control or shDvl plasmid and labelled for V5-MTCL1 (magenta) and AnkG (blue). Stars show the cell body and double arrows the AIS. Percentages of neurons with an enrichment of V5-MTCL1 to the AIS identified by the AnkG staining show a decrease in MTCL1 enrichment at the AIS with the shDvl. Means ± SEM are shown. Scale bars : 5 µm in A and B ; 20 µm in E and F

### MTCL1-mediated recruitment of Dvl2 to microtubules requires the DIX and PDZ domains Fig. 6

To determine which domains of Dvl2 are required for its recruitment to MTCL1-decorated microtubules, COS7 cells were co-transfected with GFP-MTCL1 and either wild-type HA-Dvl2 or Dvl2 deletion mutants lacking the DIX, PDZ, or DEP domain for 24 h **(Fig. 6A)**. As expected, wild-type Dvl2 formed condensate-like structures and showed robust recruitment to GFP-MTCL1-positive microtubules. In contrast, deletion of the DIX domain abolished condensate formation and strongly impaired Dvl2 recruitment to MTCL1-positive microtubules. Deletion of the PDZ domain did not prevent condensate formation, although the condensates appeared less spherical, and only partial recruitment or juxtaposition to GFP-MTCL1-positive microtubules was observed. By comparison, deletion of the DEP domain had no major effect on Dvl2 recruitment, as the mutant remained efficiently associated with MTCL1-decorated microtubules **(Fig. 6A)**. These observations were supported by quantitative analysis. Scoring of cells displaying Dvl2 recruitment to GFP-MTCL1-positive microtubules showed a marked reduction for both the ΔDIX and ΔPDZ mutants compared with wild-type Dvl2, whereas the ΔDEP mutant behaved similarly to the wild-type protein **(Fig. 6B)**. Recruitment increased after 48 h in all conditions, but remained markedly impaired for ΔDIX and predominantly partial for ΔPDZ. In contrast, ΔDEP recruitment approached that of wild-type Dvl2 at this later time point, suggesting slower recruitment kinetics rather than a strict requirement for the DEP domain **(Fig. 6C)**. Together, these data indicate that efficient MTCL1-mediated recruitment of Dvl2 requires the DIX and PDZ domains, whereas the DEP domain is dispensable in this context.

**Fig. 6.**
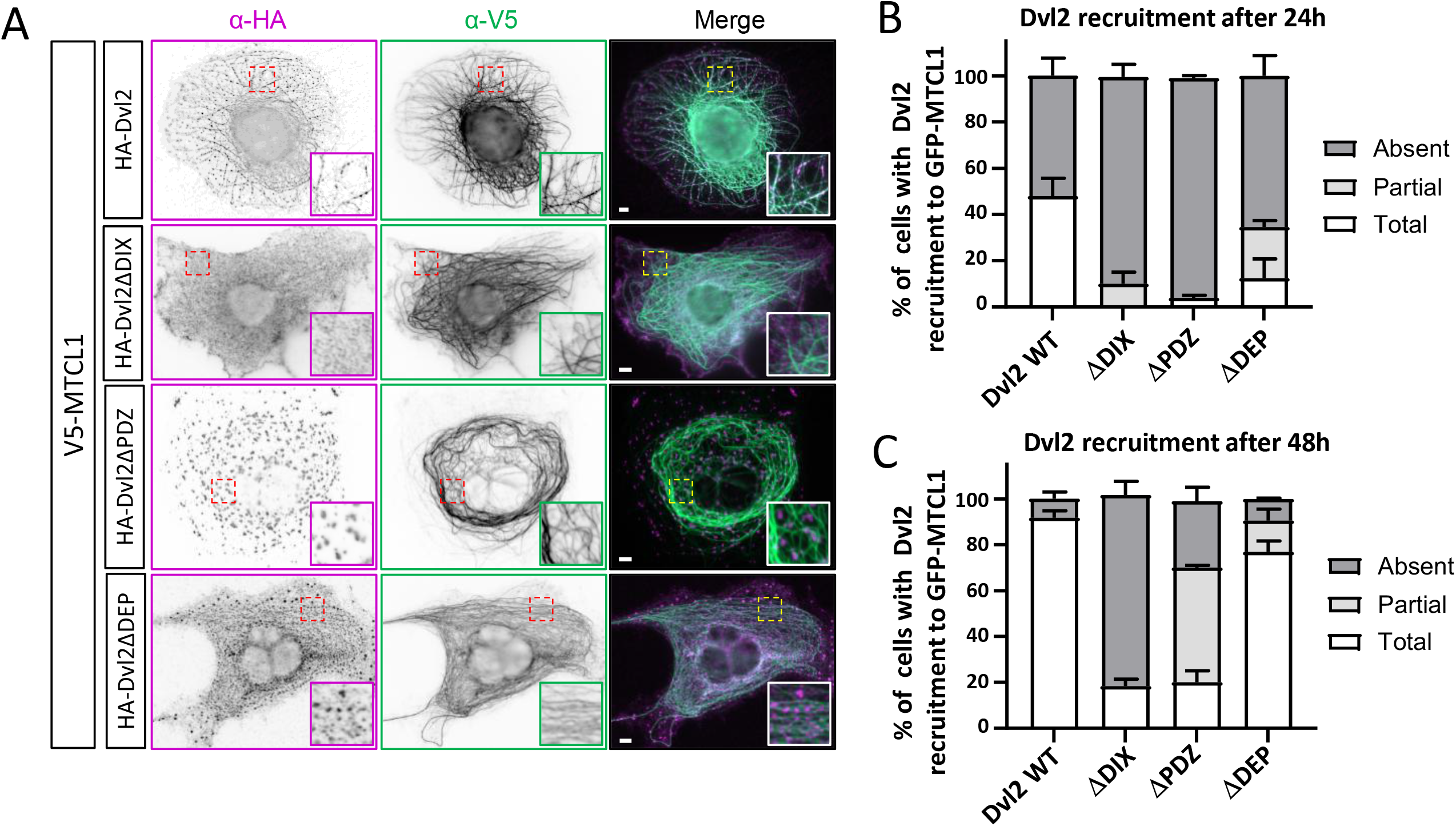
Dvl2 recruitment to MT is DIX and PDZ dependent. **(A)** COS7 cells transfected for 24hrs with GFP-MTCL1 either with HA-Dvl2 or Dvl2 deletion mutants for the DIX, PDZ, and DEP domains. **(B)** Quantification of the specific HA-Dvl2 recruitment to MTCL1+ MTs after 24h. HA-Dvl2DDIX is no longer able to form condensates and is no longer recruited to GFP-MTCL1+ MTs. HA-Dvl2DPDZ is still able to form condensates (that resume non spherical shape) that are partially recruited and juxtaposed to GFP-MTCL1+ MTs. DEP loss has no major impact on Dvl2 recruitment. **(C)** Quantification of the specific Dvl2-myc recruitment to MTCL1+ MTs after 48h shows the same as described for 24h transfection, except that percentages of recruitment are higher for all the conditions. Scale bar : 5 µm in A

### Evolved Dvl2 PDZ domain binders selectively block its function and perturb its condensate organization and neuronal AIS accumulation. Fig. 7, 8 and S9

To target the Dvl2 PDZ domain intracellularly, we employed a protein directed evolution approach to generate binders suitable for intracellular engagement and imaging ^89,90^. Using recombinant Dvl2 PDZ domain as bait, we screened a 10Fn3-based phage display library ^90,91^ and isolated seven high-affinity clones after three rounds of selection, as assessed by phage-ELISA (**Fig. 7A**). We prioritized the top two candidates, Xph335 and Xph334 (**Fig. 7B**), for further characterization. AlphaFold3 ^92^ predicted the structures of the Xph335-Dvl2 and Xph334-Dvl2 complexes, with notably high confidence for Xph335 (**Fig. 7C**). In both cases, the diversified FG loop, the main interaction site, adopted a β-hairpin conformation that occupied the canonical PDZ domain binding groove. PDZ domains classically recognize C-terminal motifs, typically ending with a hydrophobic residue at the P₀ position and a serine or threonine at P₋₂. However, numerous studies have demonstrated that PDZ domains, particularly those of Dishevelled proteins, can also engage internal sequences with high affinity ^93–95^. The Dvl2 PDZ domain indeed exhibits a remarkably flexible binding cleft that can accommodate both C-terminal and internal ligands, with the latter presenting a conserved aspartate side chain to substitute for the C-terminal carboxylate^96^. The predicted common binding mode for Xph335 and Xph334 exemplifies this domain flexibility but does not require an aspartate residue to mimic the C-terminal carboxylate of canonical ligands. By occupying the PDZ domain groove, as illustrated by the X-ray crystal structure of Dvl2 bound to a canonical C-terminal peptide, pepC1 ^96^, Xph335 and Xph334 would therefore sterically block partner recognition. To confirm these predictions, we performed epitope binning using the canonical Dvl2 ligand pepC1^96^. Increasing concentrations of pepC1 displaced both evolved binders, confirming competitive occupancy of the PDZ domain binding groove by Xph335, Xph334, and pepC1 (**Fig. 7D**). Together, these data demonstrate that Xph335 and Xph334 bind the canonical PDZ groove and thus can serve as selective inhibitors of Dvl2 protein–protein interactions. We next assessed whether these binders recognize Dvl2 condensates in cells and perturb their organization. In COS-7 cells co-expressing HA-Dvl2 and mScarlet alone (control), HA-Dvl2 was observed predominantly in condensates of standard size, whereas mScarlet displayed a diffuse cytoplasmic signal with nuclear enrichment. In contrast, upon co-expression of Xph334-mScarlet or Xph335-mScarlet, both binders accumulated at HA-Dvl2-positive condensates in cells displaying condensed phenotypes, indicating intracellular recognition of Dvl2 assemblies **(Fig. 7D)**. This was supported by Manders coefficient analysis, which increased from a mean value of 0.34 in the mScarlet control to 0.78 and 0.83 for Xph334 and Xph335, respectively, consistent with robust colocalization of both binders with Dvl2 condensates **(Fig. 7E)**. Importantly, binder expression also altered the distribution of Dvl2 condensate phenotypes. In the control condition, 75% of cells displayed standard condensates, 17% displayed small condensates, and 8% showed a diffuse Dvl2 pattern **(Fig. 7F)**. In the presence of Xph334, these values shifted to 41%, 36%, and 23%, respectively. In the presence of Xph335, they shifted further to 30%, 14%, and 56%, respectively **(Fig. 7F)**. These statistically significant changes indicate that Xph334 and Xph335 do not behave as inert reporters under these conditions, but instead interfere with the formation and/or maintenance of normal Dvl2 condensates. In addition, we next evaluated the intracellular specificity of these binders against additional PDZ-domain-containing proteins, namely the AIS-associated proteins Scribble (Scrib) ^54^ and PSD93 ^97^, and Shank3, a scaffold protein known to undergo condensate formation ^98,99^. COS-7 cells were co-transfected with GFP-Scrib, PSD93-mGL, or GFP-Shank3, together with either mScarlet, Xph334, or Xph335 **(Fig. S9)**. In all cases, no detectable colocalization was observed, indicating that Xph334 and Xph335 do not measurably interact with Scribble, PSD93, or Shank3 under these experimental conditions **(Fig. S9)**. Taken together, these data show that Xph334 and Xph335 are selective intracellular binders of Dvl2 that robustly recognize Dvl2 condensates while concomitantly perturbing their organization in COS-7 cells. Finally, we asked whether the Dvl2-selective PDZ binders identified in heterologous cells also affect Dvl2 organization in neurons **(Fig. 8)**. To do so, we expressed mScarlet alone, Xph334, or Xph335 in cultured neurons and examined endogenous Dvl2 together with AnkG. Compared with control neurons, neurons expressing Xph334 or Xph335 showed an altered distribution of both Dvl2 and AnkG along the proximal axon, with the AIS appearing shortened, discontinuous, or fragmented rather than forming a single continuous segment **(Fig. 8A)**. Consistent with these observations, quantification revealed a shift from predominantly normal AIS profiles in control neurons toward fragmented, and for Xph335 particularly, very fragmented AIS phenotypes, with the strongest and statistically significant effect observed for Xph335 **(Fig. 8B)**. Together, these data indicate that selective interference with the Dvl2 PDZ domain is sufficient not only to perturb Dvl2 condensate organization in non-neuronal cells, but also to impair the normal accumulation of Dvl2 in the proximal axon and compromise AIS integrity in neurons.

**Figure 7.**
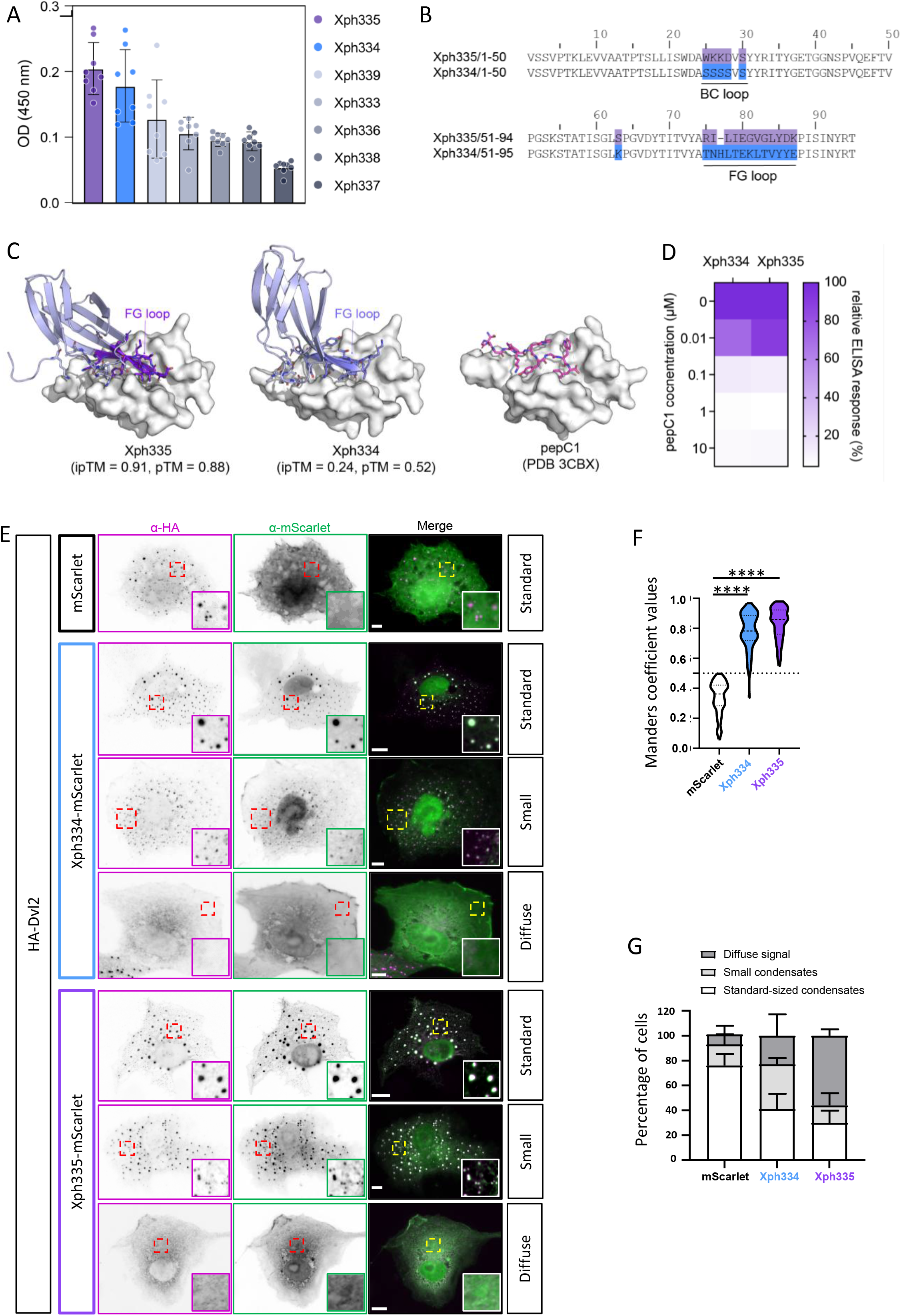
Xph334 and Xph335 colocalize with HA-Dvl2 condensates and impair their normal assembly in COS-7 cells. (**A**) Phage-ELISA screening of evolved Dvl2 PDZ domain binders (Xph333–Xph339). Binding responses were measured across 8 independent colonies for each binder. Data are presented as mean ± SD, with binders sorted by response level. (**B**) Amino acid sequences of the top-performing binders, Xph335 and Xph334. Diversified regions, primarily corresponding to the BC and FG loops, are highlighted in purple (Xph335) and blue (Xph334). (**C**) Structural predictions of Xph335 and Xph334 in complex with the Dvl2 PDZ domain, generated using AlphaFold3. For comparison, the X-ray crystal structure of Dvl2 bound to a canonical PDZ-binding peptide (pepC1, PDB 3CBX) is shown. In all structures, the Dvl2 PDZ domain is depicted as a grey surface in the same orientation. Binder residues within 5 Å of Dvl2 are shown as sticks, while others are in cartoon representation. Confidence metrics for AlphaFold3 predictions include the predicted template modeling (pTM) and interface pTM (ipTM) scores. The models suggest that both evolved binders engage the Dvl2 PDZ domain primarily via their FG loops (purple for Xph335, blue for Xph334), occupying the canonical PDZ domain-binding groove. (**D**) Epitope binning of Xph335 and Xph334 against pepC1 by phage ELISA. Each value represents the mean of 3 independent colonies. Increasing concentrations of pepC1 reduce binding of Xph335 and Xph334, indicating competition for an overlapping epitope on the Dvl2 PDZ domain. (**E**) Immunofluorescence analysis of COS-7 cells co-transfected with plasmids encoding HA-Dvl2 and either mScarlet alone (control), Xph334-mScarlet, or Xph335-mScarlet. In control cells, HA-Dvl2 predominantly formed condensates of standard size, whereas mScarlet displayed a diffuse cytoplasmic signal with nuclear enrichment. In cells expressing Xph334-mScarlet or Xph335-mScarlet, HA-Dvl2 exhibited three phenotypic classes: standard condensates, small condensates, or a diffuse pattern. In cells retaining visible condensates, both binders colocalized with HA-Dvl2-positive structures. (**F**) Quantification of colocalization between HA-Dvl2 and mScarlet or binder-mScarlet using the Manders coefficient. A threshold of 0.5 was used to distinguish absence from presence of colocalization. (**G**) Quantification of the percentage of cells displaying standard condensates, small condensates, or diffuse HA-Dvl2 signal in each condition. In control cells, 75% of cells displayed standard condensates, 17% small condensates, and 8% a diffuse pattern. In the presence of Xph334, 41% of cells displayed standard condensates, 36% small condensates, and 23% diffuse HA-Dvl2. In the presence of Xph335, the corresponding values were 30%, 14%, and 56%. Differences between conditions were significant. Scale bar : 5 µm in D

**Fig 8:**
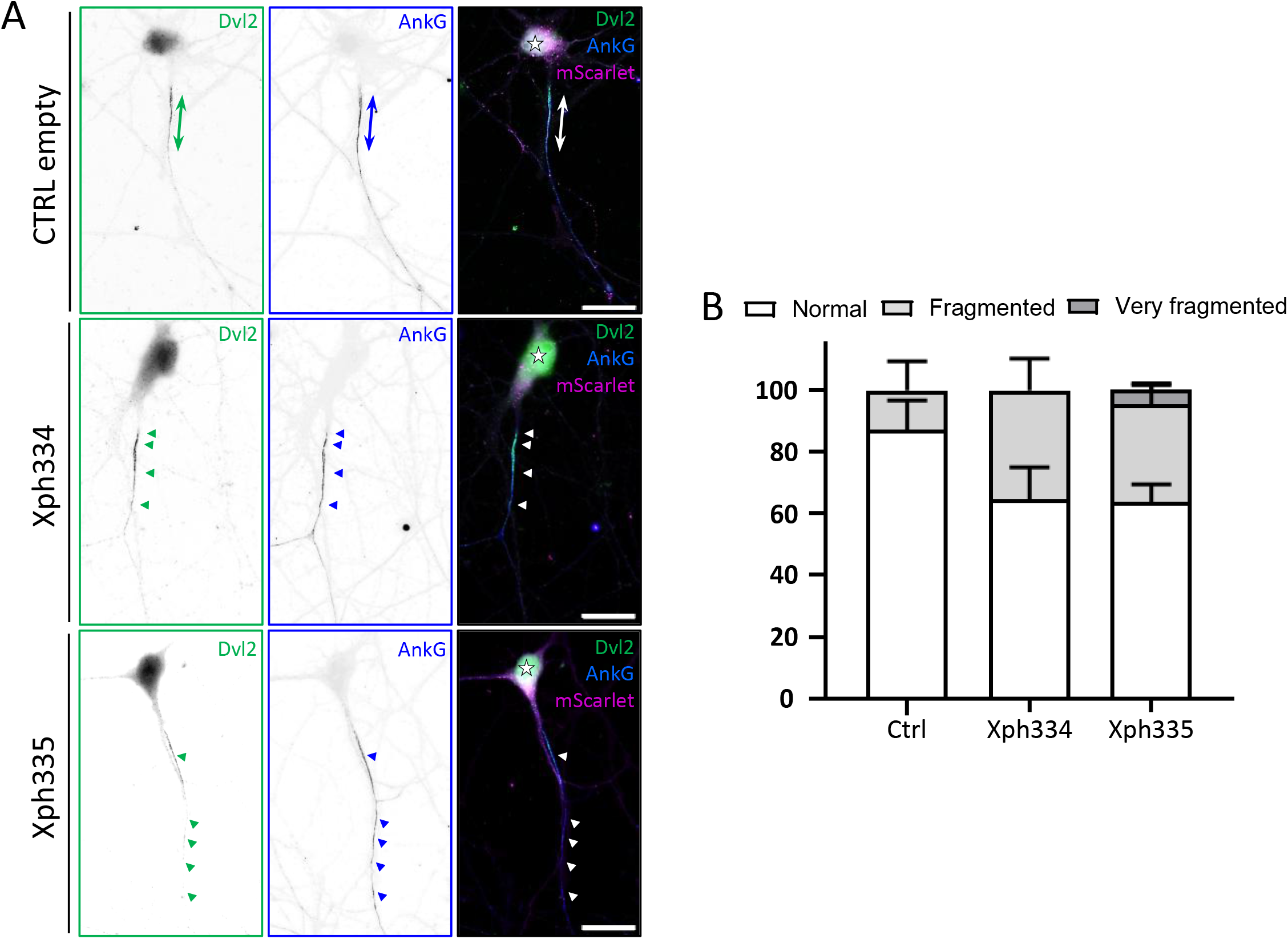
PDZ-specific inhibition of Dvl2 impairs AIS formation. (**A**) Illustration of DIV7 neurons transfected with a control (mScarlet), Xph334 or Xph335 plasmid and labelled for Dvl2 (green) and AnkG (blue). Stars show the cell body and double arrows the AIS. Dvl-induced AIS fragmentation is illustrated as small, interspersed AnkG+ clusters. (**B**) Percentages of neurons per phenotype described as normal AIS (1 fragment), fragmented (2-3 fragments) and very fragmented (≥4 fragments) with binders. Means ± SEM are shown. Scale bar : 20 µm in A

## DISCUSSION

Our study identifies Dvl2 as a functional component of the AIS and a central organizer of its proximal microtubule cytoskeleton. Dvl2 is enriched at the AIS *in vivo* and *in vitro*, where it localizes upstream of Ankyrin-G and closely associates with the microtubule cross-linkers TRIM46 and MTCL1 from early AIS formation through neuronal maturation. Dvl depletion disrupts AIS architecture, microtubule organization, AIS protein accumulation, and action potential initiation. Mechanistically, MTCL1 recruits Dvl2 to microtubules through its DIX and PDZ domains, while PDZ-targeting intracellular binders perturb Dvl2 condensates and reproduce AIS defects. These findings extend the emerging role of biomolecular condensates in neuronal organization^72,100^ to the AIS and support condensate-dependent Dvl2 organization as a determinant of AIS integrity.

### Dvl2 marks a proximal AIS compartment linked to microtubule organization

A major finding is that Dvl2 defines a proximal AIS subdomain between the soma and the main Ankyrin-G-positive compartment **(Fig.1 and Fig. S1-S3)**. This is particularly relevant given the structural and molecular heterogeneity of the AIS, whose proximal and distal modules support distinct functions^5,6^. The progressive displacement of Dvl2 relative to Ankyrin-G from DIV4 onward further indicates that Dvl2 is dynamically positioned during AIS maturation rather than passively recruited with Ankyrin-G, consistent with a specialized role in early AIS assembly and cytoskeletal organization. Moreover, Dvl2, but not Dvl3, was enriched at the AIS despite validated antibody specificity **(Fig.S3-S4)**, suggesting isoform-specific recruitment through distinct sequence features, partners, or biophysical properties. However, Dvl3 detection in a TRIM46-associated BioID screen^101^, may reflect methodological differences or an antibody-undetected AIS pool, highlighting the need to clarify Dishevelled isoform specialization in neuronal polarity. The apparent close association of Dvl2 with TRIM46 supports its role in microtubule organization **(Fig.2)**. Although TRIM46 promotes proximal axonal microtubule bundling and neuronal polarity^23,24^, recent *in vivo* studies suggest it governs MT fasciculation rather than AIS integrity itself^102,103^. Expansion microscopy and STORM localized Dvl2 puncta along stable MT bundles within the AIS, supporting its incorporation into a proximal MT-associated scaffold, although this does not establish direct MT binding. Finally, the presence of Dvl2 as early as DIV1, during axon specification, suggests that it contributes to the establishment of axonal identity rather than being recruited only after AIS assembly. Together, these findings identify Dvl2 as an early, *bona fide* MT-associated component of the AIS.

### Dvl proteins are required for AIS assembly and excitability

The functional importance of Dvl2 at the AIS is demonstrated by the pronounced defects caused by early Dvl depletion, including thinner, elongated, fragmented, or absent AnkG-positive segments. The concomitant reduction in AnkG, Dvl2, TRIM46, Pk2, Scrib, and Pan-Nav indicates that Dvl coordinates the assembly of structural, signaling, and excitable AIS components **(Fig.3 and Fig.S5)**. Notably, expansion microscopy revealed reduced and disorganized TRIM46-positive bundles, indicating defective MT assembly or stabilization as a core feature of the phenotype **(Fig. 3)**. The increased length but reduced compaction of residual AISs suggests that Dvl loss impairs consolidation of a coherent proximal axonal domain rather than AIS formation itself, implicating Dvl2 in AIS boundary refinement and internal organization **(Fig. S6)**. The electrophysiological data further demonstrate that these structural defects are functionally consequential **(Fig.4 and Fig.S7)**. Dvl knockdown depolarized action potential threshold, reduced action potential amplitude, weakened the axonal component of the spike, and increased the delay between axonal and somatic components of the AP derivative. Together with the reduction in Pan-Nav labeling and the distal shift in AIS onset, these observations strongly support the conclusion that Dvl-dependent disruption of AIS architecture compromises normal spike initiation at the AIS. The fact that repetitive firing was preserved, despite clear defects in AIS-dependent initiation parameters, is consistent with the idea that somatodendritic conductances remain sufficient to support ongoing firing once threshold is reached, while the specialized efficiency of the AIS trigger zone is selectively impaired. In this sense, the physiological phenotype matches the structural one: the AIS is not entirely absent in all cells, but its organization is degraded enough to weaken its normal role as the privileged site of action potential initiation.

### MTCL1 mediates Dvl2 recruitment to proximal axonal microtubules

Our observations extend the concept that Dishevelled include regulation of MT-associated biological functions including their stability^74,75^, spindle dynamics^104^, centrosome behavior^105,106^ and cilia growth^84,107^. However, Dishevelled recruitment to AIS microtubules remains unclear and may rely on indirect, context-dependent interactions rather than direct binding^108^. Supporting this model, an MT-binding-deficient MTCL1 mutant fails to recruit HA-Dvl2 to MTs and instead partitions into HA-Dvl2 condensates, identifying MTCL1 as a key link between Dvl2 and the microtubule network **(Fig.S8)**. These findings extend the classical view of Dishevelled as a Wnt/PCP transducer^55–57^, highlighting its broader role as multivalent scaffold proteins that assemble signaling complexes ^65–67^ into biomolecular condensates^68–71^. Together with our previous findings on Pk2, these data support a broader model in which core PCP proteins, including Dvl2, act locally at the AIS to organize the proximal axonal cytoskeleton and establish subcellular polarity. Mechanistically, our results identify MTCL1 as a key partner in Dvl2 recruitment to MTs and AIS compartmentalization **(Fig.5 and Fig.S8)**. In COS-7 cells, MTCL1, but not TRIM46, redistributed Dvl2 from punctate condensates onto microtubules, while reciprocal co-immunoprecipitation confirmed their association. In neurons, MTCL1 depletion reduced Dvl2 accumulation at the AIS, whereas Dvl depletion disrupted proximal MTCL1 enrichment, indicating reciprocal functional cooperation. We therefore propose that MTCL1 recruits or stabilizes Dvl2 on proximal axonal microtubules, while Dvl2 supports the organization of the MTCL1-positive scaffold. This mechanism links a signaling scaffold to a structural microtubule organizer and may explain the selective enrichment of Dvl2 at the proximal AIS.

### Dvl2 condensate organization is functionally important for AIS integrity

Our domain-mapping experiments refine this mechanism by showing that the DIX and PDZ domains, but not the DEP domain, are required for efficient MTCL1-mediated recruitment of Dvl2 **(Fig.6)**. The requirement for the DIX domain is notable because this domain is crucial for Dishevelled oligomerization^66^. Its loss nearly abolished condensate formation and impaired recruitment to MTCL1-positive MTs, indicating that Dvl2 higher-order assembly enables engagement with microtubule-associated machinery. This supports reciprocal crosstalk between cytoskeletal filaments and condensates, with MTs acting as platforms that concentrate or pattern phase-separated assemblies^109,110^. We propose that Dvl2 recruitment to neuronal microtubules partly relies on LLPS rather than a strictly stoichiometric interaction, although the contribution of condensates to AIS assembly and proximal axonal organization requires further investigation. The PDZ domain was required through a distinct mechanism: its deletion preserved condensate formation but produced distorted assemblies with reduced recruitment to MTCL1-decorated MTs. Thus, the PDZ domain likely mediates interactions that capture, stabilize, or orient Dvl2 assemblies on the MT lattice. Altogether, these results support a model in which Dvl2 recruitment to AIS-associated microtubules depends on both self-assembly and selective protein–protein interactions. Because several microtubule-associated proteins undergo LLPS and organize microtubules through condensate-like states^111^, future studies should test whether MTCL1 phase-separates alone or with Dvl2 and contributes to AIS microtubule bundling. Supporting this possibility, FuzDrop^112^ predicts maximal LLPS propensity for human MTCL1 (UniProtKB Q9Y4B5; pLLPS = 1.0000), suggesting that MTCL1 may promote Dvl2-associated condensates at the AIS. This suggests that MTCL1 may actively contribute to Dvl2-associated condensate formation at the AIS. The experiments using PDZ-directed binders provide an important complementary line of evidence **(Fig.7 and Fig.S9)**. Xph334 and Xph335 selectively targeted Dvl2 condensates without detectable colocalization with Scrib, PSD93, or Shank3. mportantly, these binders were not neutral reporters but shifted Dvl2 toward smaller or more diffuse assemblies in COS-7 cells. They therefore act as condensate-modifying tools with dissolver-like activity, consistent with emerging concepts of condensate-targeting molecules^113,114^. In neurons, the binders disrupted Dvl2 and AnkG organization and increased AIS fragmentation, with Xph335 having the strongest effect. These findings indicate that Dvl2 organization and PDZ-dependent interactions are required for AIS accumulation and maintenance. These findings support Dvl2 condensates as functional assemblies rather than overexpression artifacts and establish the binders as tools for studying Dvl2 condensate function in neuronal, Wnt-driven, and cancer-related contexts^115,116^.

### A possible convergence between PCP scaffolding and local Wnt/GSK3 signaling at the AIS

An additional possibility raised by our data is that Dvl2 function at the AIS may extend beyond PCP signaling to intersect with the canonical Wnt pathway, where Dishevelled may promote GSK3 inhibition and β-catenin stabilization^55^. Consistently, GSK3 and β-catenin regulate AnkG and voltage-gated sodium channel accumulation at the AIS, thereby modulating neuronal excitability^117^. Chronic lithium treatment also restored inhibitory function and neuronal excitability through GSK-3β inhibition in an *Ank3* variant mouse model, linking this pathway to AnkG-dependent disease mechanisms^118^. Because Dvl depletion reduces AnkG and Pan-Nav accumulation and impairs action potential initiation, Dvl2 may integrate with local GSK3/β-catenin signaling to regulate AIS cytoskeletal organization and sodium channel stabilization. Although speculative, this model is consistent with the GSK3-dependent control of MT stability by Dishevelled.

### Limitations and perspectives

Our study has several limitations that should be considered. First, shDvl broadly targets Dishevelled, leaving open whether the AIS phenotype reflects redundancy or cooperation among Dvl proteins, despite the apparent absence of Dvl3 enrichment. Second, while our imaging and recruitment assays strongly support an association of Dvl2 with AIS microtubules, the precise biochemical interface between Dvl2, MTCL1, and other AIS proteins remains unresolved. Third, although informative, the COS-7 assay does not reproduce AIS complexity and relies on interactor overexpression. At endogenous levels, other partners, including TRIM46, may also recruit Dvl2 to microtubules, particularly given the concentration- and stoichiometry-dependent nature of LLPS^119^, which may influence Dvl2 condensate behavior and partner recruitment. These limitations do not alter the central conclusion but define key questions for future investigation. It will be critical to determine whether Dvl2 directly regulates microtubule stability, bundling, or post-translational modifications in the proximal axon. It will also be important to establish whether Dvl2 acts upstream of AnkG recruitment, in parallel with TRIM46/MTCL1-dependent microtubule organization, or in a feedback loop linking these processes. Another key issue is whether Dvl2 at the AIS is controlled by upstream Wnt/PCP signals or whether it performs a locally adapted scaffold function largely independent of canonical ligand-triggered signaling. Finally, Dvl2-dependent AIS mechanisms should be examined in disorders involving axonal excitability, such as autism and epilepsy, where PCP signaling and the AIS may represent convergent pathogenic hubs ^120^

## METHODS

### Experimental model

#### Animals

All procedures involving animals were performed in accordance with the European Union Directive 2010/63/EU. Mice and rats had ad libitum access to food and water and were housed in polypropylene cages under controlled conditions (22 ± 2°C; 12 h light/12 h dark cycle, lights on from 07:00 to 19:00). Sprague–Dawley rats were obtained from Janvier Labs and used at ages ranging from embryonic day 18.5 (E18.5) to postnatal day 25 (P25). C57BL/6J mice were bred in-house in the animal facility and used between postnatal days 7 and 21 (P7–P25).

### Method details

#### Plasmids

The shRNA were cloned into a pSuper vector (Oligoengine) based on previously published sequences for shDvl targeting all three *Dvl* isoforms ^76^ and for shMTCL1 ^25^. pCDNA3.1 HA-Dvl2 vector was kindly provided by Pr. M. Bienz (Cambridge, UK) ^66^. The DIX, PDZ, or DEP domains were deleted from HA-Dvl2 by site-directed mutagenesis using the Q5® Site-Directed Mutagenesis Kit (NEB). HA-Dvl1 and HA-Dvl3 constructs were generated by cloning different mouse *Dvl1* and *Dvl3* sequences (kind gift of Dr. C. Duplaa, INSERM U1034, France) using PCR and inserted into the pcDNA3-3xHA to create in-frame fusions with the 3xHA sequence. pCDNA3.1 hDvl2-mycHIS was a kind gift from Pr. S. Sokol (MSSM, New York City, USA). pcDNA3-hDvl2-PDZbinder334-mScarlet3 and pcDNA3-hDVL2-PDZbinder335-mScarlet3 were generated after insertion of Xph334 and Xph335 sequences into pmScarlet3_C1 obtained from Addgene (#189753)^121^. Full-length human *Scribble*, cloned in pEGFP-C1, was generously provided by Prof. I. Macara (UVA, Charlottesville, VA, USA). Full-length rat *Shank3* cDNA has been described and used previously^98^. pCMV6-mTRIM46-myc-Flag was obtained from OriGene (#MR206757). pEGFP-C1-GFP-mMTCL1, pCAGGS-MCX-V5-mMTCL1 and its respective ΔN, ΔKR and ΔN/ΔKR contructs lacking MT-binding sites were a kind gift of Pr. A. Suzuki (Yokohama City University, Japan) ^88^. All plasmid DNA and shRNA constructs were sequence-verified by Eurofins Genomics.

#### Antibodies

The antibodies used in this study are listed below; working dilutions for each application are provided in the corresponding tables. Primary antibodies against Dishevelled proteins included rabbit polyclonal anti-Dvl2 antibodies from Millipore and Cell Signaling Technology, and a mouse monoclonal anti-Dvl3 antibody (clone 4D3; Santa Cruz Biotechnology). Antibodies from Synaptic Systems included guinea pig polyclonal and rabbit polyclonal anti-Ankyrin-G (AnkG), as well as guinea pig polyclonal and recombinant anti-TRIM46 antibodies. In addition, AnkG and Pan-Nav channels were detected using culture supernatants from HEK293T cells transfected with the corresponding antibody-expression plasmids (Addgene #128635 and #128623, respectively), as previously described by Andrews et al.^122^. For immunolabeling of cultured neurons, we also used a rabbit polyclonal anti-Prickle2 (Pk2) antibody^123^ kindly provided by D.Wu (NIDCD/NIH, USA), a rabbit polyclonal anti-Scrib antibody generated in-house (AbMM468), and a mouse monoclonal anti-acetylated tubulin antibody (clone 6-11B-1; Sigma-Aldrich). Epitope-tagged proteins were detected using chicken polyclonal anti-GFP (Abcam), rabbit polyclonal anti-GFP (Millipore), mouse monoclonal anti-GFP (clone JL-8; Takara), mouse monoclonal anti-HA (clone HA.11; BioLegend), rabbit polyclonal anti-HA (Sigma-Aldrich), mouse monoclonal anti-Myc (clone 9E10; Invitrogen), mouse monoclonal anti-V5 (clone SV5-Pk1; Invitrogen), chicken polyclonal anti-mCherry (EnCor Biotechnology), and rabbit polyclonal anti-DsRed (Living Colors; Takara). Fluorescent secondary antibodies conjugated to Alexa Fluor 488, 594, or 647 were obtained from Jackson ImmunoResearch, whereas Alexa Fluor 546-conjugated secondary antibodies were obtained from Invitrogen. For immunoblotting, horseradish peroxidase (HRP)-conjugated goat anti-mouse and goat anti-rabbit secondary antibodies were obtained from Jackson ImmunoResearch, and an HRP-conjugated mouse anti-rabbit secondary antibody was obtained from GE Healthcare.

#### Immunohistochemistry

Animals were deeply anesthetized by subcutaneous injection of xylazine (Pacman®, 20 mg/kg), followed by intraperitoneal administration of pentobarbital (Euthasol®, 800 mg/kg), and then transcardially perfused with phosphate buffer (PB) followed by 4% paraformaldehyde (PFA) in PB. Brains were removed and post-fixed in 4% PFA for 2 hrs. at 4°C. After post-fixation, brains were rinsed three times in PB and cryoprotected overnight in 20% sucrose in PB. Coronal sections (40–50 µm) were prepared using a VT1000S vibratome (Leica). Free-floating sections were permeabilized in 0.25–0.3% Triton X-100 in 1× PBS for 1 hr. at room temperature (RT) and blocked for 2 h at RT in 1× PBS containing 5% bovine serum albumin (BSA) and 5% normal goat serum (NGS). Primary antibodies were diluted in 1× PBS and incubated with the sections for either overnight or 48 h at 4°C, depending on the antibody. After washing, sections were incubated with fluorescently conjugated secondary antibodies diluted in 1× PBS for 1 h at RT. Sections were then mounted using ProLong Gold Antifade Mountant (Life Technologies).

#### Neuronal cell culture, transfection and immunofluorescence

Primary hippocampal neuron cultures were prepared on 15- or 18-mm glass coverslips coated with poly-L-lysine (PLL) and laminin. Coverslips were incubated with 30 µg/mL PLL in borate buffer for 2 h at 37°C, rinsed with sterile water (Braun), and subsequently coated overnight at 37°C with 2 µg/mL laminin diluted in water. The following day, coverslips were rinsed with sterile water and maintained in Neurobasal medium (Gibco) until cell plating. Primary hippocampal cultures were prepared from embryonic day 18.5 (E18.5) Sprague–Dawley rats. Hippocampi were dissected and collected in Hank’s Balanced Salt Solution (HBSS; Gibco) supplemented with HEPES and penicillin–streptomycin (Gibco), and subsequently rinsed in ice-cold chopping solution containing HBSS, HEPES, and DNase (Sigma-Aldrich). For enzymatic dissociation, hippocampi were incubated in pre-warmed chopping solution supplemented with trypsin (Gibco) for 12–15 min at 37°C with occasional gentle agitation. Trypsin activity was stopped by repeated washes with chopping solution, and the tissue was mechanically dissociated by gentle trituration using a Pasteur pipette. Cells were counted and plated in Neurobasal medium supplemented with B27 (50×; Gibco) and 2% fetal bovine serum (FBS). Four hours after plating, the medium was replaced with astrocyte-conditioned medium. Neurons were maintained at 37°C in a humidified atmosphere containing 5% CO₂. When required, neurons were transfected before plating using the Amaxa Rat Neuron Nucleofector Kit (Lonza) according to the manufacturer’s instructions. Nucleofection was performed using 5 × 10⁵ neurons per condition. For Dvl depletion experiments, cells were nucleofected with shDvl plasmid (8 µg), with pSuper used as the corresponding control plasmid (4 µg).

For experiments involving MTCL1, V5-MTCL1 was used at 1 µg. Xph334-mScarlet and Xph335-mScarlet were each used at 1.5 µg, with empty mScarlet plasmid (3 µg) used as the corresponding control. Following nucleofection, transfected neurons were mixed with 1 × 10⁵ non-transfected neurons. For immunocytochemistry, neurons were fixed at the indicated time points for 10 min at room temperature (RT) in 4% PFA containing 4% sucrose. For experiments requiring detergent extraction, 0.25% Triton X-100 was included in the fixation solution. Cells were subsequently permeabilized and blocked for 1 h at RT in PBS containing 5% BSA and 0.3% Triton X-100, followed by incubation with primary antibodies overnight at RT. After washing, cells were incubated with Alexa Fluor-conjugated secondary antibodies (Jackson ImmunoResearch) for 30 min at RT. Coverslips were mounted using Vectashield Vibrance Antifade Mounting Medium (Vector Laboratories). For U-ExM experiments, neurons were fixed for 10 min at RT in 4% PFA containing 4% sucrose and subsequently washed three times in PBS before further processing as described below.

#### Astrocytes conditioned medium

The day before astrocyte preparation, 75-cm² culture flasks were coated with poly-L-lysine (10 µg/mL) for 2 h at 37°C and rinsed with sterile water (B. Braun). Cortices from three embryonic day 18.5 (E18.5) Sprague–Dawley rats were pooled in Hank’s Balanced Salt Solution (HBSS; Gibco) supplemented with HEPES (Thermo Fisher Scientific), and rinsed in ice-cold chopping solution containing HBSS, HEPES, and DNase (Sigma-Aldrich). For enzymatic dissociation, the cortices were incubated in pre-warmed chopping solution containing trypsin (Thermo Fisher Scientific) for 10–15 min at 37°C with occasional gentle agitation. Trypsin activity was stopped by addition of blocking solution containing HBSS and HEPES, and the tissue was mechanically dissociated by gentle trituration with a Pasteur pipette. Cells were centrifuged for 10 min at 1,000 rpm at room temperature, and the pellet was resuspended in plating medium consisting of DMEM (Gibco) supplemented with 10% fetal bovine serum (FBS; Corning). After 1 week, astrocytes were detached with trypsin and replated into poly-L-lysine-coated culture dishes (10 µg/mL). Once confluent, the cultures were incubated for successive 3-day periods in Neurobasal medium supplemented with B27 (50×; Gibco) to generate astrocyte-conditioned medium. Astrocytes were maintained at 37°C in a humidified atmosphere containing 5% CO₂.

#### Ultrastructure expansion microscopy (U-ExM)

This protocol was adapted from the previously described ultrastructure expansion microscopy (U-ExM) procedure ^86^. Neurons were fixed at DIV7 as described above and stored in PBS at 4°C until processing. Samples were anchored by incubation for 3 h at room temperature (RT) in PBS containing 1% acrylamide (AA; Sigma-Aldrich) and 0.8% formaldehyde (FA; Sigma-Aldrich). After washing, gelation was performed for 30 min at 37°C in a solution containing 10% AA, 16.5% sodium acrylate, 0.1% N,N′-methylenebisacrylamide, 5% TEMED, and 5% ammonium persulfate (APS) in PBS. Coverslips were then incubated for 15 min at RT with agitation in denaturation buffer containing 50 mM Tris, 200 mM NaCl, and 200 mM SDS, pH 6.8, to facilitate detachment of the gel. The gels were subsequently transferred to fresh denaturation buffer and incubated for 30 min at 85°C with agitation. Initial expansion was achieved by three successive washes in Milli-Q water at RT. Once maximal expansion was reached, gel diameter was measured to determine the expansion factor. Coverslips were then incubated for 15 min at RT with agitation in denaturation buffer containing 50 mM Tris, 200 mM NaCl, and 200 mM SDS, pH 6.8, to facilitate detachment of the gel. The gels were subsequently transferred to fresh denaturation buffer and incubated for 30 min at 85°C with agitation. Initial expansion was achieved by three successive washes in Milli-Q water at RT. Once maximal expansion was reached, gel diameter was measured to determine the expansion factor.

For post-expansion immunostaining, gels were divided into pieces using a gel punch and incubated overnight at 37°C with agitation in PBS containing 2% bovine serum albumin (BSA), 0.03% sodium azide, and the appropriate primary antibodies. Gels were then washed four times for 5 min in PBS containing 0.1% Tween 20 and incubated for 2.5 h at 37°C with agitation in the same blocking solution containing fluorescent secondary antibodies and DAPI. After four additional washes, gels were stored in PBS at 4°C until imaging. Immediately before imaging, gels were re-expanded by two 10-min washes in Milli-Q water at RT and mounted between a microscope slide and coverslip without mounting medium.

#### Image acquisition and processing

Most imaging of brain sections was performed using a Zeiss AxioImager Z1 microscope equipped with a 63×/1.40 NA oil-immersion objective, an AxioCam MRm camera, ZEN software, and a Colibri 7 LED light source (Zeiss; excitation channels: UV 385/30 nm, V 423/44 nm, B 469/38 nm, C 511/44 nm, G 555/30 nm, Y 590/27 nm, and R 631/33 nm). Z-stacks spanning 9.6 µm were acquired as 40 optical sections with a step size of 0.24 µm. For dissociated neurons, Z-stacks comprising five optical sections were acquired with a step size of 0.25 µm, corresponding to a total depth of 1 µm. For U-ExM samples imaged using the Zeiss Apotome module, Z-stacks of 8–12 optical sections were acquired with a step size of 0.25 µm. Additional U-ExM imaging was performed using a Leica TCS SP8 confocal microscope. COS-7 cells were imaged as single optical sections. All multichannel images were acquired sequentially to minimize spectral crosstalk. Image processing and analysis were performed using Fiji/ImageJ (https://imagej.net/Fiji).

#### Image Quantification

Image analysis was performed in Fiji/ImageJ using a custom macro (developed by F. Cordelières, Bordeaux Imaging Centre) to automate sum-intensity projection and axon tracing with the Simple Neurite Tracer (SNT) plugin. Regions of interest (ROIs) measuring 20 pixels in width and 70–100 µm in length were extracted, and fluorescence intensity data were exported as XLS files, including distance and intensity values. Neurons were selected for analysis based on following criteria: (1) Having a clearly identifiable cell body (2) no overlapping neurites or AIS or artefacts on the AIS of interest (3) inclusion of Axon carrying dendrite with measurement beginning at the axon-dendrite intersection rather than the cell body (4) only one AIS measured per neuron. Data Processing and Analysis Fluorescence intensity profiles were processed using R and Python scripts (developed by M.Gonzalez, Institute of Condensed Matter Chemistry of Bordeaux; available at https://sourceforge.net/projects/isn33/files/ALSA/). Profiles were smoothed with a 5 µm sliding window and normalized between 1 (maximum smoothed fluorescence, corresponding to the AIS peak position) and 0 (minimum smoothed fluorescence)^124^. The axonal initial segment (AIS) start and end positions were determined at the proximal and distal axonal locations, respectively, where the normalized and smoothed profile of AnkG channel declined to 0.35. These positions were then mapped back to the raw XLS files to calculate the mean fluorescence intensity per neuron. AIS Fragmentation Analysis AIS fragmentation was assessed using a 5 µm sliding window on normalized profiles. The number of AIS fragments exceeding the 0.35 threshold was counted, with one fragment classified as a normal AIS and more than one fragment as a fragmented AIS. The distance between the start of the first fragment and the end of the last fragment was measured to quantify fragmentation extent. The TRIM46 labeling U-ExM images, in the converted form of the apotome, were processed in Fiji®. A line outside the neuron was drawn to convert it into a “plot profile” in order to determine the background noise threshold. Then a second line was drawn along the axis perpendicular to the AIS in its first third. This line was converted into a “plot profile,” and after subtracting the background noise, the signal was characterized by the number of fluorescence intensity peaks in the TRIM46 signal; at the same time, the width of the GFP signal was measured, providing information on the thickness of the AIS.

#### STORM imaging

Stochastic Optical Reconstruction Microscopy (STORM) imaging was performed at the Bordeaux Imaging Center (BIC UAR 3420) using the Leica SR GSD microscope (Leica Microsystems, Mannheim, Germany). DIV21 neurons cultured on 18-mm glass coverslips were processed for immunofluorescence as described above and mounted in Ludin chambers immediately before acquisition. For alignment and drift correction, coverslips were incubated with fluorescent fiducial beads (Life Technologies Tetraspeck microspheres 0.1µm T7279) diluted in 1× PBS for 10–15 min at room temperature, washed three times in 1× PBS, and finally mounted in freshly prepared STORM imaging buffer. The STORM imaging buffer consisted of a Tris/NaCl-based buffer supplemented with glucose, β-mercaptoethylamine/MEA, and an oxygen-scavenging enzymatic system containing pyranose oxidase and catalase. The pH of the buffer was verified before use and maintained between 7.5 and 8.0 to ensure efficient fluorophore photoswitching. The imaging chamber was carefully closed to avoid air bubbles and to preserve stable imaging conditions during acquisition. Image acquisition was performed using TIRF illumination previously aligned using fluorescent beads. Low-resolution TIRF images were first acquired to identify neurons of interest, assess staining quality, and verify the presence of fiducial beads within or near the region of interest. Single-molecule sequential acquisitions were performed in STORM mode, primarily using the far-red channel for Alexa Fluor 647, then using the orange channel for Alexa Fluor 532. The 642 nm laser and 532 nm laser respectively were used for fluorophore photoswitching and single-molecule detection, with laser power progressively adjusted to obtain a sparse and stable blinking regime. For each neuron, low-resolution reference images were acquired before each STORM sequence. Then, sequential image sequences were acquired under TIRF illumination, with acquisition parameters adjusted to ensure optimal single-molecule density while minimizing photobleaching. Raw data were saved in Leica-compatible image formats and subsequently processed for single-molecule localization and image reconstruction. Data were analyzed using PALMTracer, a MetaMorph (Molecular Devices, Sunnyvale, USA) plug-in developed at the Interdisciplinary Institute for Neuroscience (IINS, UMR5297, CNRS / University of Bordeaux)^125^. Briefly, single-molecule localization was performed using a combination of wavelet-based segmentation and Gaussian fitting, drift correction and channel alignment was performed using fiducial markers ^126^. Measurement of Manders coefficient to evaluate co-localization was performed directly on molecules coordinates using a parameter-free tessellation based analysis ^127^.

#### HEK293T and COS-7 culture, transfection and immunofluorescence

Human embryonic kidney 293T cells expressing the SV40 large T antigen (HEK293T; ATCC CRL-11268-G1™) and COS-7 cells (ATCC CRL-1651™) were maintained in Advanced Dulbecco’s modified Eagle’s medium (DMEM; Gibco) supplemented with 10% fetal bovine serum (FBS; Gibco), 1% L-glutamine (200 mM; Gibco), and 1% penicillin–streptomycin (10,000 U/mL; Gibco). Cells were cultured at 37°C in a humidified atmosphere containing 5% CO₂ and passaged according to the supplier’s recommendations. For HEK293T transfection, cells were seeded in 10-cm dishes and transfected at 30–40% confluence using linear polyethyleneimine (PEI; Polysciences, 1 mg/mL) at a ratio of 3 µL PEI per 1 µg plasmid DNA. Plasmids were diluted in 400 µL serum-free DMEM, mixed with PEI, vortexed, and incubated for 15 min at room temperature. The transfection mixture was then added dropwise to the cells. After 24 h, cells were harvested and processed for immunoprecipitation and pull-down assays as described below. For COS-7 transfection, cells were plated in six-well plates containing three 15-mm coverslips per well and transfected at 30–40% confluence using a calcium phosphate method, as previously described ^128^. For each transfection, plasmid DNA was mixed with 12.5 µL CaCl₂ and adjusted to a final volume of 100 µL with water. Unless otherwise indicated, 1 µg of each plasmid was used, whereas GFP-MTCL1 was used at 0.5 µg. The DNA/CaCl₂ solution was added dropwise to 100 µL of 2× HEPES-buffered saline (HBS), incubated for 30 min at room temperature in the dark, and then applied to the cells. After 5 h, cells were washed three times with serum-free DMEM and incubated for 10 min before replacement with complete medium. Cells were fixed 24 h later and processed for immunocytochemistry. COS-7 cells were fixed for 10 min at room temperature in 4% PFA containing 4% sucrose and permeabilized for 10 min in 0.25% Triton X-100 in PBS. Cells were blocked for 30 min at room temperature in PBS containing 5% bovine serum albumin (BSA; Euromedex) and 0.05% Tween 20 (Sigma-Aldrich). Primary antibodies were diluted in PBS and incubated for 1 h at room temperature. After washing, cells were incubated for 45 min at room temperature with fluorescent secondary antibodies diluted in PBS containing 1% BSA. Coverslips were mounted using Fluoromount-G antifade medium (Electron Microscopy Sciences).

#### Co-immunoprecipitation assays

HEK293T cells were seeded in 10-cm dishes and transfected at 50–60% confluence with GFP-MTCL1, pcDNA3.1-hDvl2-Myc-His, or both constructs using polyethylenimine (PEI) at an approximate DNA:PEI ratio of 1:3. Twenty-four hours after transfection, cells were washed with PBS, collected by scraping into ice-cold PBS, centrifuged at 1,500 rpm for 5 min at 4°C, and flash-frozen on dry ice. Cell pellets were lysed in ice-cold buffer containing 50 mM Tris-HCl, pH 7.5, 150 mM NaCl, 2 mM EDTA, and protease inhibitors. Lysates were homogenized using a Polytron, sonicated on ice, supplemented with 1% Triton X-100, and incubated for 3 h at 4°C. Input fractions were collected before clarification by ultracentrifugation at 84,035 × g for 40 min at 4°C. Clarified lysates were incubated overnight at 4°C with 20 µL GFP-Trap® Agarose beads (ChromoTek) pre-equilibrated in lysis buffer. Beads were then washed sequentially with buffers containing increasing NaCl concentrations from 50 to 150 mM. Bound proteins were eluted in 2× Laemmli sample buffer by heating for 5 min at 95°C, separated by 8% SDS-PAGE, transferred overnight at 4°C in transfer buffer containing 10% methanol, and analyzed by immunoblotting.

#### Whole-cell patch-clamp recordings/Electrophysiology

For whole-cell patch-clamps recordings we used nucleofected rat cultured hippocampal neurons between 8-10 days in vitro (control 9 ± 0.9 DIV, n = 14; shDvl 8.9 ± 0.8 DIV, n = 15). Single glass coverslips with adhering neurons were transferred into room temperature recording solution composed of (all in mM) 125 NaCl, 3 KCl, 25 HEPES, 2 CaCl_2_, 1 MgCl_2_ and 10 glucose. After placing the coverslip under an up-right microscope (LN-scope) neurons were continuously perfused with same solution at room temperature. Neurons were selected based on their expression of EGFP fluorescence and from each cover slip usually more than one neuron was sampled. The presented results are from four independent cultures and from each culture, control and shDvl knock-down neurons were recorded. Fluorescence exposure was limited in time and intensity to reduce phototoxicity. Whole-cell patch clamp recordings were performed with glass pipettes (BF150-75-10, Sutter Instruments) filled with intracellular solution that was composed of (in mM) 130 K-Gluconate, 10 KCl, 10 HEPES, 4 Mg2+-ATP, 0.3 Na+-GTP, 10 Na2-phosphocreatine, 15 biocytin and a pH of 7.25 (adjusted with KOH). All chemicals were purchased from Sigma-Aldrich. No liquid junction potential correction was applied to recordings. All experiments were controlled by a patch-clamp amplifier (Double IPA, Sutter Instruments, Novato, CA, USA) and SutterPatch software. Action potentials were sampled at 50 kHz and all other recordings were sampled with a minimum frequency of 10 kHz. A 5 kHz four-pole Bessel filter was applied to all recordings. Bridge balance and cell capacitance were compensated in current-clamp mode based on short negative current injections effectively reducing the voltage error. Analysis of recordings was automated in SutterPatch for membrane potential, input resistance, firing frequency and firing threshold. Input resistance was determined from small current injections (-5 pA to -10 pA) around the resting membrane potential in current clamp. Single action potentials were evoked by short 3 ms current injections and selected for analysis when the action potential was initiated shortly after the current injection. All action potential analysis was performed with Axograph (Axograph Scientific, NSW, Australia). Action potential threshold detection was set to 10 V/s and the amplitude was calculated between the absolute peak and the threshold. The AP half-width was determined from 50% of the maximum peak amplitude **(Fig. S7A)**. To determine the contribution of the axonal and somatic action potential, the first derivative (dV/dt) of the action potential was calculated in Axograph. The corresponding first and second peak of the derived AP voltage was used corresponding to the AIS AP (first peak) and the somatic AP (second peak, **(Fig. S7B)**). Post-hoc immunolabelling was performed from recorded and biocytin filled neurons. To recover the morphology, streptavidin coupled with Alexa555 (S21381, 1:500, Invitrogen) was used and combined with the immunolabelling protocol for cultured neurons to label AnkyrinG (see section). Neurons were re-identified based on live-fluorescence images of their somatic morphology. Post-hoc fluorescent images were acquired and for analysis the AnkG labeling was used to determine onset and length of the AIS.

#### Dvl2 PDZ domain binders selection and production

##### Plasmid construction

Plasmids were generated using either the In Vivo Assembly (IVA) method ^129^ or standard restriction cloning. The phagemid pSEX90 was derived from the commercial pSEX81 Surface Expression Phagemid Vector (PROGEN, Cat. No. PR3005) and modified as previously described^90^, with an additional BbvCI restriction site downstream of the pIII gene, unlike pSEX84. For bacterial production of the human Dvl2 PDZ domain (residues 264–353), the domain was subcloned into the pBIG vector to include an N-terminal biotin acceptor peptide tag, a deca-His tag, and a Tobacco Etch Virus (TEV) cleavage site^90^. For pepC1 peptide production, a bdSUMO domain (from pDG02583, a gift from Dirk Görlich, Addgene #104129)^130^ and the peptide sequence with a short linker were introduced into the pIG vector by IVA. For mammalian cell expression, Xph335 and Xph334 genes were subcloned into pmScarlet3_C1 (a gift from Dorus Gadella, Addgene #189753)^121^ using IVA. pcDNA3.1-PSD93-mGreenLantern was obtained by subcloning the PSD-93 isoform 2 rat sequence into pcDNA3.1-mGreenLantern (a gift from Gregory Petsko, Addgene plasmid # 161912)^131^ in frame with the fluorescent protein using BamHI and EcoRI as restriction sites.

##### Library construction

Libraries were constructed as previously described^90^. Single-stranded DNA (ssDNA) was obtained from E. coli CJ236 dut-ung-strain (TaKaRa, Cat. No. E4141S). Combinatorial libraries diversifying the BC and FG loops of the 10Fn3 scaffold were generated using pFunkel mutagenesis^132^, inspired by Kunkel mutagenesis. Diversity was introduced using phosphorylated oligonucleotides encoding randomized NNK codons: 4–5 NNK codons at position 27, 1 NNK codon at position 29, and 12–13 NNK codons at position 75, corresponding to the BC and FG loops, respectively.

##### Biopanning procedure

Phage libraries were produced by inoculating an aliquot of the frozen bacterial library into 300 ml of 2×YT medium supplemented with carbenicillin (50 µg/ml). During exponential growth (OD₆₀₀ = 0.4– 0.6), cells were infected with Hyperphage M13KO7ΔpIII helper phage (Progen, Cat No. PRHYPE) at a multiplicity of infection (MOI) of 20 and incubated for 1 h at 37 °C. IPTG (2.5 µM) and kanamycin (30 µg/ml) were added, and cultures were grown overnight at 37 °C with shaking (200 rpm). Cultures were centrifuged (16,000 × g, 10 min, 4 °C), and phage particles were precipitated by adding one-fifth volume of PEG/NaCl solution, followed by incubation for 5 min at room temperature and centrifugation (16,000 × g, 10 min, 4 °C). Phage pellets were resuspended in PBT2 buffer (50 mM Na₂HPO₄, 50 mM NaH₂PO₄, 150 mM NaCl, 0.2% Tween-20, 1% BSA, pH 7.4) at 1.4 ml per selection target. Insoluble material was removed by centrifugation (20,000 × g, 5 min, 4 °C). Biopanning was performed using a KingFisher Duo robotic platform. Streptavidin-coated magnetic beads (Dynabeads M-280, Thermo Fisher Scientific) were equilibrated in PT2 buffer (50 mM Na₂HPO₄, 50 mM NaH₂PO₄, 150 mM NaCl, 0.2% Tween-20, pH 7.4) and incubated for 10 min at room temperature with the biotinylated target. Beads were blocked with 5 µM biotin in PBT2 buffer for 15 min, washed, and then incubated with the phage solution for 15 min at room temperature with shaking. Beads were washed five times with 1 ml PT2 buffer, followed by a final wash with PBS, and recovered in 100 µl PBS. Bound phages were eluted with 100 µl glycine (100 mM, pH 2.7) for 10 min with shaking, and the eluate was neutralized with 50 µl NaHCO₃ (100 mM, pH 9.0). Three rounds of panning were performed with decreasing concentrations of the biotinylated fusion target. Neutralized phage eluates were used to infect E. coli XL1-Blue cells (OD₆₀₀ = 0.4–0.6) for 1 h at 37 °C with shaking. Aliquots were plated on 2×YT agar supplemented with carbenicillin (50 µg/ml) and glucose (10 mM) to estimate phage titers and recover individual clones. The remaining culture was supplemented with helper phage (1 × 10¹⁰ phage/ml) and incubated for 1 h at 37 °C, then transferred to 25 mL of 2×YT medium containing carbenicillin (50 µg/ml), kanamycin (25 µg/ml), and IPTG (2.5 µM), and grown overnight at 37 °C with shaking (200 rpm). Phages were precipitated with PEG/NaCl as described above and resuspended in 1.4 ml PBT2 buffer. Individual colonies were picked for downstream phage-ELISA screening.

##### Phage-ELISA

Phage-ELISAs were performed as previously described^90^. Briefly, 96 clones from the final panning round were grown in 2×YT medium supplemented with carbenicillin and M13KO7 helper phage, and phage-containing supernatants were collected for screening. Streptavidin-coated plates were incubated with biotinylated targets and blocked prior to the addition of phage supernatants. Bound phages were detected using an HRP-conjugated anti-M13 monoclonal antibody (SinoBiological, Cat. No. 11973-MM05T-H, diluted 1:5,000 in PBT) and TMB substrate, and absorbance was measured at 450 nm.

##### Protein production

Proteins were expressed and purified as previously described^90^. His-tagged proteins were produced in E. coli BL21 CodonPlus (DE3)-RIPL competent cells (Agilent, Cat. No. 230280) using auto-induction protocols^133^ at 16 °C for 20 h. Proteins were first isolated under native conditions by IMAC using Ni-charged resins and further purified by size-exclusion chromatography (SEC). Purified proteins were concentrated, aliquoted, and flash-frozen in liquid nitrogen for storage at -80 °C. For biotinylated protein production, homemade E. coli BL21 (DE3) competent cells (Thermo Fisher Scientific, Cat. No. C600003) containing the pACYC-mCh-BirA plasmid were transformed with the target gene in the pbIG vector. Transformed bacteria were selected on LB plates containing chloramphenicol and kanamycin to maintain both plasmids. 5 ml LB starter cultures with appropriate antibiotics were inoculated into 300 ml ZYM-5052 auto-inducing medium containing kanamycin (30 µg/ml) and chloramphenicol (34.5 µg/ml). Cultures were grown in sterile glass vessels for 5 h at 37 °C (250 rpm). Biotin (50 µM) was added, and the temperature was lowered to 16 °C for target protein expression (∼20 h). Proteins were purified as described for His-tagged proteins. Biotinylation yields were evaluated by SDS-PAGE (Sorenson et al., 2015). Data analysis and graphical representation were performed using GraphPad Prism version 10.

## Supporting information

Belmonte et al. Supplemental figures

## Data and statistical analysis

Data analysis and graphical representation were performed using GraphPad Prism version 10 (GraphPad Software, Boston, MA, USA). Details of data presentation, sample sizes, and statistical tests are provided in Table 1. Data distributions were assessed for normality using either the D’Agostino–Pearson or Shapiro–Wilk test. Statistical comparisons were then performed using the appropriate parametric or non-parametric test, with variance equality considered where applicable. Biological replicates are referred to as independent experiments. Statistical significance was defined as *P < 0.05, **P < 0.01, ***P < 0.001, and ****P < 0.0001; ns indicates no significant difference.

## LEGENDS (Supplementary Figures)

**Figure S1: Dvl2 is enriched at the AIS in the mouse cortex and DG.** (A) Illustrations of coronal section of brains at P7 immunolabeled for Dvl2 (magenta) and AnkG (green) in the mouse cortex. Dvl2 is enriched in the proximal part of the AIS (bracket, see high magnifications). (B-C) Illustrations of coronal section of brains at P21 immunolabeled for Dvl2 (magenta) and AnkG (green) in the mouse cortex (A) and mouse hippocampus of the DG (B). Dvl2 is enriched in the proximal part of the AIS (bracket, see high magnifications). Dotted circles outline the cell body and bracket the AIS. Scale bars: 10 µm in A to E.

**Figure S2 Dvl2 localizes proximally to AnkG, including in axon-carrying dendrite (AcD) neurons in vitro.** (A-C) DIV3 (A), DIV4 (B) and DIV10 (C) cultured hippocampal neurons in vitro immunolabeled for Dvl2 (magenta), AnkG (green). Linescans of normalized fluorescence intensity at the AIS of Dvl2 (magenta) and AnkG (green) at different timepoints indicate that Dvl2 is getting enriched at the proximal part of the AIS (see inset, magenta arrowheads) and colocalizes partially with AnkG (bracket and see inset, green arrowheads) from DIV4. Values are normalized to the maximum intensity of each marker and displayed with ± SEM. (D) DIV7 hippocampal axon-carrying dendrite (AcD) neurons in vitro immunolabeled for Dvl2 (magenta), AnkG (green). Scale bars : 20 µm in A, 10 µm in B and C.

**Figure S3: Characterization of Dvl2 and Dvl3 antibodies in neurons.** (**A**) Schematic of the mouse full-length Dvl2 protein with the different domains and the corresponding amino acid numbers. Dvl2 antibody (Rb #1) recognizes the sequence between 600 and 614aa (red). The specificity of the Dvl2 antibody is explained by the differences in the amino acids sequences of the 3 paralogues. In contrast, the homology in sequence between Mus Musculus, Homo Sapiens and Rattus norvegicus for Dvl2 explains the ability of the antibody to be used in different species. Dvl2 antibody (Rb #2) recognizes the sequence around position 638 (red). (**B**) Schematic of the mouse full-length Dvl3 protein with the different domains and the corresponding amino acid numbers. Dvl3 antibody recognizes the sequence between 607 and 704aa (red). (**C**) DIV7 hippocampal neurons in vitro immunolabeled for Dvl2 (magenta), Dvl3 (green) and AnkG (blue). Dvl2 staining partially overlaps with AnkG immunolabeling at AIS but there is no Dvl3 labelling. (**D**) DIV7 hippocampal neurons in vitro immunolabeled for Dvl2 (magenta) with the second antibody (Rb #2) and AnkG (green). Dvl2 staining partially overlaps with AnkG immunolabeling at AIS. Scale bar: 10 µm in C and D.

**Figure S4: Characterization of Dvl2 and Dvl3 antibodies in COS7 cells.** (**A**) Anti-HA blot confirms expression of HA-Dvl1, HA-Dvl2, and HA-Dvl3 in the corresponding transfected cells, with no signal in the WT cos-7 cells. Anti-GAPDH was used as loading control. (**B**) Anti-Dvl2 (Rb #1) detects a strong band at approximately 100 kDa, consistent with the migration of Dvl2 in western blot, which runs above its predicted molecular weight. Faint bands appear in lines 1, 2 and 3 corresponding to endogenous Dvl2 already present in Cos-7 cells. Anti-GAPDH was used as loading control. (**C**) Anti-Dvl2 (Rb #2) detects a strong band at approximately 100 kDa, consistent with the migration of Dvl2 in western blot, which runs above its predicted molecular weight. Anti-GAPDH was used as loading control. (**D**) Anti-Dvl3 detects a strong band at approximately 100 kDa, consistent with Dvl3 molecular weight when overexpressed in Cos-7 cells. Anti-GAPDH was used as loading control. (**E-G**) Cos7 cells transfected with HA-Dvl1, HA-Dvl2 or HA-Dvl3 immunolabeled with anti-Dvl2 (Rb #1) (E), anti-Dvl2 (Rb #2) (F) or anti-Dvl3 (G) in magenta and HA in green. Anti-Dvl2 (Rb #1) and anti-Dvl2 (Rb #2) labeling colocalize only with HA-Dvl2 respectively in (E) and (F) and anti-Dvl3 labeling colocalize only with HA-Dvl3 (G). Scale bar : 5 µm in E to G.

**Figure S5: Dishevelled regulates AIS formation (additional markers).** (A, B, C) Illustration of DIV7 neurons transfected with a control or shDvl plasmid and labelled for TRIM46 (A), Pk2 (B) or Scribble (Scrib) (C) (magenta) and GFP (green). Stars show the cell body and double arrows or arrowhead the AIS. Dvl-induced AIS fragmentation is illustrated as small, interspersed TRIM46+, Pk2+ or Scribble+ clusters shown by the arrowheads. (D, E, F) Quantifications of fluorescence intensity levels of TRIM46 (D), Pk2 (E) or Scribble (F) in shDvl neurons (excluding no AIS phenotypes) show downregulations in protein levels (****p<0.0001 or ***p<0.001 from Mann-Whitney’s test. TRIM46 (n=61 neurons CTRL, n=50 shDvl), Pk2 (n=33 neurons CTRL, n=40 shDvl) and Scrib (n=35 neurons CTRL, n=39 shDvl), from at least 3 independent experiments). Scalebars : 20 µm in (A, B, C).

**Figure S6: Dishevelled regulates AIS maintenance.** (A) Illustration of DIV17 neurons transfected at DIV13 with a control or shDvl plasmid and labelled for Dvl2 (magenta) and AnkG (blue). Stars show the cell body and double arrows the AIS. The shDvl-induced AIS fragmentation is illustrated as small, interspersed AnkG+ clusters as in (arrowheads) and Dvl2 labelling is decreased. (B) Quantification of fluorescence intensity level of Dvl2 in shDvl neurons at DIV17 (excluding no AIS phenotypes) show downregulation Dvl2 levels. (C) Percentages neurons per phenotype at DIV17 described as normal AIS, fragmented, very fragmented and loss of AIS with shDvl. Means ± SEM are shown. Scale bars : 20 µm in A.

**Figure S7: Dvl depletion lowers the current threshold for repetitive firing without altering maximal action potential frequency.** (A) Action potential analysis schematic for the most important parameters. (B) 1st derivative (dV/dt) of the action potential recorded at the soma showing the two components corresponding to the backpropagating AIS peak and the somatic peak. (C) Example recordings during long positive current injections show the generation of several action potentials in control and shDvl neurons. Several action potentials are elicited in each cell, however, firing patters differ probably as the neuronal populations were mixed hippocampal cells. (D) Population data of maximum AP frequency shows no difference between groups. (E) The shDvl neurons require much lower current injections to generate several action potentials consistent with the measured higher input resistance.

**Figure S8: MTCL1 MT-binding sites are required for Dvl2 recruitment to MT.** (**A**) COS7 cells transfected for 24 hrs with HA-Dvl2 either with V5-MTCL1 or MTCL1 deletion mutants for the KR, N, and both KR and N domains. **(B)** Quantification of the specific HA-Dvl2 recruitment to MTCL1+ MTs. HA-Dvl2 colocalizes with V5-MTCL1DKR which is less efficiently localized on MTs, but HA-Dvl2 does not colocalize with V5-MTCL1DN or V5-MTCL1DKRDN. Scale bar : 5 µm in A

**Figure S9: Xph334 and Xph335 do not recognize other PDZ-domain proteins like Scribble, PSD93 and Shank3.** COS7 cells were co-transfected with GFP-Scrib (A), PSD93-mGL (B), or GFP-Shank3 (C) (magenta) and either mScarlet, Xph334 or Xph335 (green). Merged images show no overlap between magenta and green signals, confirming absence of colocalization. Quantification of colocalization using Manders’ coefficient confirms the lack of colocalization (D–F). Scale bar : 5 µm in A to C

## ACKNOWLEDGEMENTS

We thank Pr. M.Bienz, Dr. C. Duplaa, Pr. S. Sokol, Pr. I. Macara, Pr. D. Görlich and Pr. A. Suzuki for kindly sharing cDNA constructs. We thank D. Wu (NIDCD/NIH, USA) for supplying Prickle2 antibody. We thank A. Martin, S. Seshadri, I. Seynat and L.Badruna for technical assistance, and J. Dompierre (IBGC, Bordeaux) for guidance with U-ExM. We thank the personnel of the Animal Facility of the Neurocentre Magendie for the animal care. Confocal and STORM imaging were done at the Bordeaux Imaging Center (BIC), a service unit of the CNRS-INSERM and Bordeaux University, member of the national infrastructure France BioImaging supported by the French National Research Agency (ANR-24-INBS-0005 FBI BIOGEN). The help of F.Cordelières (BIC) for the AIS macro for ImageJ is acknowledged. We thank M.Gonzalez (ICMCB, Bordeaux) for development and assistance with the fluorescence-profile analysis scripts. We thank M. Rasband (Baylor College of Medicine, Houston, USA) and A. Freal (Amsterdam UMC, Netherlands) for their constructive and insightful comments.

## FUNDING

This work was supported by INSERM to M.Mont. and N.S. and the French National Research Agency (ANR-23-CE16-0019-01 “NeupoComp” to M.S., A.B., M.Mont. and J.E.). A.B. was supported by the Region Aquitaine (2018.599) and the University of Bordeaux. J.V. was supported by a grant from « La Maison du Cerveau », Bordeaux. Our teams and the Biochemistry and Biophysics Platform (BioProt) are all part of the of the Bordeaux Neurocampus and supported by the GPR Brain_2030 “Nanocoding” project. The funding bodies had no influence in the design, data collection or interpretation of the experiments.

## DECLARATION OF GENERATIVE AI USE IN THE WRITING PROCESS

During the preparation of this manuscript, the authors used ChatGPT (OpenAI) to assist with language editing, reformulation, and improvement of the clarity and readability of selected sections of the text. All scientific interpretations, analyses, conclusions, and final wording were critically reviewed, edited, and validated by the authors, who take full responsibility for the content of the manuscript.

## AUTHOR CONTRIBUTIONS (CRediT taxonomy)

Conceptualization, M.Mont., M.S., A.B., N.S. and J.E.; Data curation, S.B., S.P.-C., M.Mont., M.S., A.B. and J.E.; Formal analysis, S.B., S.P.-C., J.V., C.I., A.D.-R., M.Don., E.C., E.R., L.V., M.Dav., M.Mo., M.S., A.B. and J.E.; Funding acquisition, M.Mont., M.S., A.B., N.S. and J.E.; Investigation, S.B., S.P.-C., J.V., C.I., A.D.-R., M.Don., E.C., E.R., L.V., N.A., M.M.M., M.Dav., M.Mo., M.Mont., M.S., A.B. and J.E.; Methodology, S.B., S.P.-C., J.V., C.I., M.Mont., M.S., A.B., N.S. and J.E.; Project administration, M.Mont., N.S. and J.E.; Resources, S.B., S.P.-C., J.V., C.I., M.Dav., M.Mo. and M.S.; Software, S.B.; Supervision, S.P.-C., M.Mont., M.S., A.B., N.S. and J.E.; Validation, S.B., S.P.-C., M.Mont., M.S., A.B., N.S. and J.E.; Visualization, S.B., S.P.-C., J.V., C.I., A.D.-R., M.Don., L.V., N.A., M.Mo., M.Mont., M.S., A.B. and J.E.; Writing – original draft, S.B., S.P.-C., M.S., A.B. and J.E.; Writing – review & editing, S.B., S.P.-C., C.I., A.D.-R., M.S., A.B., N.S. and J.E.

## DECLARATION OF INTERESTS

The authors declare no competing interests

---

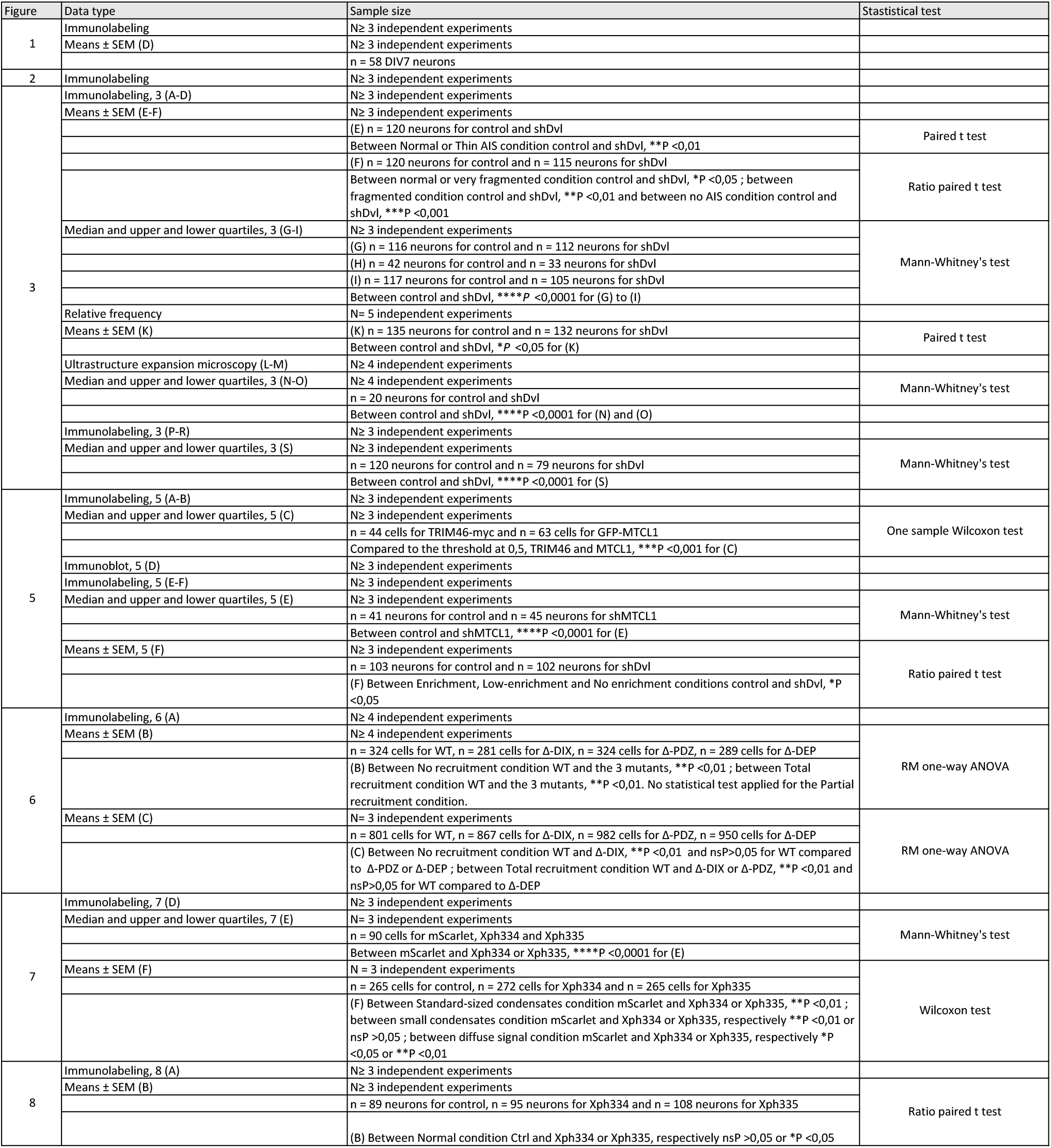

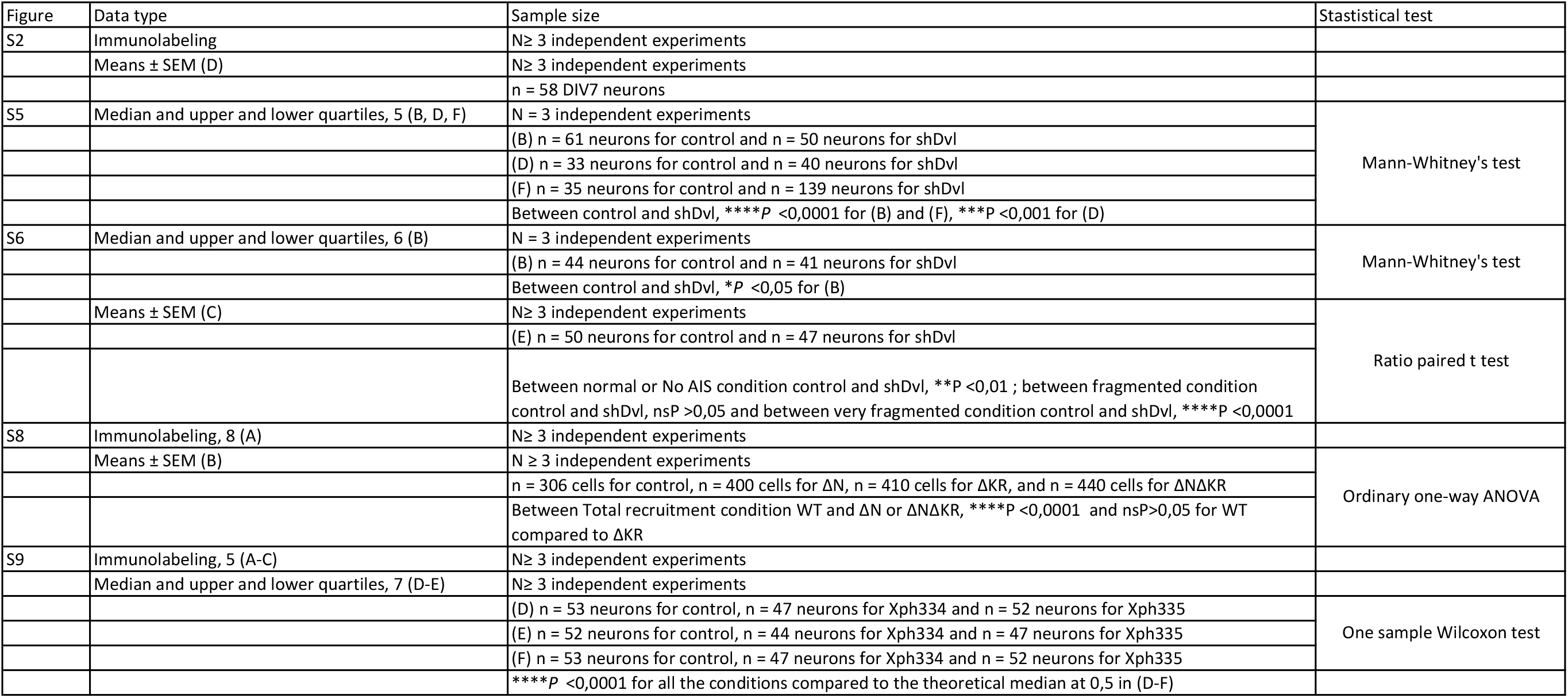

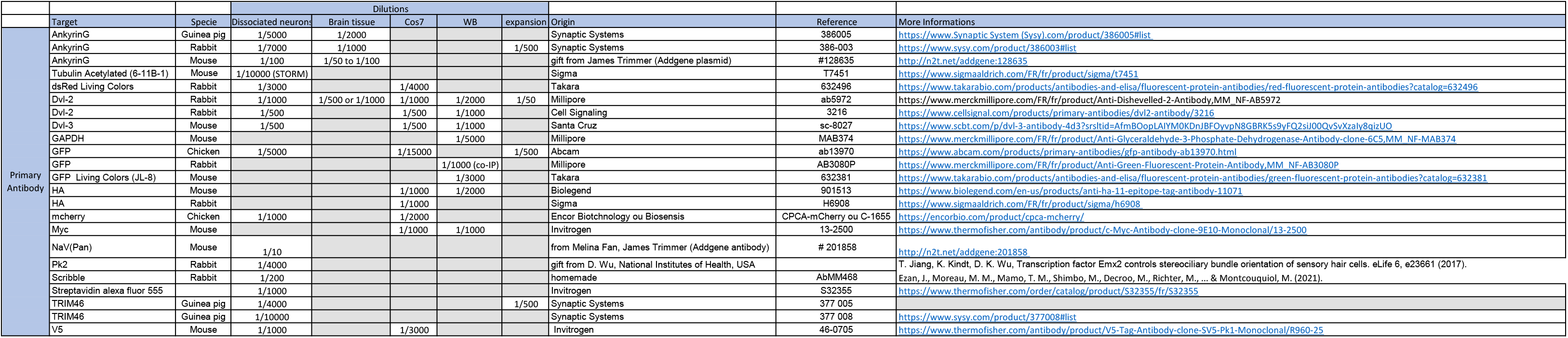

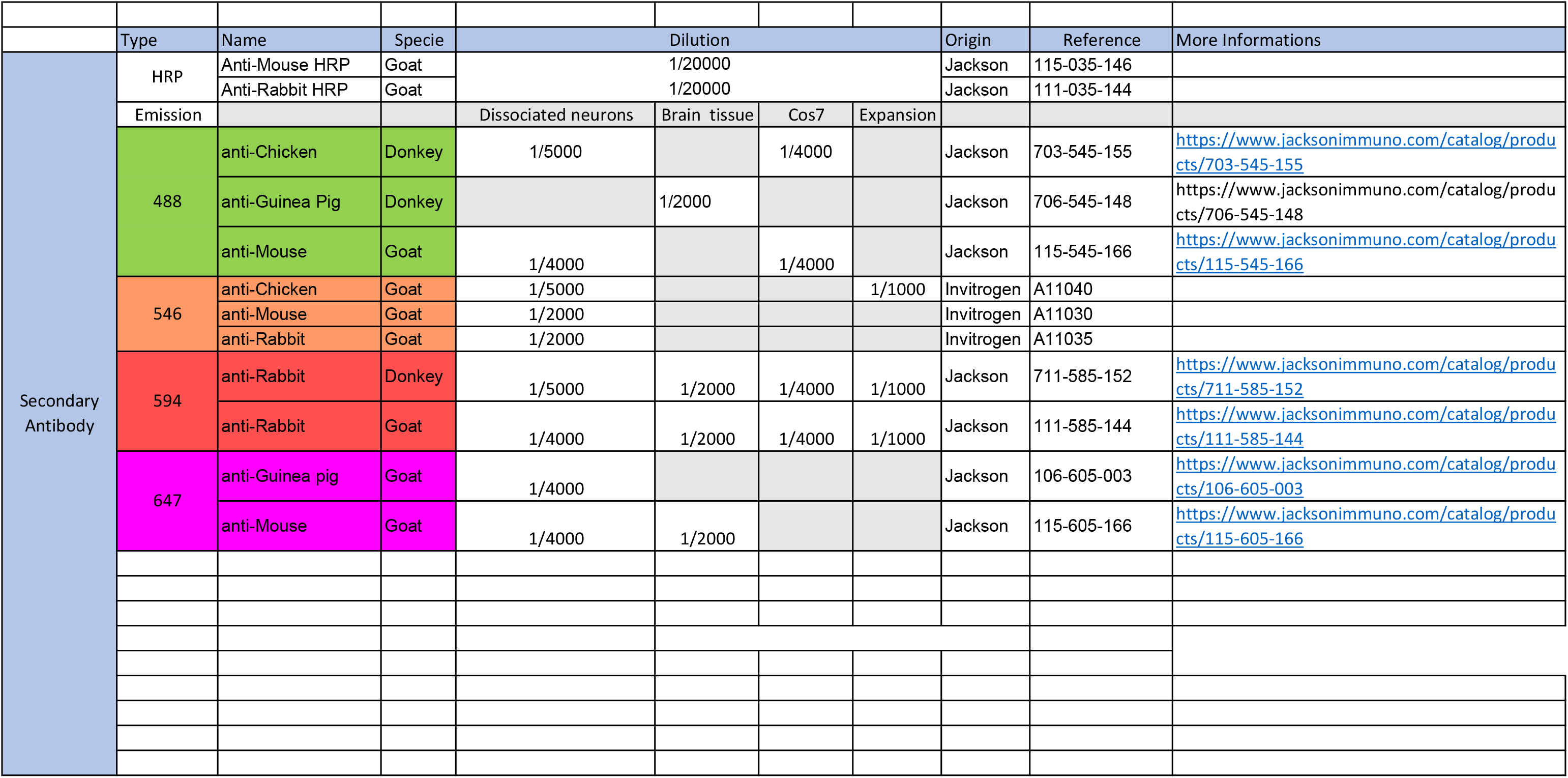

## Notes

### Competing Interest Statement

The authors have declared no competing interest.

## REFERENCES

1. Schelski, M. & Bradke, F. Neuronal polarization: From spatiotemporal signaling to cytoskeletal dynamics. Mol. Cell. Neurosci. 84, 11–28 (2017).

2. Yogev, S. & Shen, K. Establishing Neuronal Polarity with Environmental and Intrinsic Mechanisms. Neuron 96, 638–650 (2017).

3. Leterrier, C. The Axon Initial Segment: An Updated Viewpoint. J. Neurosci. Off. J. Soc. Neurosci. 38, 2135–2145 (2018).

4. Huang, C. Y.-M. & Rasband, M. N. Axon initial segments: structure, function, and disease. Ann. N. Y. Acad. Sci. 1420, 46–61 (2018).

5. Jenkins, P. M. & Bender, K. J. Axon initial segment structure and function in health and disease. Physiol. Rev. 105, 765–801 (2025).

6. Fréal, A. & Hoogenraad, C. C. The dynamic axon initial segment: From neuronal polarity to network homeostasis. Neuron 113, 649–669 (2025).

7. Kole, M. H. P. et al. Action potential generation requires a high sodium channel density in the axon initial segment. Nat. Neurosci. 11, 178–186 (2008).

8. Battefeld, A., Tran, B. T., Gavrilis, J., Cooper, E. C. & Kole, M. H. P. Heteromeric Kv7.2/7.3 channels differentially regulate action potential initiation and conduction in neocortical myelinated axons. J. Neurosci. Off. J. Soc. Neurosci. 34, 3719–3732 (2014).

9. Kole, M. H. & Brette, R. The electrical significance of axon location diversity. Curr. Opin. Neurobiol. 51, 52–59 (2018).

10. Buffington, S. A. & Rasband, M. N. The axon initial segment in nervous system disease and injury. Eur. J. Neurosci. 34, 1609–1619 (2011).

11. Fujitani, M., Otani, Y. & Miyajima, H. Pathophysiological Roles of Abnormal Axon Initial Segments in Neurodevelopmental Disorders. Cells 10, 2110 (2021).

12. Garrido, J. J. Contribution of Axon Initial Segment Structure and Channels to Brain Pathology. Cells 12, 1210 (2023).

13. Song, D.-Y., Yuan, L., Yang, W., Li, W. & Li, J.-Y. Axon Initial Segment: Structure, Biological Functions, Diseases, and Therapeutic Targets. MedComm 7, e70689 (2026).

14. Fréal, A. et al. Feedback-Driven Assembly of the Axon Initial Segment. Neuron 104, 305–321.e8 (2019).

15. Kapitein, L. C. & Hoogenraad, C. C. Building the Neuronal Microtubule Cytoskeleton. Neuron 87, 492–506 (2015).

16. Zhong, G. et al. Developmental mechanism of the periodic membrane skeleton in axons. eLife 3, e04581 (2014).

17. Leterrier, C. et al. Nanoscale Architecture of the Axon Initial Segment Reveals an Organized and Robust Scaffold. Cell Rep. 13, 2781–2793 (2015).

18. Winckler, B., Forscher, P. & Mellman, I. A diffusion barrier maintains distribution of membrane proteins in polarized neurons. Nature 397, 698–701 (1999).

19. Nakada, C. et al. Accumulation of anchored proteins forms membrane diffusion barriers during neuronal polarization. Nat. Cell Biol. 5, 626–632 (2003).

20. Jenkins, S. M. & Bennett, V. Ankyrin-G coordinates assembly of the spectrin-based membrane skeleton, voltage-gated sodium channels, and L1 CAMs at Purkinje neuron initial segments. J. Cell Biol. 155, 739–746 (2001).

21. Komada, M. & Soriano, P. [Beta]IV-spectrin regulates sodium channel clustering through ankyrin-G at axon initial segments and nodes of Ranvier. J. Cell Biol. 156, 337–348 (2002).

22. Hedstrom, K. L., Ogawa, Y. & Rasband, M. N. AnkyrinG is required for maintenance of the axon initial segment and neuronal polarity. J. Cell Biol. 183, 635–640 (2008).

23. van Beuningen, S. F. B. et al. TRIM46 Controls Neuronal Polarity and Axon Specification by Driving the Formation of Parallel Microtubule Arrays. Neuron 88, 1208–1226 (2015).

24. Harterink, M. et al. TRIM46 Organizes Microtubule Fasciculation in the Axon Initial Segment. J. Neurosci. Off. J. Soc. Neurosci. 39, 4864–4873 (2019).

25. Satake, T. et al. MTCL1 plays an essential role in maintaining Purkinje neuron axon initial segment. EMBO J. 36, 1227–1242 (2017).

26. Simons, M. & Mlodzik, M. Planar cell polarity signaling: from fly development to human disease. Annu. Rev. Genet. 42, 517–540 (2008).

27. Vladar, E. K., Antic, D. & Axelrod, J. D. Planar cell polarity signaling: the developing cell’s compass. Cold Spring Harb. Perspect. Biol. 1, a002964 (2009).

28. Wallingford, J. B. Planar cell polarity and the developmental control of cell behavior in vertebrate embryos. Annu. Rev. Cell Dev. Biol. 28, 627–653 (2012).

29. Ezan, J. & Montcouquiol, M. Revisiting planar cell polarity in the inner ear. Semin. Cell Dev. Biol. 24, 499–506 (2013).

30. Goodrich, L. V. & Strutt, D. Principles of planar polarity in animal development. Development 138, 1877–1892 (2011).

31. Singh, J. & Mlodzik, M. Planar cell polarity signaling: coordination of cellular orientation across tissues. Wiley Interdiscip. Rev. Dev. Biol. 1, 479–499 (2012).

32. Butler, M. T. & Wallingford, J. B. Planar cell polarity in development and disease. Nat. Rev. Mol. Cell Biol. 18, 375–388 (2017).

33. Wang, Y. & Nathans, J. Tissue/planar cell polarity in vertebrates: new insights and new questions. Development 134, 647–658 (2007).

34. Devenport, D. The cell biology of planar cell polarity. J. Cell Biol. 207, 171–179 (2014).

35. Davey, C. F. & Moens, C. B. Planar cell polarity in moving cells: think globally, act locally. Development 144, 187–200 (2017).

36. Koca, Y., Collu, G. M. & Mlodzik, M. Wnt-frizzled planar cell polarity signaling in the regulation of cell motility. Curr. Top. Dev. Biol. 150, 255–297 (2022).

37. Zou, Y. Wnt signaling in axon guidance. Trends Neurosci. 27, 528–532 (2004).

38. Ezan, J. et al. Early loss of Scribble affects cortical development, interhemispheric connectivity and psychomotor activity. Sci. Rep. 11, 9106 (2021).

39. Ezan, J. et al. Neuron-Specific Deletion of Scrib in Mice Leads to Neuroanatomical and Locomotor Deficits. Front. Genet. 13, 872700 (2022).

40. Feng, B. et al. Planar cell polarity signaling components are a direct target of β-amyloid-associated degeneration of glutamatergic synapses. Sci. Adv. 7, eabh2307 (2021).

41. Freitas, A. E., Gorodetski, L., Lim, W. L. & Zou, Y. Emerging roles of planar cell polarity proteins in glutamatergic synapse formation, maintenance and function in health and disease. Dev. Dyn. Off. Publ. Am. Assoc. Anat. 252, 1068–1076 (2023).

42. Jarjour, A. A. et al. The formation of paranodal spirals at the ends of CNS myelin sheaths requires the planar polarity protein Vangl2. Glia 68, 1840–1858 (2020).

43. Messéant, J. et al. Wnt proteins contribute to neuromuscular junction formation through distinct signaling pathways. Development 144, 1712–1724 (2017).

44. Robert, B. J. A. et al. Vangl2 in the Dentate Network Modulates Pattern Separation and Pattern Completion. Cell Rep. 31, 107743 (2020).

45. Depret, N. et al. The correct connectivity of the DG-CA3 circuits involved in declarative memory processes depends on Vangl2-dependent planar cell polarity signaling. Prog. Neurobiol. 246, 102728 (2025).

46. Tissir, F., Bar, I., Jossin, Y., De Backer, O. & Goffinet, A. M. Protocadherin Celsr3 is crucial in axonal tract development. Nat. Neurosci. 8, 451–457 (2005).

47. Sowers, L. P. et al. Disruption of the non-canonical Wnt gene PRICKLE2 leads to autism-like behaviors with evidence for hippocampal synaptic dysfunction. Mol. Psychiatry 18, 1077–1089 (2013).

48. Sans, N., Ezan, J., Moreau, M. M. & Montcouquiol, M. Planar Cell Polarity Gene Mutations in Autism Spectrum Disorder, Intellectual Disabilities, and Related Deletion/Duplication Syndromes. in Neuronal and Synaptic Dysfunction in Autism Spectrum Disorder and Intellectual Disability 189–219 (Elsevier, 2016). doi:10.1016/B978-0-12-800109-7.00013-3.

49. Hakanen, J., Ruiz-Reig, N. & Tissir, F. Linking Cell Polarity to Cortical Development and Malformations. Front. Cell. Neurosci. 13, (2019).

50. Yang, Y., Mao, Y., Zhang, Y. & Xiong, T. Evolving Insights into Prickle2 in Neurodevelopment and Neurological Disorders. Mol. Neurobiol. 62, 8540–8550 (2025).

51. Li, H.-Y., Shen, Y., Vong, K. I., Kahle, K. T. & Gleeson, J. G. De novo mutations and environmental modifiers: lessons from neural tube defects. Trends Genet. TIG 42, 339–349 (2026).

52. Dorrego-Rivas, A. et al. The core PCP protein Prickle2 regulates axon number and AIS maturation by binding to AnkG and modulating microtubule bundling. Sci. Adv. 8, eabo6333 (2022).

53. Micinski, D. & Hotulainen, P. Actin polymerization and longitudinal actin fibers in axon initial segment plasticity. Front. Mol. Neurosci. 17, 1376997 (2024).

54. Zhang, W. et al. A hierarchy of PDZ domain scaffolding proteins clusters the Kv1 K+ channel protein complex at the axon initial segment. Sci. Adv. 11, eadv1281 (2025).

55. Gao, C. & Chen, Y.-G. Dishevelled: The hub of Wnt signaling. Cell. Signal. 22, 717–727 (2010).

56. Mlodzik, M. The Dishevelled Protein Family: Still Rather a Mystery After Over 20 Years of Molecular Studies. Curr. Top. Dev. Biol. 117, 75–91 (2016).

57. Shi, D.-L. Decoding Dishevelled-Mediated Wnt Signaling in Vertebrate Early Development. Front. Cell Dev. Biol. 8, 588370 (2020).

58. Wallingford, J. B. & Habas, R. The developmental biology of Dishevelled: an enigmatic protein governing cell fate and cell polarity. Dev. Camb. Engl. 132, 4421–4436 (2005).

59. Sharma, M., Castro-Piedras, I., Simmons, G. E. & Pruitt, K. Dishevelled: A masterful conductor of complex Wnt signals. Cell. Signal. 47, 52–64 (2018).

60. Ahmad-Annuar, A. et al. Signaling across the synapse: a role for Wnt and Dishevelled in presynaptic assembly and neurotransmitter release. J. Cell Biol. 174, 127–139 (2006).

61. Narvaes, R. F. & Furini, C. R. G. Role of Wnt signaling in synaptic plasticity and memory. Neurobiol. Learn. Mem. 187, 107558 (2022).

62. Belinson, H. et al. Prenatal β-catenin/Brn2/Tbr2 transcriptional cascade regulates adult social and stereotypic behaviors. Mol. Psychiatry 21, 1417–1433 (2016).

63. Mudher, A. et al. Dishevelled regulates the metabolism of amyloid precursor protein via protein kinase C/mitogen-activated protein kinase and c-Jun terminal kinase. J. Neurosci. Off. J. Soc. Neurosci. 21, 4987–4995 (2001).

64. Wang, L. et al. Structural and Functional Insights into Dishevelled-Mediated Wnt Signaling. Cells 13, 1870 (2024).

65. Bilic, J. et al. Wnt induces LRP6 signalosomes and promotes dishevelled-dependent LRP6 phosphorylation. Science 316, 1619–1622 (2007).

66. Schwarz-Romond, T. et al. The DIX domain of Dishevelled confers Wnt signaling by dynamic polymerization. Nat. Struct. Mol. Biol. 14, 484–492 (2007).

67. Bienz, M. Signalosome assembly by domains undergoing dynamic head-to-tail polymerization. Trends Biochem. Sci. 39, 487–495 (2014).

68. Sear, R. P. Dishevelled: a protein that functions in living cells by phase separating. Soft Matter 3, 680–684 (2007).

69. Kang, K., Shi, Q., Wang, X. & Chen, Y.-G. Dishevelled phase separation promotes Wnt signalosome assembly and destruction complex disassembly. J. Cell Biol. 221, e202205069 (2022).

70. Vamadevan, V., Chaudhary, N. & Maddika, S. Ubiquitin-assisted phase separation of dishevelled-2 promotes Wnt signalling. J. Cell Sci. 135, jcs260284 (2022).

71. Kravec, M. et al. Carboxy-terminal polyglutamylation regulates signaling and phase separation of the Dishevelled protein. EMBO J. 43, 5635–5666 (2024).

72. Wu, X., Cai, Q., Feng, Z. & Zhang, M. Liquid-Liquid Phase Separation in Neuronal Development and Synaptic Signaling. Dev. Cell 55, 18–29 (2020).

73. Bencsik, N., Kimsanaliev, D., Tárnok, K. & Schlett, K. Stress-Induced Membraneless Organelles in Neurons: Bridging Liquid-Liquid Phase Separation and Neurodevelopmental Dysfunction. Int. J. Mol. Sci. 26, 9068 (2025).

74. Ciani, L., Krylova, O., Smalley, M. J., Dale, T. C. & Salinas, P. C. A divergent canonical WNT-signaling pathway regulates microtubule dynamics: dishevelled signals locally to stabilize microtubules. J. Cell Biol. 164, 243–253 (2004).

75. Krylova, O., Messenger, M. J. & Salinas, P. C. Dishevelled-1 regulates microtubule stability: a new function mediated by glycogen synthase kinase-3beta. J. Cell Biol. 151, 83–94 (2000).

76. Zhang, X. et al. Dishevelled promotes axon differentiation by regulating atypical protein kinase C. Nat. Cell Biol. 9, 743–754 (2007).

77. Wynshaw-Boris, A. Dishevelled: in vivo roles of a multifunctional gene family during development. Curr. Top. Dev. Biol. 101, 213–235 (2012).

78. Metcalfe, C. et al. Dvl2 promotes intestinal length and neoplasia in the ApcMin mouse model for colorectal cancer. Cancer Res. 70, 6629–6638 (2010).

79. Pulvirenti, T. et al. Dishevelled 2 signaling promotes self-renewal and tumorigenicity in human gliomas. Cancer Res. 71, 7280–7290 (2011).

80. Thome, C. et al. Axon-carrying dendrites convey privileged synaptic input in hippocampal neurons. Neuron 83, 1418–1430 (2014).

81. Han, Y. et al. Unveiling the cell biology of hippocampal neurons with dendritic axon origin. J. Cell Biol. 224, e202403141 (2025).

82. Harnoš, J. et al. Dishevelled-3 conformation dynamics analyzed by FRET-based biosensors reveals a key role of casein kinase 1. Nat. Commun. 10, 1804 (2019).

83. Gegg, M. et al. Flattop regulates basal body docking and positioning in mono-and multiciliated cells. eLife 3, e03842 (2014).

84. Mahuzier, A. et al. Dishevelled stabilization by the ciliopathy protein Rpgrip1l is essential for planar cell polarity. J. Cell Biol. 198, 927–940 (2012).

85. Dotti, C. G., Sullivan, C. A. & Banker, G. A. The establishment of polarity by hippocampal neurons in culture. J. Neurosci. Off. J. Soc. Neurosci. 8, 1454–1468 (1988).

86. Gambarotto, D. et al. Imaging cellular ultrastructures using expansion microscopy (U-ExM). Nat. Methods 16, 71–74 (2019).

87. Dorrego-Rivas, A. et al. The core PCP protein Prickle2 regulates axon number and AIS maturation by binding to AnkG and modulating microtubule bundling. Sci. Adv. 8, eabo6333 (2022).

88. Sato, Y. et al. The novel PAR-1-binding protein MTCL1 has crucial roles in organizing microtubules in polarizing epithelial cells. J. Cell Sci. 126, 4671–4683 (2013).

89. Rimbault, C. et al. Engineering paralog-specific PSD-95 recombinant binders as minimally interfering multimodal probes for advanced imaging techniques. eLife 13, e69620 (2024).

90. Rimbault, C. et al. Engineering selective competitors for the discrimination of highly conserved protein-protein interaction modules. Nat. Commun. 10, 4521 (2019).

91. Sha, F., Salzman, G., Gupta, A. & Koide, S. Monobodies and other synthetic binding proteins for expanding protein science. Protein Sci. 26, 910–924 (2017).

92. Abramson, J. et al. Accurate structure prediction of biomolecular interactions with AlphaFold 3. Nature 630, 493–500 (2024).

93. Grillo-Bosch, D., Choquet, D. & Sainlos, M. Inhibition of PDZ domain-mediated interactions. Drug Discov. Today Technol. 10, e531–e540 (2013).

94. Mu, Y., Cai, P., Hu, S., Ma, S. & Gao, Y. Characterization of Diverse Internal Binding Specificities of PDZ Domains by Yeast Two-Hybrid Screening of a Special Peptide Library. PLoS ONE 9, e88286 (2014).

95. Ye, F. & Zhang, M. Structures and target recognition modes of PDZ domains: recurring themes and emerging pictures. Biochem. J. 455, 1–14 (2013).

96. Zhang, Y. et al. Inhibition of Wnt signaling by Dishevelled PDZ peptides. Nat. Chem. Biol. 5, 217–219 (2009).

97. Ogawa, Y. et al. Postsynaptic density-93 clusters Kv1 channels at axon initial segments independently of Caspr2. J. Neurosci. Off. J. Soc. Neurosci. 28, 5731–5739 (2008).

98. Durand, C. M. et al. SHANK3 mutations identified in autism lead to modification of dendritic spine morphology via an actin-dependent mechanism. Mol. Psychiatry 17, 71–84 (2012).

99. Zeng, M. et al. Reconstituted Postsynaptic Density as a Molecular Platform for Understanding Synapse Formation and Plasticity. Cell 174, 1172–1187.e16 (2018).

100. Ryan, V. H. & Fawzi, N. L. Physiological, Pathological, and Targetable Membraneless Organelles in Neurons. Trends Neurosci. 42, 693–708 (2019).

101. Hamdan, H. et al. Mapping axon initial segment structure and function by multiplexed proximity biotinylation. Nat. Commun. 11, 100 (2020).

102. Melton, A. J. et al. TRIM46 Is Required for Microtubule Fasciculation In Vivo But Not Axon Specification or Axon Initial Segment Formation. J. Neurosci. Off. J. Soc. Neurosci. 44, e0976242024 (2024).

103. Belmonte, S. Evaluating TRIM46’s Role in Axon Initial Segment Formation. J. Neurosci. Off. J. Soc. Neurosci. 45, e0337252025 (2025).

104. Kikuchi, K., Niikura, Y., Kitagawa, K. & Kikuchi, A. Dishevelled, a Wnt signalling component, is involved in mitotic progression in cooperation with Plk1. EMBO J. 29, 3470–3483 (2010).

105. Cervenka, I. et al. Dishevelled is a NEK2 kinase substrate controlling dynamics of centrosomal linker proteins. Proc. Natl. Acad. Sci. U. S. A. 113, 9304–9309 (2016).

106. Schubert, A. et al. Superresolution microscopy localizes endogenous Dvl2 to Wnt signaling-responsive biomolecular condensates. Proc. Natl. Acad. Sci. U. S. A. 119, e2122476119 (2022).

107. Shnitsar, I. et al. PTEN regulates cilia through Dishevelled. Nat. Commun. 6, 8388 (2015).

108. Kohoutová, Š. Microtubule-associated proteins as mediators of interactions between Dishevelled and microtubules. https://dspace.cuni.cz/handle/20.500.11956/204070 (2025).

109. Wiegand, T. & Hyman, A. A. Drops and fibers - how biomolecular condensates and cytoskeletal filaments influence each other. Emerg. Top. Life Sci. 4, 247–261 (2020).

110. Volkov, V. A. & Akhmanova, A. Phase separation on microtubules: from droplet formation to cellular function? Trends Cell Biol. 34, 18–30 (2024).

111. Duan, D. & Koleske, A. J. Phase separation of microtubule-binding proteins - implications for neuronal function and disease. J. Cell Sci. 137, jcs263470 (2024).

112. Hatos, A., Tosatto, S. C. E., Vendruscolo, M. & Fuxreiter, M. FuzDrop on AlphaFold: visualizing the sequence-dependent propensity of liquid-liquid phase separation and aggregation of proteins. Nucleic Acids Res. 50, W337–W344 (2022).

113. Patel, A. et al. Principles and functions of condensate modifying drugs. Front. Mol. Biosci. 9, 1007744 (2022).

114. Kuznetsova, K. et al. CD-CODE 2.0: an enhanced condensate knowledgebase integrating pathobiology, condensate modulating drugs, and host-pathogen interactions. Nucleic Acids Res. 54, D375–D382 (2026).

115. Mitrea, D. M., Mittasch, M., Gomes, B. F., Klein, I. A. & Murcko, M. A. Modulating biomolecular condensates: a novel approach to drug discovery. Nat. Rev. Drug Discov. 21, 841–862 (2022).

116. Fleming, C. & Schuijers, J. Targeting biomolecular condensates: beyond dissolution. BMC Biol. 24, 59 (2026).

117. Tapia, M. et al. GSK3 and β-catenin determines functional expression of sodium channels at the axon initial segment. Cell. Mol. Life Sci. CMLS 70, 105–120 (2013).

118. Caballero-Florán, R. N., Dean, K. P., Nelson, A. D., Min, L. & Jenkins, P. M. Lithium restores inhibitory function and neuronal excitability through GSK-3β inhibition in a bipolar disorder-associated Ank3 variant mouse model. Neuropharmacology 279, 110649 (2025).

119. McSwiggen, D. T., Mir, M., Darzacq, X. & Tjian, R. Evaluating phase separation in live cells: diagnosis, caveats, and functional consequences. Genes Dev. 33, 1619–1634 (2019).

120. Marchetto, M. C. et al. Altered proliferation and networks in neural cells derived from idiopathic autistic individuals. Mol. Psychiatry 22, 820–835 (2017).

121. Gadella, T. W. J. et al. mScarlet3: a brilliant and fast-maturing red fluorescent protein. Nat. Methods 20, 541–545 (2023).

122. Andrews, N. P. et al. A toolbox of IgG subclass-switched recombinant monoclonal antibodies for enhanced multiplex immunolabeling of brain. eLife 8, e43322 (2019).

123. Jiang, T., Kindt, K. & Wu, D. K. Transcription factor Emx2 controls stereociliary bundle orientation of sensory hair cells. eLife 6, e23661 (2017).

124. Grubb, M. S. & Burrone, J. Activity-dependent relocation of the axon initial segment fine-tunes neuronal excitability. Nature 465, 1070–1074 (2010).

125. Butler, C. et al. Multi-Dimensional Spectral Single Molecule Localization Microscopy. Front. Bioinforma. 2, 813494 (2022).

126. Kechkar, A., Nair, D., Heilemann, M., Choquet, D. & Sibarita, J.-B. Real-time analysis and visualization for single-molecule based super-resolution microscopy. PloS One 8, e62918 (2013).

127. Levet, F. & Sibarita, J.-B. PoCA: a software platform for point cloud data visualization and quantification. Nat. Methods 20, 629–630 (2023).

128. Piguel, N. H. et al. Scribble1/AP2 complex coordinates NMDA receptor endocytic recycling. Cell Rep. 9, 712–727 (2014).

129. García-Nafría, J., Watson, J. F. & Greger, I. H. IVA cloning: A single-tube universal cloning system exploiting bacterial In Vivo Assembly. Sci. Rep. 6, 27459 (2016).

130. Pleiner, T., Bates, M. & Görlich, D. A toolbox of anti–mouse and anti–rabbit IgG secondary nanobodies. J. Cell Biol. 217, 1143–1154 (2018).

131. Campbell, B. C. et al. mGreenLantern: a bright monomeric fluorescent protein with rapid expression and cell filling properties for neuronal imaging. Proc. Natl. Acad. Sci. U. S. A. 117, 30710–30721 (2020).

132. Firnberg, E. & Ostermeier, M. PFunkel: Efficient, Expansive, User-Defined Mutagenesis. PLoS ONE 7, e52031 (2012).

133. Studier, F. W. Protein production by auto-induction in high-density shaking cultures. Protein Expr. Purif. 41, 207–234 (2005).

