## Supplementary material for "Dishevelled2 Condensates control Axon Initial Segment (AIS) assembly and function through MTCL1-Dependent microtubule recruitment": Belmonte et al. Supplemental figures

*Belmonte et al. SUPPLEMENTARY FIGURES*

**Figure list:**

**Fig. 1.** Dvl2 is enriched at the AIS *in vivo* and *in vitro.*

**Fig.S1.** Dvl2 is enriched at the AIS in the mouse cortex and DG

**Fig. S2.** Dvl2 localizes proximally to AnkG, including in axon-carrying dendrite (AcD) neurons *in vitro*

**Fig. S3.** Characterization of Dvl2 and Dvl3 antibodies in neurons

**Fig. S4.** Characterization of Dvl2 and Dvl3 antibodies in Cos7 cells

**Fig. 2.** Dvl2 is an early bona fide MT-associated component of the AIS.

**Fig. 3.** Dishevelled regulates AIS formation.

**Fig. S5.** Dishevelled regulates AIS formation (additional markers).

**Fig. S6.** Dishevelled regulates AIS maintenance.

**Fig. 4.** Axonal excitability is reduced in shDvl knockdown neurons.

**Fig. S7.** Dvl depletion lowers the current threshold for repetitive firing without altering maximal action potential frequency.

**Fig. 5.** Dvl2 recruitment to MT is MTCL1 dependent.

**Fig. S8.** MTCL1 MT-binding sites are required for Dvl2 recruitment to MT

**Fig. 6.** Dvl2 recruitment to MT is PDZ dependent.

**Fig. 7.** PDZ-specific inhibition of Dvl2 impairs condensate formation

**Fig. S9.** Xph334 and Xph335 do not recognize other PDZ-domain proteins like Scribble, PSD93 and Shank3

**Fig. 8.** PDZ-specific inhibition of Dvl2 impairs AIS formation

### Figure S1

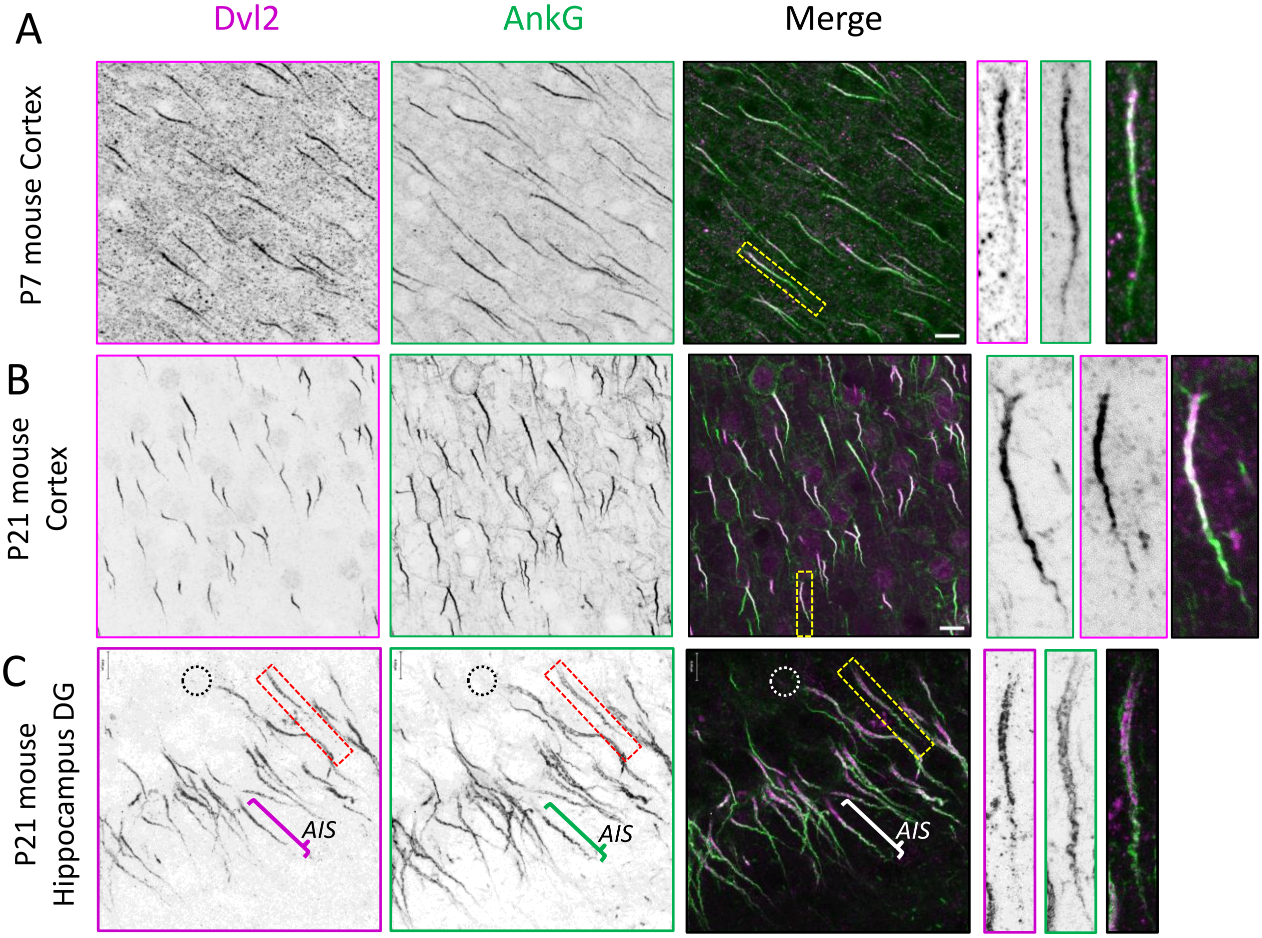

A
Dvl2
AnkG
Merge
P7 mouse Cortex
B
P21 mouse Cortex
C
P21 mouse Hippocampus DG
AIS
AIS
AIS

**Figure S1. Dvl2 is enriched at the AIS in the mouse cortex and DG.**

(**A**) Illustrations of coronal section of brains at P7 immunolabeled for Dvl2 (magenta) and AnkG (green) in the mouse cortex. Dvl2 is enriched in the proximal part of the AIS (bracket, see high magnifications).

(**B-C**) Illustrations of coronal section of brains at P21 immunolabeled for Dvl2 (magenta) and AnkG (green) in the mouse cortex (A) and mouse hippocampus of the DG (B). Dvl2 is enriched in the proximal part of the AIS (bracket, see high magnifications). Dotted circles outline the cell body and bracket the AIS.

Scale bars: 10 µm in A to E.

### Figure S2

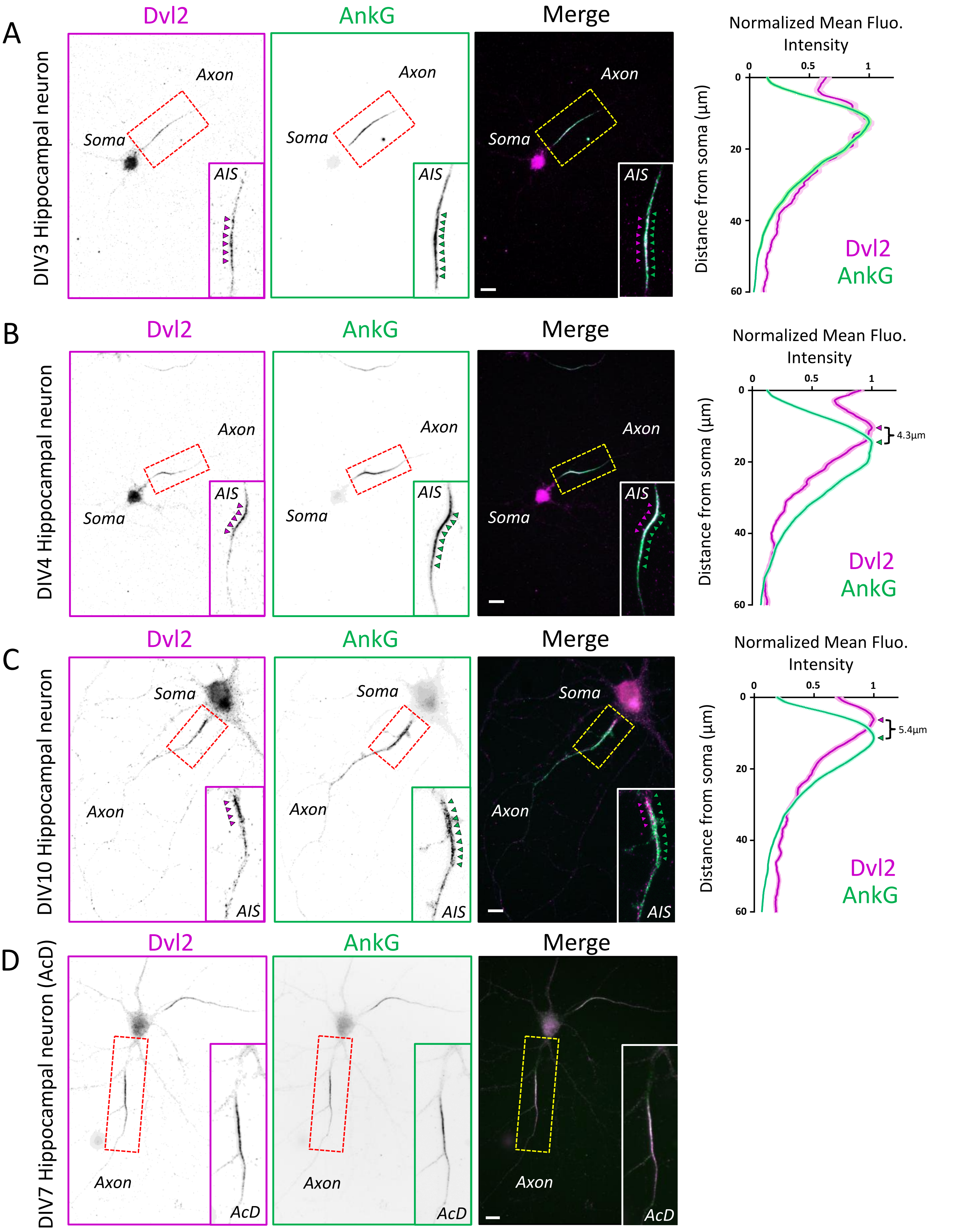
A
B
C
D
Dvl2
AnkG
Merge
DIV3 Hippocampal neuron
DIV4 Hippocampal neuron
DIV10 Hippocampal neuron
DIV7 Hippocampal neuron (AcD)
Soma
Axon
AIS
AcD
Normalized Mean Fluo. Intensity
Distance from soma (µm)
0
0.5
1
20
40
60
4.3µm
5.4µm

**Figure S2. Dvl2 localizes proximally to AnkG, including in axon-carrying dendrite (AcD) neurons in vitro.** (A-C) DIV3 (A), DIV4 (B) and DIV10 (C) cultured hippocampal neurons *in vitro* immunolabeled for Dvl2 (magenta), AnkG (green). Linescans of normalized fluorescence intensity at the AIS of Dvl2 (magenta) and AnkG (green) at different timepoints indicate that Dvl2 is getting enriched at the proximal part of the AIS (see inset, magenta arrowheads) and colocalizes partially with AnkG (bracket and see inset, green arrowheads) from DIV4. Values are normalized to the maximum intensity of each marker and displayed with ± SEM. (**D**) DIV7 hippocampal axon-carrying dendrite (AcD) neurons *in vitro* immunolabeled for Dvl2 (magenta), AnkG (green).
Scale bars : 20 µm in A, 10 µm in B and C.

### Figure S3

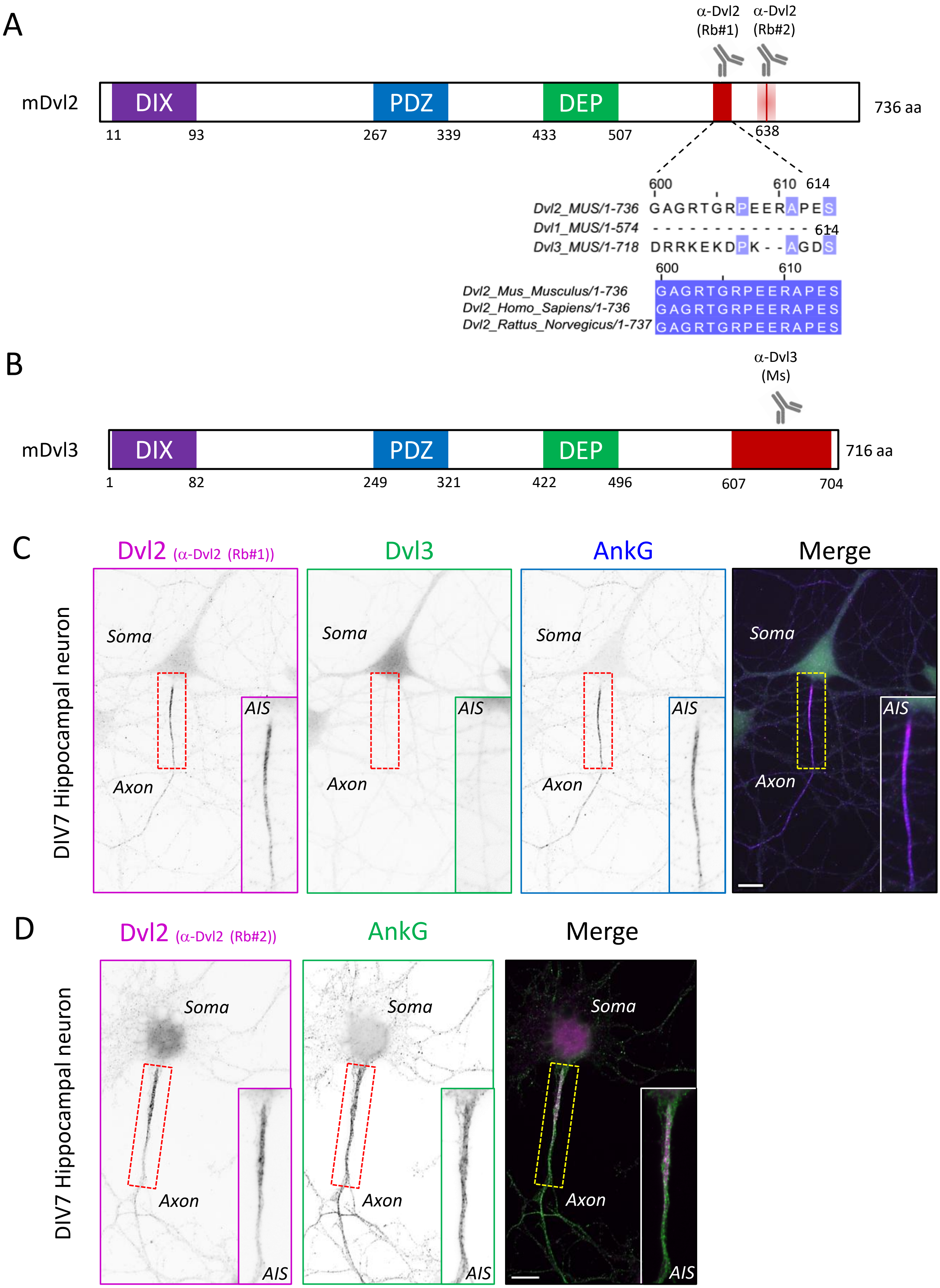
A
α-Dvl2 (Rb#1)
α-Dvl2 (Rb#2)
mDvl2
DIX
PDZ
DEP
736 aa
11
93
267
339
433
507
638
600
610
614
Dvl2_MUS/1-736 GAGRTGRPEERAPES
Dvl1_MUS/1-574 - - - - - - - - - - - - - - 614
Dvl3_MUS/1-718 DRRKEKDPK - - AGDS
600
610
Dvl2_Mus_Musculus/1-736 GAGRTGRPEERAPES
Dvl2_Homo_Sapiens/1-736 GAGRTGRPEERAPES
Dvl2_Rattus_Norvegicus/1-737 GAGRTGRPEERAPES
B
α-Dvl3 (Ms)
mDvl3
DIX
PDZ
DEP
716 aa
1
82
249
321
422
496
607
704
C
Dvl2 (α-Dvl2 (Rb#1))
Dvl3
AnkG
Merge
DIV7 Hippocampal neuron
Soma
Axon
AIS
D
Dvl2 (α-Dvl2 (Rb#2))
AnkG
Merge
DIV7 Hippocampal neuron
Soma
Axon
AIS

**Fig. S3. Characterization of Dvl2 and Dvl3 antibodies in neurons.** (**A**) Schematic of the mouse full-length Dvl2 protein with the different domains and the corresponding amino acid numbers. Dvl2 antibody (Rb #1) recognizes the sequence between 600 and 614aa (red). The specificity of the Dvl2 antibody is explained by the differences in the amino acids sequences of the 3 paralogues. In contrast, the homology in sequence between Mus Musculus, Homo Sapiens and Rattus norvegicus for Dvl2 explains the ability of the antibody to be used in different species. Dvl2 antibody (Rb #2) recognizes the sequence around position 638 (red). (**B**) Schematic of the mouse full-length Dvl3 protein with the different domains and the corresponding amino acid numbers. Dvl3 antibody recognizes the sequence between 607 and 704aa (red). (**C**) DIV7 hippocampal neurons in vitro immunolabeled for Dvl2 (magenta), Dvl3 (green) and AnkG (blue). Dvl2 staining partially overlaps with AnkG immunolabeling at AIS but there is no Dvl3 labelling. (**D**) DIV7 hippocampal neurons in vitro immunolabeled for Dvl2 (magenta) with the second antibody (Rb #2) and AnkG (green). Dvl2 staining partially overlaps with AnkG immunolabeling at AIS. Scale bar : 10 µm in C and D.

### Figure S4

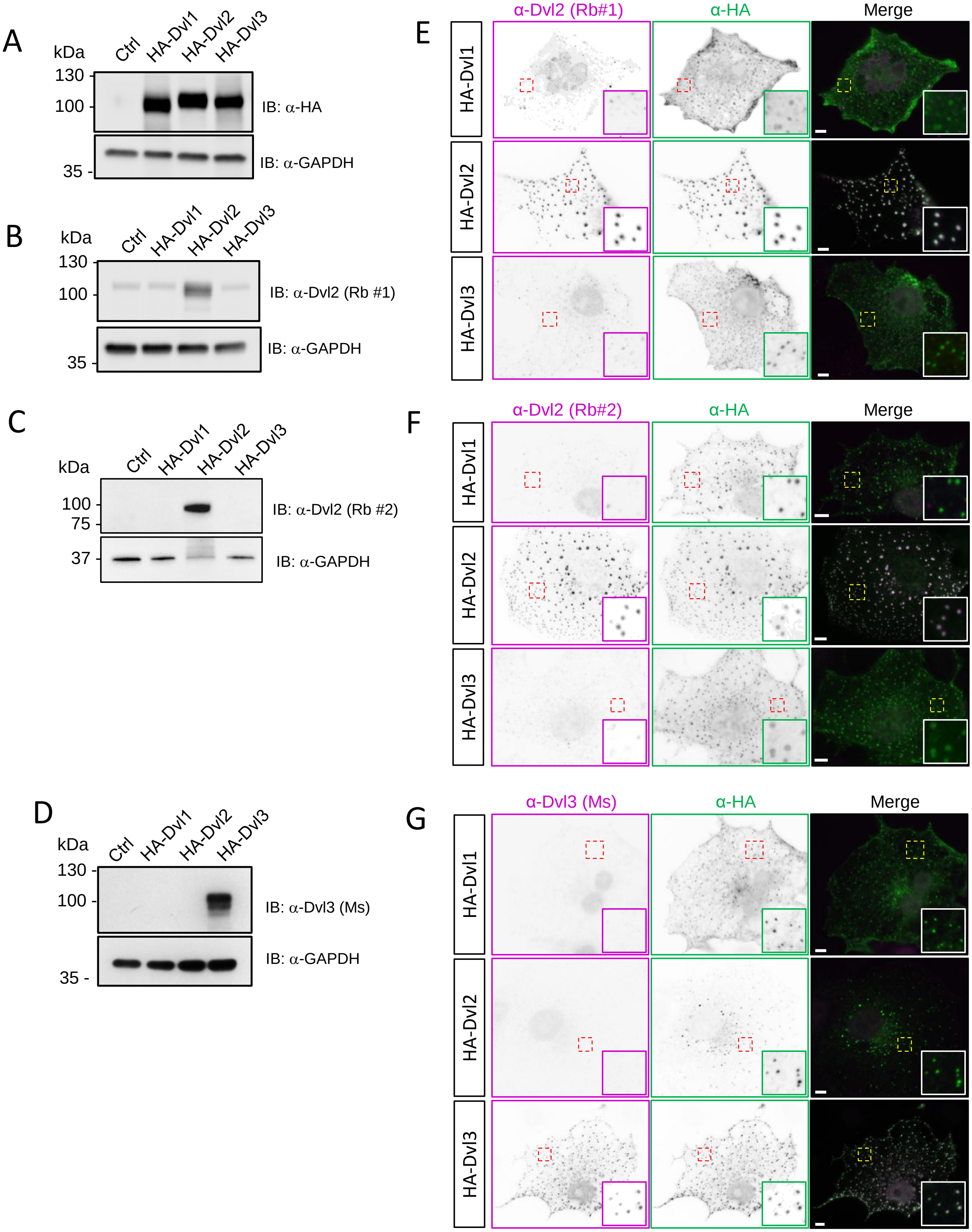
A
kDa
Ctrl
HA-Dvl1
HA-Dvl2
HA-Dvl3
130
100
35
IB: α-HA
IB: α-GAPDH
B
kDa
130
100
35
IB: α-Dvl2 (Rb #1)
IB: α-GAPDH
C
kDa
100
75
37
IB: α-Dvl2 (Rb #2)
IB: α-GAPDH
D
kDa
130
100
35
IB: α-Dvl3 (Ms)
IB: α-GAPDH
E
α-Dvl2 (Rb#1)
α-HA
Merge
F
α-Dvl2 (Rb#2)
α-HA
Merge
G
α-Dvl3 (Ms)
α-HA
Merge

**Fig. S4. Characterization of Dvl2 and Dvl3 antibodies in Cos7 cells**

(**A**) Anti-HA blot confirms expression of HA-Dvl1, HA-Dvl2, and HA-Dvl3 in the corresponding transfected cells, with no signal in the WT cos-7 cells. Anti-GAPDH was used as loading control. (**B**) Anti-Dvl2 (Rb #1) detects a strong band at approximately 100 kDa, consistent with the migration of Dvl2 in western blot, which runs above its predicted molecular weight. Faint bands appear in lines 1, 2 and 3 corresponding to endogenous Dvl2 already present in Cos-7 cells. Anti-GAPDH was used as loading control. (**C**) Anti-Dvl2 (Rb #2) detects a strong band at approximately 100 kDa, consistent with the migration of Dvl2 in western blot, which runs above its predicted molecular weight. Anti-GAPDH was used as loading control. (**D**) Anti-Dvl3 detects a strong band at approximately 100 kDa, consistent with Dvl3 molecular weight when overexpressed in Cos-7 cells. Anti-GAPDH was used as loading control. (**E-G**) Cos7 cells transfected with HA-Dvl1, HA-Dvl2 or HA-Dvl3 immunolabeled with anti-Dvl2 (Rb #1) (E), anti-Dvl2 (Rb #2) (F) or anti-Dvl3 (G) in magenta and HA in green. Anti-Dvl2 (Rb #1) and anti-Dvl2 (Rb #2) labeling colocalize only with HA-Dvl2 respectively in (E) and (F) and anti-Dvl3 labeling colocalize only with HA-Dvl3 (G). Scale bar : 5 µm in E to G.

### Figure S5

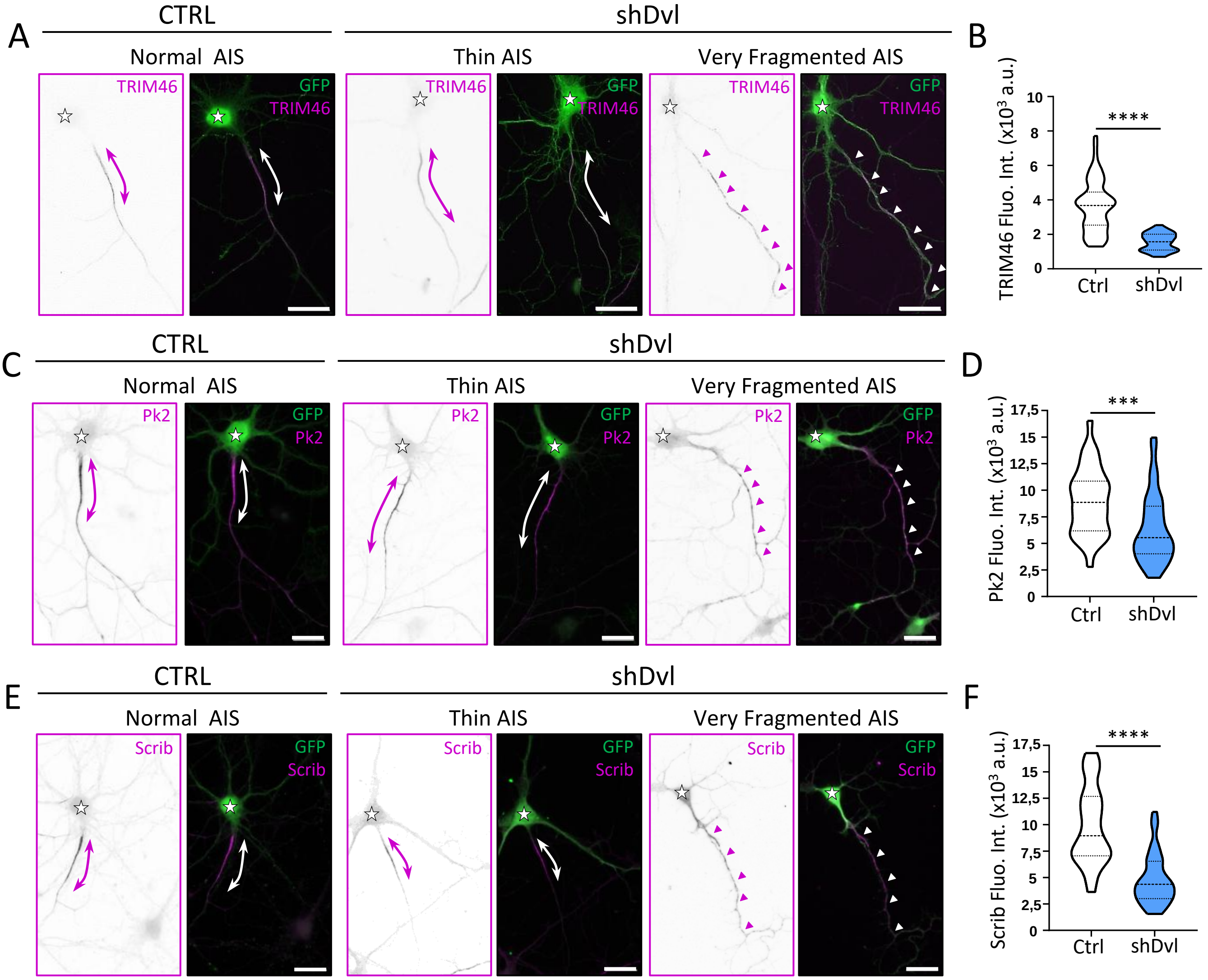

A
CTRL
shDvl
Normal AIS
Thin AIS
Very Fragmented AIS
TRIM46
GFP
TRIM46
B
TRIM46 Fluo. Int. (x10³ a.u.)
****
Ctrl
shDvl
C
Pk2
D
Pk2 Fluo. Int. (x10³ a.u.)
***
E
Scrib
F
Scrib Fluo. Int. (x10³ a.u.)
****

**Fig S5: Dishevelled regulates AIS formation (additional markers).** (A, B, C) Illustration of DIV7 neurons transfected with a control or shDvl plasmid and labelled for TRIM46 (A), Pk2 (B) or Scribble (Scrib) (C) (magenta) and GFP (green). Stars show the cell body and double arrows or arrowhead the AIS. Dvl-induced AIS fragmentation is illustrated as small, interspersed TRIM46+, Pk2+ or Scribble+ clusters shown by the arrowheads. (D, E, F) Quantifications of fluorescence intensity levels of TRIM46 (D), Pk2 (E) or Scribble (F) in shDvl neurons (excluding no AIS phenotypes) show downregulations in protein levels (****$p<0.0001$ or ***$p<0.001$ from Mann-Whitney's test. TRIM46 (n=61 neurons CTRL, n=50 shDvl), Pk2 (n=33 neurons CTRL, n=40 shDvl) and Scrib (n=35 neurons CTRL, n=39 shDvl), from at least 3 independent experiments). Scalebars : 20 µm in (A, B, C).

### Figure S6

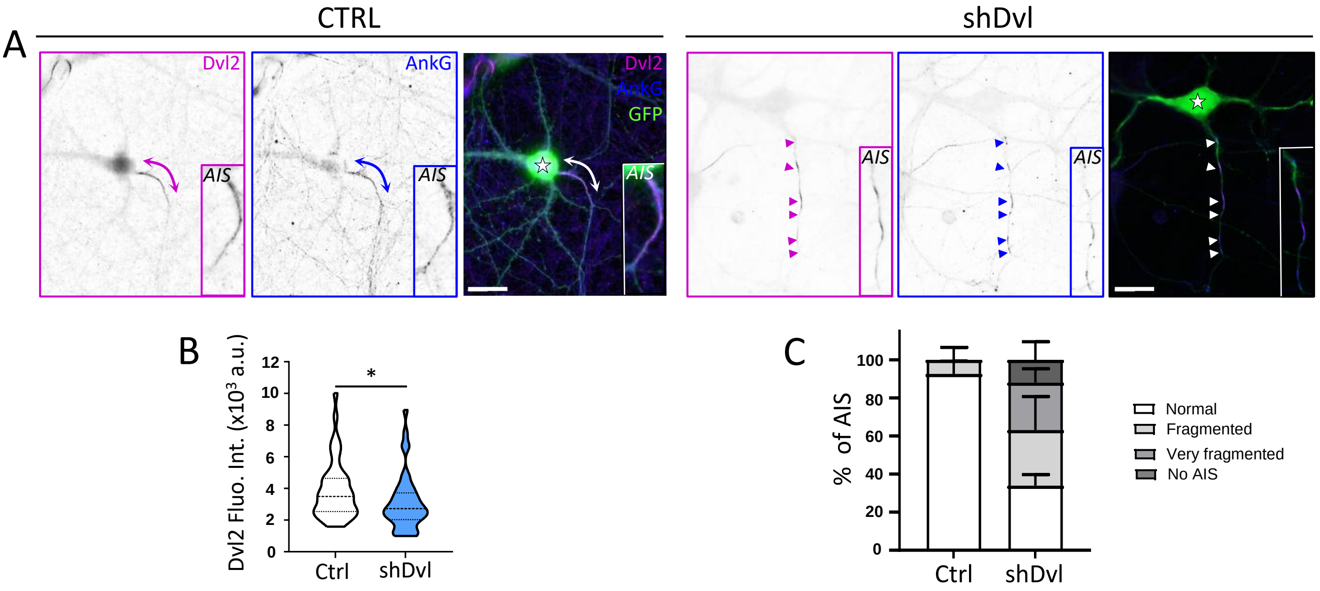
A
CTRL
shDvl
Dvl2
AnkG
GFP
AIS
B
Dvl2 Fluo. Int. (x10³ a.u.)
12
10
8
6
4
2
0
*
Ctrl
shDvl
C
% of AIS
100
80
60
40
20
0
Ctrl
shDvl
Normal
Fragmented
Very fragmented
No AIS

**Fig S6: Dishevelled regulates AIS maintenance.** (A) Illustration of DIV17 neurons transfected at DIV13 with a control or shDvl plasmid and labelled for Dvl2 (magenta) and AnkG (blue). Stars show the cell body and double arrows the AIS. The shDvl-induced AIS fragmentation is illustrated as small, interspersed AnkG+ clusters as in (arrowheads) and Dvl2 labelling is decreased. (B) Quantification of fluorescence intensity level of Dvl2 in shDvl neurons at DIV17 (excluding no AIS phenotypes) show downregulation Dvl2 levels. (C) Percentages neurons per phenotype at DIV17 described as normal AIS, fragmented, very fragmented and loss of AIS with shDvl. Means ± SEM are shown. Scale bars : 20 µm in A.

### Figure S7

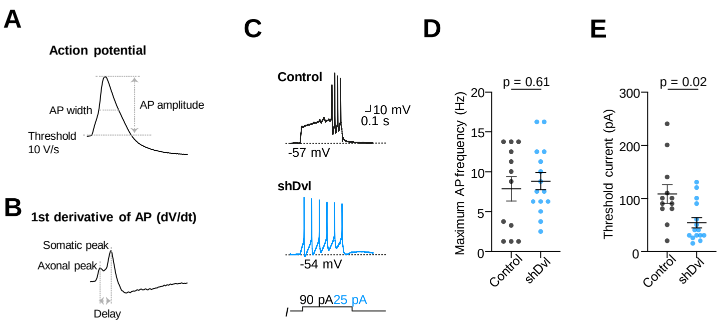

A
Action potential
AP width
AP amplitude
Threshold
10 V/s
B
1st derivative of AP (dV/dt)
Somatic peak
Axonal peak
Delay
C
Control
10 mV
0.1 s
-57 mV
shDvl
-54 mV
90 pA
25 pA
I
D
p = 0.61
Maximum AP frequency (Hz)
20
15
10
5
0
Control
shDvl
E
p = 0.02
Threshold current (pA)
300
200
100
0
Control
shDvl

**Figure S7: Dvl depletion lowers the current threshold for repetitive firing without altering maximal action potential frequency.** (A) Action potential analysis schematic for the most important parameters. (B) 1st derivative (dV/dt) of the action potential recorded at the soma showing the two components corresponding to the backpropagating AIS peak and the somatic peak. (C) Example recordings during long positive current injections show the generation of several action potentials in control and shDvl neurons. Several action potentials are elicited in each cell, however, firing patters differ probably as the neuronal populations were mixed hippocampal cells. (D) Population data of maximum AP frequency shows no difference between groups. (E) The shDvl neurons require much lower current injections to generate several action potentials consistent with the measured higher input resistance.

### Figure S8

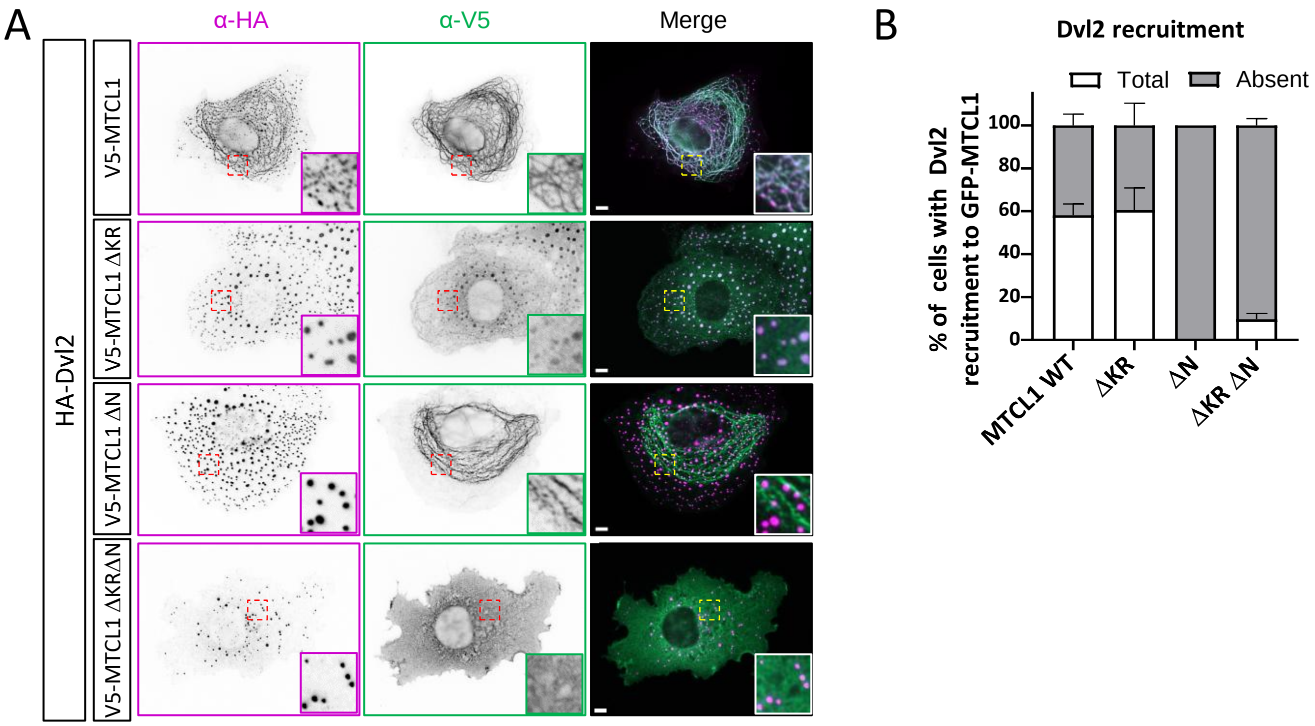
A
α-HA
α-V5
Merge
HA-Dvl2
V5-MTCL1
V5-MTCL1 ΔKR
V5-MTCL1 ΔN
V5-MTCL1 ΔKRΔN
B
Dvl2 recruitment
Total
Absent
% of cells with Dvl2 recruitment to GFP-MTCL1
0
20
40
60
80
100
MTCL1 WT
ΔKR
ΔN
ΔKR ΔN

**Fig. S8 MTCL1 MT-binding sites are required for Dvl2 recruitment to MT. (A)** COS7 cells transfected for 24 hrs with HA-Dvl2 either with V5-MTCL1 or MTCL1 deletion mutants for the KR, N, and both KR and N domains. **(B)** Quantification of the specific HA-Dvl2 recruitment to MTCL1+ MTs. HA-Dvl2 colocalizes with V5-MTCL1DKR which is less efficiently localized on MTs, but HA-Dvl2 does not colocalize with V5-MTCL1DN or V5-MTCL1DKRDN. Scale bar : 5 µm in A

### Figure S9

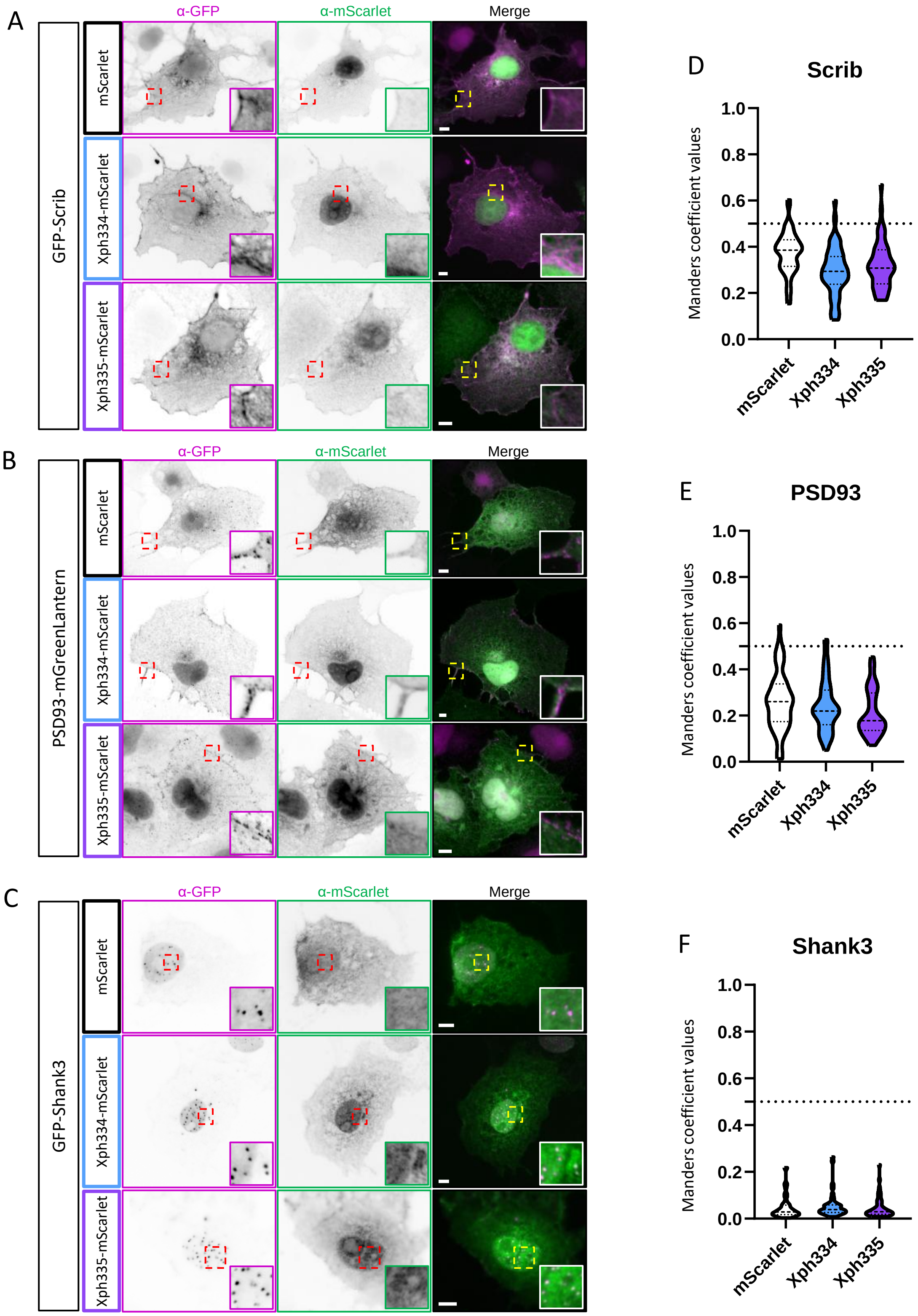

A
B
C
D
E
F
α-GFP
α-mScarlet
Merge
GFP-Scrib
PSD93-mGreenLantern
GFP-Shank3
mScarlet
Xph334-mScarlet
Xph335-mScarlet
Scrib
PSD93
Shank3
Manders coefficient values
1.0
0.8
0.6
0.4
0.2
0.0
mScarlet
Xph334
Xph335

**Fig. S9. Xph334 and Xph335 do not recognize other PDZ-domain proteins like Scribble, PSD93 and Shank3.** COS7 cells were co-transfected with GFP-Scrib (A), PSD93-mGL (B), or GFP-Shank3 (C) (magenta) and either mScarlet, Xph334 or Xph335 (green). Merged images show no overlap between magenta and green signals, confirming absence of colocalization. Quantification of colocalization using Manders' coefficient confirms the lack of colocalization (D–F). Scale bar : 5 µm in A to C
